# Evolutionary investigations of the biosynthetic diversity in the skin microbiome using *lsa*BGC

**DOI:** 10.1101/2022.04.20.488953

**Authors:** Rauf Salamzade, J.Z. Alex Cheong, Shelby Sandstrom, Mary Hannah Swaney, Reed M. Stubbendieck, Nicole Lane Starr, Cameron R. Currie, Anne Marie Singh, Lindsay R. Kalan

**Affiliations:** Department of Medical Microbiology and Immunology, School of Medicine and Public Health, University of Wisconsin, Madison, WI, USA; Department of Bacteriology, University of Wisconsin—Madison, Madison, Wisconsin, USA; Department of Pediatrics, School of Medicine and Public Health, University of Wisconsin, Madison, WI, USA; Department of Medicine, School of Medicine and Public Health, University of Wisconsin, Madison, WI, USA

## Abstract

We developed *lsa*BGC, a bioinformatics suite that introduces several new methods to expand on the available infrastructure for genomic and metagenomic-based comparative and evolutionary investigation of biosynthetic gene clusters (BGCs). Through application of the suite to four genera commonly found in skin microbiomes, we uncover multiple novel findings on the evolution and diversity of their BGCs. We show that the virulence associated carotenoid staphyloxanthin in *Staphylococcus aureus* is ubiquitous across the *Staphylococcus* genus but has largely been lost in the skin-commensal species *Staphylococcus epidermidis*. We further identify thousands of novel single nucleotide variants (SNVs) within BGCs from the *Corynebacterium tuberculostearicum* sp. complex, which we describe here to be a narrow, multi-species clade that features the most prevalent *Corynebacterium* in healthy skin microbiomes. Although novel SNVs were approximately ten times as likely to correspond to synonymous changes when located in the top five percentile of conserved sites, *lsaBGC* identified SNVs which defied this trend and are predicted to underlie amino acid changes within functionally key enzymatic domains. Ultimately, beyond supporting evolutionary investigations, *lsa*BGC provides important functionalities to aid efforts for the discovery or synthesis of natural products.

## Introduction

The secondary metabolome of bacteria has served as a valuable reservoir of natural products with great societal benefit^1^. Historically, many commercialized secondary metabolites, such as antibiotics, have been identified from particular taxonomic groups, such as the genus of *Streptomyces* within Actinomycetota^1–3^. Several studies have explored the secondary metabolome of these metabolically rich microbial taxa to develop a comprehensive understanding of their chemical diversity and unique traits^4–7^. Most of these studies begin by first identifying biosynthetic gene clusters within genomes using the popular antiSMASH software^8^ and then group BGCs into gene cluster families (GCFs) using BiG-SCAPE^9^. This approach, similar to other bioinformatics software for genomics annotation and analysis of BGCs^10–12^, uses a set of key protein domains as their base to group BGCs predicted to encode similar metabolites. However, when working in the context of a single genus or species, it becomes more appropriate to use full protein sequences which enables the use of advanced algorithms for the identification of orthologs and paralogs^13^ to reliably determine evolutionary relationships between BGCs.

Recent studies which explored the vertical descent and conservation of BGCs across the genus of *Salinospora* and the fungal species *Aspergillus flavus* have shown that intra-genus or intra-species evolution can result in chemical and regulatory diversification of the encoded metabolites^6,14^. Thus, improved understanding of the evolutionary emergence of BGCs could be informative towards elucidating the function of individual genes and encoded metabolites^9,15–20^.

Due to difficulties in the cultivation of many bacterial taxa, several recent studies have begun to explore metagenomic datasets to unearth novel secondary metabolites^21–23^. While most of these endeavors have relied on performing metagenomic assembly, recently, two read-based approaches were described to reliably and sensitively search for key BGC domains in metagenomes^22,24^. This highlights a larger trend in which these approaches primarily aim to identify highly novel secondary metabolites and thus prioritize investigation of poorly studied or newly discovered taxa^25^. However, provided that most metabolites have been mined from select genera, mostly within the *Actinomycetia* class^2^, there could be tremendous reward for efforts to advance our understanding of cataloged but uncharacterized BGCs from well-studied taxa. This is especially important because many species within the metabolically rich *Streptomyces* only have limited representative genomes, thus methods to leverage metagenomic datasets and identify homologous instances of rare BGCs can lead to critical insight into their function and ecological distribution^4^.

To aid the advancement of such taxa-specific analyses of BGCs, we introduce the comprehensive software suite *<u>L</u>ineage <u>S</u>pecific <u>A</u>nalysis* of BGCs (*lsa*BGC; https://github.com/Kalan-Lab/lsaBGC), which consists of several programs for comparative genetics of BGCs as well as metagenomic mining. We showcase the utility of *lsa*BGC in this study through application to four major genera which are commonly found in healthy human skin microbiomes. Several secondary metabolites which function as antibiotics^26–28^, virulence factors^29^, or in microbe-host interactions^30,31^ have been identified in bacteria from skin microbiomes, primarily from staphylococci. Human skin is also easily accessible to sampling and most species can be cultivated for follow up studies. Using *lsa*BGC, we reveal new ecological and evolutionary insights for both well characterized and unknown predicted metabolites, including signatures of inter-species transfer for the virulence-associated staphylococcal carotenoid staphyloxanthin and a highly conserved predicted siderophore found encoded by divergent *Corynebacterium* species. In addition, through mining skin metagenomic datasets, we find novel coding variants within the functionally important ketide synthase (KS) domain of the polyketide synthase (PKS) responsible for mycolic acid biosynthesis, a defining cell wall component of *Corynebacterium* species implicated in immune recognition.

## New Approaches

Comprehensive studies of the BGCs for specific taxa have become commonplace in the last five years owing to advances and standardization in methods to cluster BGCs into GCFs^9,12^. Deeper investigations into individual GCFs, however, remains a largely manual process. Here we introduce *lsa*BGC which simplifies such investigations with easy-to-use workflows that require minimal knowledge of the command-line. *lsa*BGC features a comprehensive set of functionalities to cluster BGCs into GCFs, compute evolutionary statistics, and perform metagenomic mining. Finalized reports include a spreadsheet which lists all genes found in GCFs together with annotation information, conservation, and evolutionary statistics to allow quick assessment of a taxa’s biosynthetic potential. Additionally, visualizations are automatically generated including a view of GCFs across a species phylogeny, which is commonly reported in recent studies^6,7,14^ but for which no software currently exists to automate the process. Finally, through utilization of the metagenomic mining functionality in *lsa*BGC, users can obtain a report on whether metagenomes contain novel alleles of known BGCs found in available assemblies.

## Results

### *lsa*BGC allows for sensitive and systematic identification of BGC homolog groups

The *lsa*BGC suite consists of eight core programs, four workflows, and several additional scripts for analyses (Fig. 1, S1; Supplementary Text). These programs allow for clustering BGCs into GCFs, refining boundaries of BGCs, and sensitive, yet specific, rapid detection of GCF instances in draft-quality assemblies. The suite additionally includes dedicated tools for GCF visualization, evolutionary and population genetic analysis of individual genes within GCFs, and base-resolution identification of novel SNVs in complex metagenomes.

**Figure 1:**
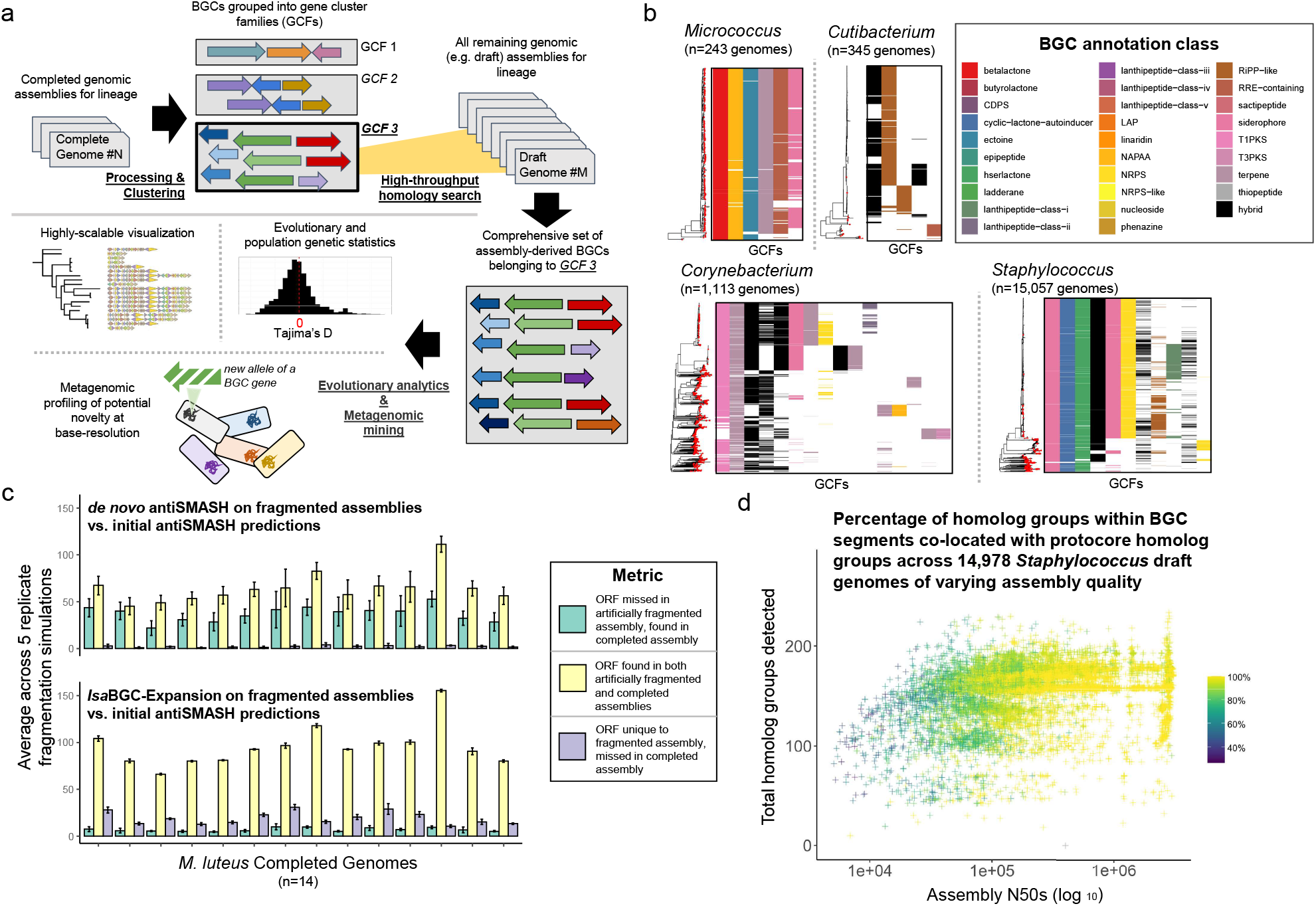
*lsa*BGC offers an efficient and sensitive means for GCF homology detection. **a**) A schematic overview of the full *lsa*BGC suite. **b**) Neighbor-joining trees were constructed for each of the four genera based on pairwise MASH ANI estimates between genomes and heatmaps showcase the presence of GCFs found in > 5% of genomes. Red dots on neighbor-joining trees signify complete or chromosomal genomes used for initial identification of BGCs. Heatmap cell color indicates the BGC class as predicted by antiSMASH. **c**) Results from benchmarking *lsa*BGC-AutoExpansion sensitivity on 14 completed genomes for the species of *M. luteus*. AntiSMASH and *lsa*BGC-AutoExpansion were run on artificially fragmented genomes and resulting BGC predictions were compared to initial BGCs identified on the unfragmented, complete genomes. **d**) The relationship between the assembly quality measured by N50 and the total number of homolog groups detected across 14,978 *Staphylococcus* draft genomes is shown. The coloring corresponds to the percentage of BGC homolog groups co-located in a segment with a homolog group commonly found in protocore regions of BGC predictions by antiSMASH run on completed genomes.

To demonstrate the capabilities of the *lsa*BGC suite we applied it to four genera which are stable and integral members of the human skin microbiome^32–36^. These included three genera in the Actinomycetota (formerly Actinobacteria) phylum: *Corynebacterium*,*Cutibacterium*, and *Micrococcus* as well as the Bacillota (formerly Firmicutes) genus *Staphylococcus*. *lsa*BGC was also independently applied to the species or species-complex from each genus most commonly represented in skin microbiomes. These were the *C. tuberculostearicum* species-complex, *Cutibacterium acnes, Micrococcus luteus*, and *S. epidermidis* (Table S1).

We used *lsa*BGC-AutoProcess to run gene-calling, BGC annotation, and *de novo* homolog group delineation with complete or chromosomally complete genomes for each genus or species. We then applied *lsa*BGC-Cluster to group homologous BGCs into GCFs and systematically searched for instances of GCFs using *lsa*BGC-AutoExpansion (Fig. 1b; Table S2, S3; Supplementary Text). To assess the validity and sensitivity of our approach, two benchmarking tests using BGCs identified in *M. luteus* were performed (Fig. 1c, S3; Supplementary Text) showing that the *lsa*BGC-AutoExpansion framework leads to greater sensitivity compared to running *de novo* antiSMASH on fragmented draft-quality assemblies. This increased sensitivity was due to more robust, yet reliable, detection of auxiliary genes located on the edges of scaffolds not containing core domains used by antiSMASH for BGC detection. In addition, *lsa*BGC-AutoExpansion is highly efficient and allowed us to search for 63 GCFs identified from high-quality *Staphylococcus* genomes in 14,978 draft-quality genomes in approximately 14 hours using 40 cores and less than 150 Gb of memory. We found a positive association between detection of complete BGC instances and assembly quality (Fig. 1d; Supplementary Text).

### *lsa*BGC enables reliable comparative genomics to explore evolutionary trends of BGCs

The increased sensitivity of *lsa*BGC to uniformly detect auxiliary components of BGCs in each genome enabled us to perform reliable comparative genetic analyses across GCF instances. We focused on two of the four genera with the greatest breadth of diversity, *Corynebacterium* and *Staphylococcus*, and selected a subset of 456 and 229 dereplicated, representative genomes from each genera, respectively (Table S1). To better understand which species in these genera were most relevant to skin health, we first profiled how often distinct species were found among healthy skin microbiomes using a large, recently generated metagenomics dataset^37,38^ (Table S4). Skin-associated species (found in more than 10 of 270 metagenomes) were phylogenetically clustered for both genera, within the *S. epidermidis/aureus* clade (Fig. S4a) and within the *C. tuberculostearicum* species complex (Fig. S4b; Supplementary Text).

Association of individual homolog groups from these GCFs revealed several were enriched within skin-associated staphylococci, but only two were enriched for skin-associated corynebacteria, both encoding hypothetical proteins (Table S5). Nearly half of the homolog groups enriched for skin-associated staphylococci (5 of 12; 41.7%) were related to GCF-10, predicted to encode for an NRP synthetase corresponding to pyrazinone biosynthesis, molecules reported to regulate virulence in *S. aureus*^29,39^. Staphylococcal pyrazinones were previously identified as being a self-contained BGC consisting of two genes, *pznA* and *pznB*, specific to species commonly associated with human skin, including *S. aureus*, *S. epidermidis*, and *Staphylococcus capitis*^29,39^. Using *lsa*BGC, we discover novel variants of pyrazinones, which encapsulate seven distinct GCFs (Fig. 2a), that belong to species not commonly found on human skin, such as *Staphylococcus felis* (GCF-56), the dominant species of *Staphylococcus* on cat skin. While speciation had resulted in orthologous copies of *pznA* and *pznB* being highly diverged in sequence, it is surprising to find that the surrounding genomic context of the two genes is highly variable for all seven GCFs (Fig. 2b), suggesting their ancestral mobilization or relocation within or between genomes.

**Figure 2:**
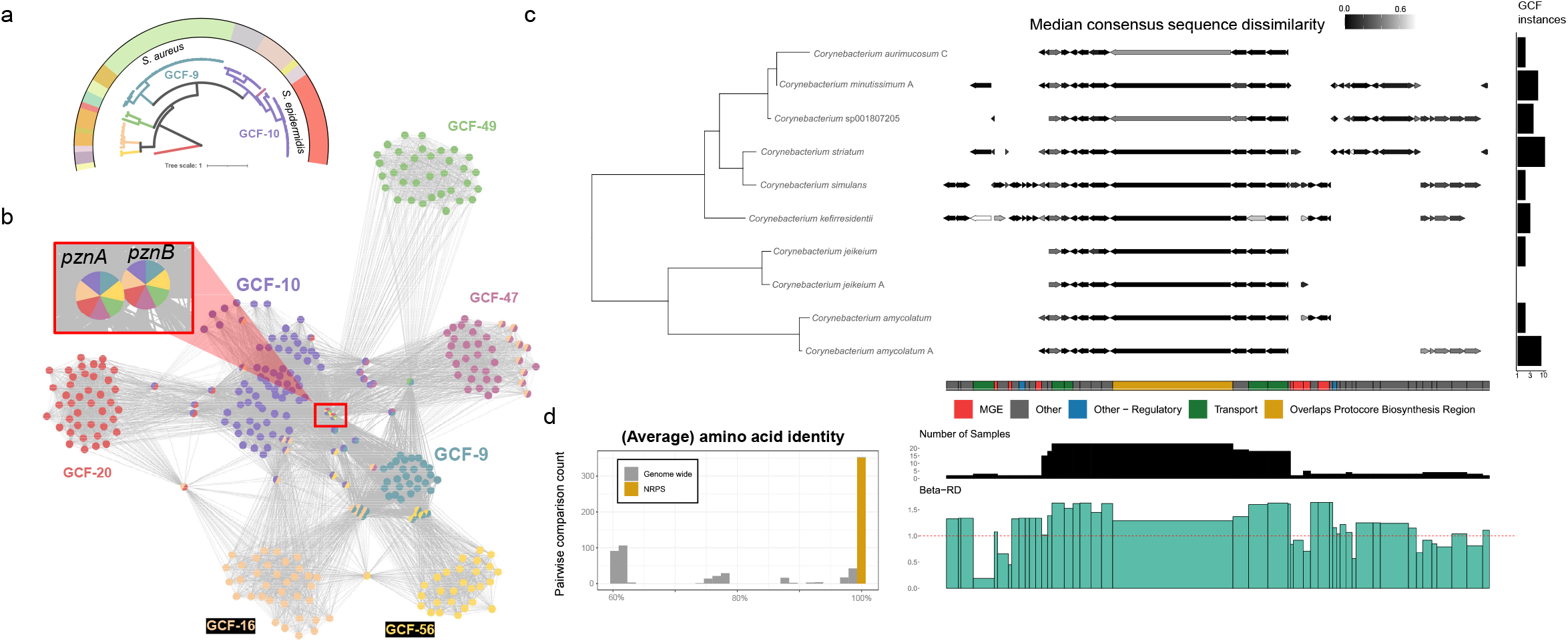
*lsa*BGC reveals new insight about characterized and unstudied BGCs from skin-associated species. **a**) A maximum-likelihood phylogeny was constructed from PznA and PznB showing the relationship between GCFs encoding for pyrazinones. Branch color corresponds to different GCFs and the outer circular color denotes the species classification for the genome from which the sequences were gathered.**b**) Homolog groups across the seven pyrazinone encoding GCFs are depicted as a network where edges indicate that two homolog groups co-occur in the same GCF. Homolog group color of nodes depict distinct GCFs.**c**) Phylogeny constructed from ribosomal proteins showing the conservation of *Corynebacterium* GCF-50 across the diverse species encoding the NRPS using results from *lsa*BGC-PopGene analysis. The gene schematics show the consensus gene order for the GCF. Gene lengths correspond to the median length across all BGC instances and only genes found in two or more genomes are shown. For each species, the gene is colored based on the median similarity of gene sequences from the species to the consensus sequence across all species. The barplot displays the total number of GCF instances identified per species across the full set of *Corynebacterium* genomes. Broad annotations for homolog groups are shown as a color strip. Below it are bar plots corresponding to the conservation of homolog groups across samples and their median Beta-RD. **d**) A histogram of the amino acid identities between GCF-50 NRP synthase instances which are complete (excluding partial instances due to assembly fragmentation included in panel **c**) is compared to the average amino acid identities (AAI) between genomes they were gathered from.

Application of *lsa*BGC-Divergence followed by Bayesian shrinkage analysis was then used to systematically assess whether pyrazinones or other *Staphylococcus* and *Corynebacterium* GCFs exhibited indications of HGT between species, as suggested by high Beta-RD values (Fig. S4cd; Table S6; Supplementary Text). Despite the small size of pyrazinones, we do not find support for the frequent mobilization of any of the seven GCFs encoding them across staphylococcal species. However, high Beta-RD posterior ranges were observed for two non-ribosomal peptide synthetase (NRPS) associated GCFs from *Corynebacterium*. One of these GCFs, predicted to encode for an NRPS-dependent siderophore^40^, was found in several species commonly isolated from skin^37^, including *C. kefirresidentii*, from the *C. tuberculostearicum* sp. complex. Using results from *lsa*BGC-PopGene for this GCF, which includes determination of a consensus gene order across BGC instances, we find that the NRPS is highly conserved in sequence across species and found flanked by mobile genetic elements (MGEs), as previously reported in *Corynebacterium jeikeium*^40^ (Fig. 2cd). The taxonomic breadth and high sequence conservation of this GCF suggest that it has undergone recent HGT between distantly related species.

We next used *lsa*BGC-PopGene to infer selective pressures acting on homolog groups within GCFs through calculating inter- and intra-species Tajima’s D using select representative genomes, obtained through genomic dereplication, for each genus, including *Cutibacterium* and *Micrococcus* (Table S1, S7; Supplementary Text). While *lsa*BGC-PopGene is highly scalable, it is important to perform uniform dereplication of genomes to accurately infer species-wide signatures of selection without being biased by highly represented strains in genomic datasets. Further, only homolog groups which were found as a single copy within the GCF context and observed in four of more genomes were considered. We found that the median intra-species Tajima’s D was centered at a value of roughly 0, in accordance with expectations of the null model (Fig. S5a). Also in alignment with expectations, we observed high values of inter-species Tajima’s D for homolog groups in GCFs found in multiple species (Fig. 5b). A focused intra-species analysis within *S. aureus* revealed that a hybrid terpene/T3PKS GCF (GCF-3) had the lowest Tajima’s D values and seven homolog groups with values below −2.0, suggestive they are either highly conserved or, potentially, under sweeping selection (Fig. 5c). The terpene component of this hybrid GCF encodes for staphyloxanthin carotenoid production^41^ and manual inspection determined that the T3PKS prediction actually corresponds to hydroxymethylglutaryl-CoA (HMG-CoA) synthase, which is involved in the mevalonate pathway for isoprenoid synthesis and is homologous to KS domains of polyketide synthases^42^. Further, the regions in between and flanking the staphyloxanthin encoding operon, *crt*, and HMG-CoA synthase include several genes important to staphylococcal virulence, some of which were found in multiple copies in certain species (Fig. S6ab). Because this GCF was predicted to be one of the four GCFs ancestrally acquired by the *S. epidermidis/aureus* clade, the five gene *crt* operon encoded by it was found in most species belonging to the clade. Surprisingly however, *crt* genes were missing in more than 95% of the *S. epidermidis* genomes, the most prevalent staphylococcal species on skin^34,43^, likely due to gene loss (Fig. 4a, S6b).

### The virulence-associated carotenoid staphyloxanthin is ubiquitous across the *Staphylococcus* genus

Although staphyloxanthin is well characterized as an *S. aureus* virulence factor^44^, we were unable to identify a report of its distribution across the *Staphylococcus* genus, with only a single study, to our knowledge, suggesting its production in another species^45^. *lsa*BGC analysis revealed a total of five staphylococcal GCFs encoding for staphyloxanthin. Following GCF-3, the second most prevalent GCF encoding staphyloxanthin, GCF-6, had a high Beta-RD distribution (Fig. S4c) and was widely distributed across divergent species in the genus (Fig. 3a), as well as *Mammaliicoccus*, the closest phylogenetic neighbors of staphylococci^46^. Constructing a phylogeny based on a concatenated alignment of *crtM* and *crtN* revealed the appropriate delineation of GCF-3 and GCF-6 despite their co-occurrence in the genomes of particular species, such as *Staphylococcus warneri* (Fig. 3b, S7a). Additional analyses showed that GCF-6 has mobilized across species and can be found on plasmids (Fig. 3bcd).

**Figure 3:**
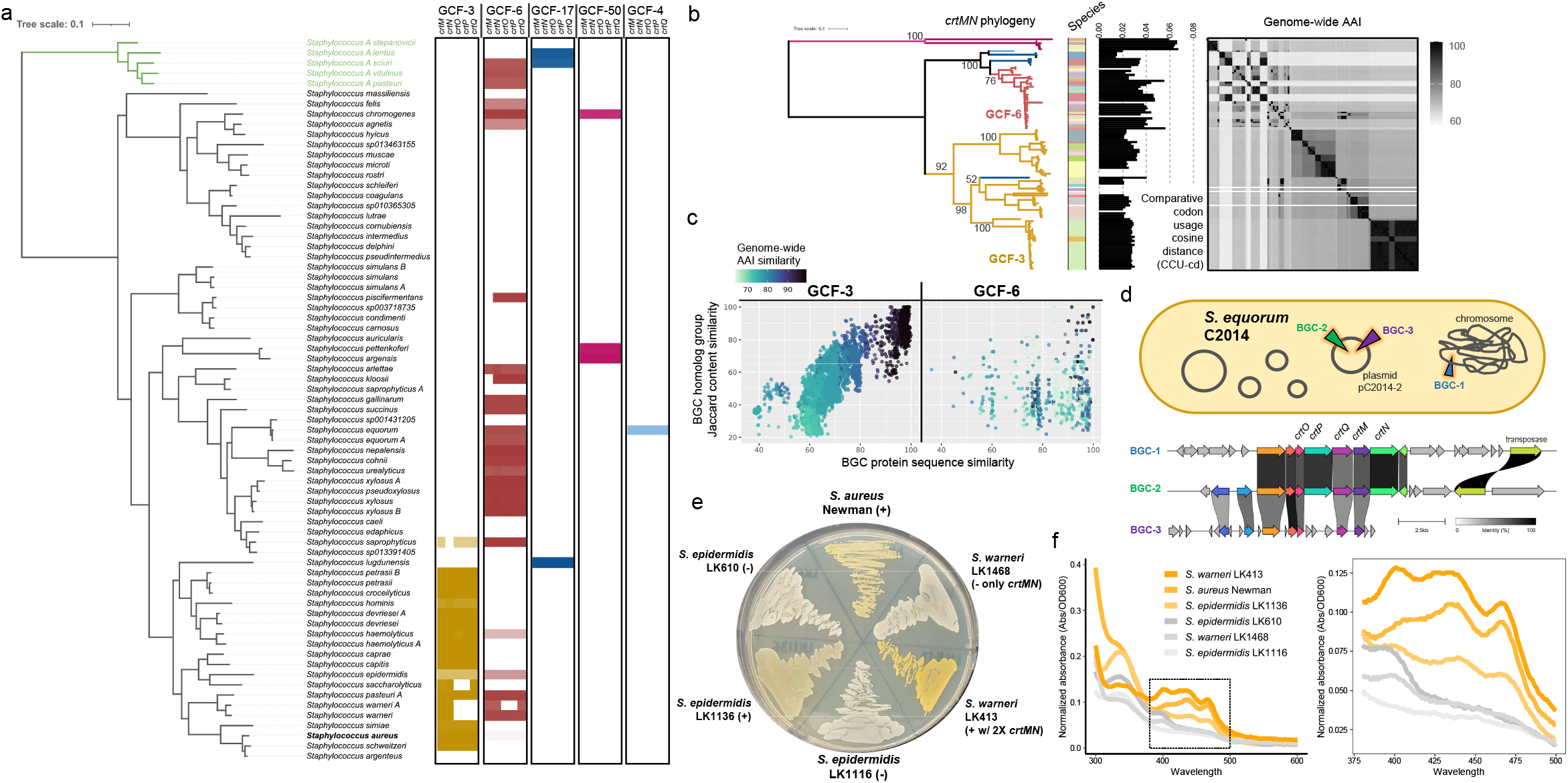
Staphyloxanthin is ubiquitous across *Staphylococcus* and found on plasmids. **a**) A maximum-likelihood phylogeny of the full *Staphylococcus* genus was constructed from ribosomal protein encoding genes. The clade in green represents the newly recognized genus of *Mammaliicoccus*. Heatmaps showcase the presence of the five *crt* genes for each species for five separate GCFs containing the *crt* operon. The shading indicates the proportion of a species found to carry homolog groups within a particular GCF context in log_10_ scale. **b**) A maximum-likelihood phylogeny was built from CrtM and CrtN protein alignments. Branch colors represent the GCF classification of the CrtMN sequences and bootstrap values are shown for key nodes where GCFs partition. The species the CrtMN sequence was extracted from is shown as a color strip followed by a bar plot depicting the comparative codon usage cosine distance (CCU-cd). CCU-cd was only calculated for the full five gene *crt* operon and represents the codon frequency dissimilarity with the codon frequency of the background genome. The heatmap showcases the genomewide average amino acid identity (AAI) between pairs of genomes from which CrtMN sequences were gathered. **c**) AAI and shared homolog group content were calculated between pairs of GCF instances from different genomes for GCF-3 and GCF-6 individually. The coloring represents genome-wide AAI similarity. **d**) A schematic of the completed genome for *S. equorum* C2014, which was predicted to feature GCF-6 in three distinct copies. Bottom panel shows the BGC alignment between the three instances of GCF-6 found within this genome. **e**) Representative isolates with and without staphyloxanthin encoding GCFs were grown on 20% BHI agar to visualize pigmentation production. **f**) Methanol extractions and wavelength absorption analysis were performed to identify signature peaks in the 400 to 500 nm range associated with the presence of staphyloxanthin in S. aureus^39,44^. Spectra are normalized to cell density.

To assess whether carriage of the *crt* operon translates to staphyloxanthin production in non-aureus species, we identified staphylococcal isolates cultivated from human skin with whole-genome sequences generated by our lab that encode a staphyloxanthin related GCF. We identified a rare instance of an *S. epidermidis* isolate with GCF-3 (only 2-3% of *S. epidermidis* genomes encode GCF-3), strain LK1136, and an *S. warneri* isolate with both GCF-3 and GCF-6, strain LK413. While we observe GCF-3 in all *S. warneri* genomes, the GCF is degraded encoding only the dehydrosqualene synthase gene, *crtM*, and the dehydrosqualene desaturase gene, *crtN* (Fig. 3a; Table S8). However, roughly half of the available *S. warneri* genomes (47.5%) encode GCF-6 in full and thus have two copies of *crtM* and *crtN*. We used long-read sequencing to generate near-complete genomes for *S. epidermidis* LK1136 and *S. warneri* LK413 and found that both strains encode GCF-3 within their chromosomes while GCF-6 is encoded on a plasmid within *S. warneri*. (Fig. S7b; Supplementary Text).

To test for phenotypic effects associated with strains carrying different staphyloxanthin GCFs, we assessed pigmentation levels in strains of *S. aureus, S. epidermidis*, and *S. warneri* with and without full *crt* operons (Table S9). Cells of *S. epidermidis* encoding GCF-3 are pigmented compared to strains without the cluster (Fig. 3e-f). Cells of *S. warneri* LK413 containing the complete *crt* operon (GCF-6) and additional copies of *crtMN* displayed the strongest golden pigmentation, confirmed as staphyloxanthin by spectrophotometry^41,47^ (Fig. 3f).

### *lsa*BGC provides a framework for mining for base-resolution novelty in BGCs

We have demonstrated how the *lsa*BGC suite can be used to identify homologous BGC instances into GCFs and subsequently uncover their evolutionary trends and phylogenetic distribution. With increasing metagenomic datasets becoming available, often from complex microbiomes, we developed *lsa*BGC-DiscoVary to directly profile BGCs and rapidly identify novel variation within GCF related genes from metagenomes (Fig. S2; Supplementary Text). The *lsa*BGC framework thus allows users to first identify all instances of GCFs in genomic assemblies to build a comprehensive database of known alleles for GCF genes and subsequently apply *lsa*BGC-DiscoVary to find novel SNVs within genes which are absent in the available assemblies. Further, because *lsa*BGC-DiscoVary extracts the subset of reads supporting the presence of novel SNVs, exhaustive methods to assess the validity of metagenomic SNV calling, which would be computationally intensive and impractical to perform for the full set of reads from a metagenome, are made accessible for further validation of novel SNVs.

We first assessed the sensitivity and specificity of *lsa*BGC-DiscoVary and found high concordance (>96% overlap) between novel SNVs reported by it when compared to an assembly-based approach using whole-genome sequencing readsets for 132 *M. luteus* isolates (Fig. S8; Table S10, S11; Supplementary Text). Next, to demonstrate *lsa*BGC-DiscoVary’s application in metagenomic datasets, its primary intended usage, we investigated the microdiversity of BGCs from *C. acnes*, the most abundant species at sebaceous sites, within healthy skin metagenomes generated by our lab^37^. We found that the BGC encoding for cutimycin, an anti-Staphylococcal thiopeptide, exhibited genomic signatures of horizontal, yet likely ancestral, acquisition by *C. acnes* and is now largely restricted to isolates belonging to either clades IB or III(Fig. 4ab, S9, S10a; Supplementary Text). Extensive metagenomic analysis demonstrated the reliability of *lsa*BGC-DiscoVary to type clade-specific alleles of genes within the cutimycin BGC in metagenomic datasets which correlated with strain presence inferences (Fig. 4cd, S10bc; Table S12). We used *lsa*BGC to further investigate novel SNVs detected within cutimycin genes and found that many were from the metagenomes of a single individual and belonged to clade III *C. acnes* (Fig. 4efgh). Through observation that the majority of novel SNVs from this individual also corresponded to cytosine to thymine changes, we discovered that clade III *C. acnes* have elevated rates of cytosine deamination, which coincides with the clade’s carriage of a truncated uracil DNA glycosylase, the enzyme responsible for the repair of such deaminations (Fig. 4ijk).

**Figure 4:**
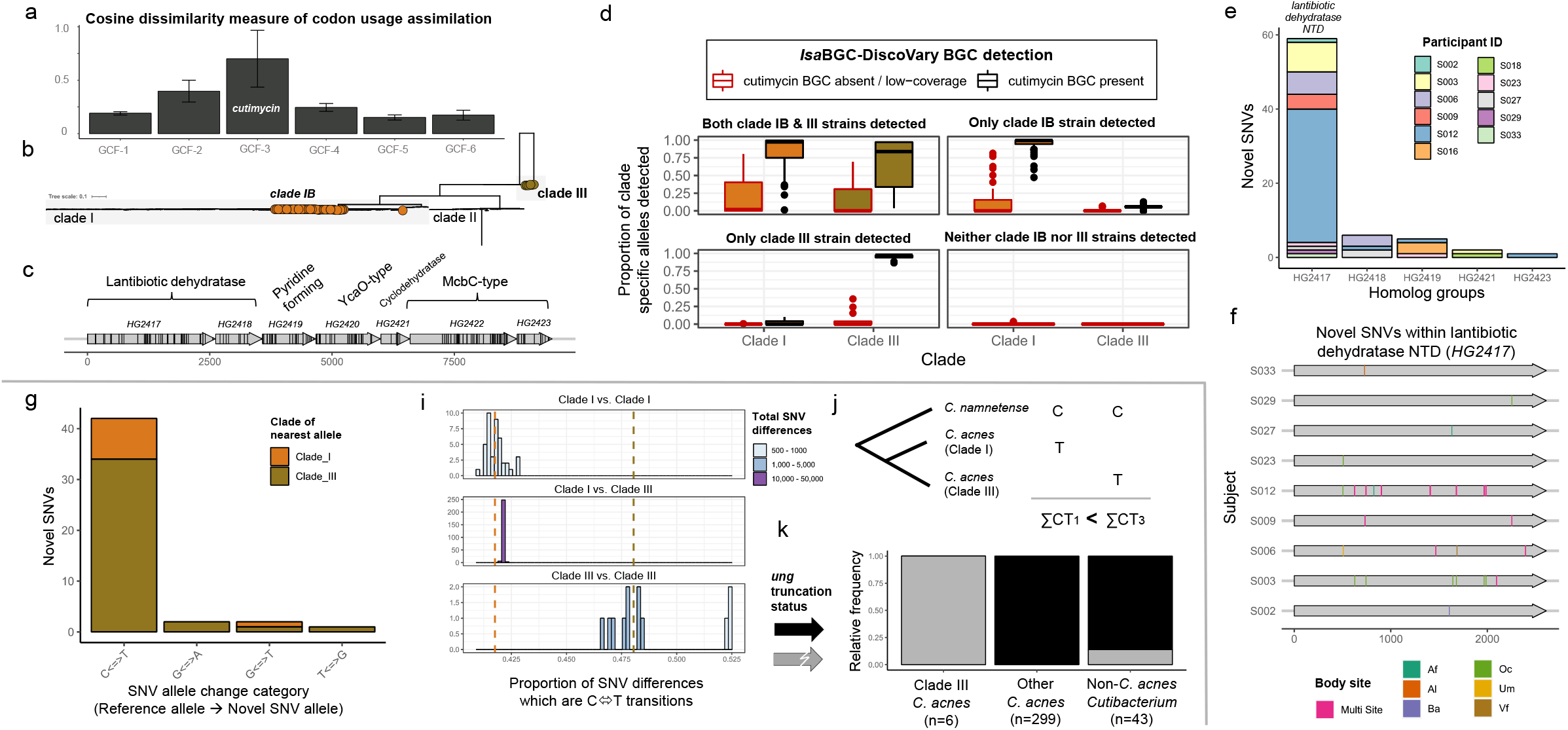
Mining for microdiversity within the cutimycin BGC leads to genetic insights for clade III *C. acnes*. **a**) Codon usage dissimilarity between BGC regions and background genomes shown for six common GCFs in *C. acnes*. **b**) The cutimycin encoding GCF is found primarily in clade I (subclade IB; orange) and clade III (tan) *C. acnes*. **c**) A schematic of the cutimycin encoding GCF. Lines in black correspond to sites of nearly-fixed differences between clades of C. *acnes*. **d**) *lsa*BGC-DiscoVary was used to identify the presence of the cutimycin GCF in 270 skin metagenomes. The proportion of alleles representative of clade IB or clade III C. *acnes* at sites of nearly fixed-differences was quantified. High-resolution strain quantification within metagenomes was similarly performed with StrainGE and used to overlay clade-specific typing results from *lsa*BGC-DiscoVary analysis. **e**) The distribution of novel SNVs identified by *lsa*BGC-DiscoVary within metagenomes, post filtering for removal of false positives, is shown across homolog groups and subjects. **f**) The location of novel SNVs is shown along the lantibiotic dehydratase N-terminal encoding gene (*HG2417*) for each subject. Coloring of the novel SNVs corresponds to which body-site(s) they were identified within. **g**) The clade distribution of reference genes on which novel SNVs were identified across the cutimycin GCF for subject S012 are shown individually for the predicted transitions or transversions. **i**) The proportion of differences which corresponded to cytosine to thymine or thymine to cytosine transitions along a core-genome alignment was calculated between pairs of *C. acnes* and categorized by clade comparisons. **j**) Comparative analysis of presumed directional deaminations from *C. namnetense*, treated as ancestral, to C. *acnes* clade IB and clade III were estimated using a multi-iteration sub-sampling based approach for sites along the core-genome alignment, whereby a greater number of suspected C to T transitions were estimated for clade III *C. acnes*. **k**) The presence of a complete uracil DNA glycosylase (*ung*), from clade I *C. acnes*, is shown across available *Cutibacterium* genomes.

### *lsa*BGC identifies novel SNVs within polyketide synthase catalytic domains

Phylogenetic and average nucleotide identity (ANI)^48^ based investigations revealed that *C. tuberculostearicum* exhibits high genomic similarity (ANI >88%) to four other species in GTDB (Fig. S4b, S11a), which we thus refer to, in aggregate, as the *C. tuberculostearicum* species complex. While metagenomic profiling found that this complex features the most prevalent *Corynebacterium* species within healthy skin microbiomes (Fig. S4b, Table S4), only 22 genomic assemblies were available for the complex at the time of this analysis. This clade thus represents an ideal taxonomic group to examine novel BGC microdiversity using *lsa*BGC-DiscoVary. We ran *lsa*BGC-DiscoVary systematically on six GCFs identified in the species complex, largely encoding for unknown metabolites, and found 40,019 putatively novel SNV instances across 105 homolog groups and 109 metagenomic samples. Of these, 5,474 instances were filtered because reads supporting their presence aligned with a higher score to other regions in a concatenated database of all *Corynebacterium* genomes from GTDB as compared to the reference database used for *lsa*BGC-DiscoVary. While ubiquitous, *C. tuberculostearicum* species complex members are present at very low abundances in some body-sites where metagenomic assembly would struggle to construct their genomes (Fig. S11b). We performed sample-specific metagenomic assemblies^49^ and found that at least 11,659 novel SNV instances detected by *lsa*BGC-DiscoVary would be missed by assembly-based investigation (Fig. 5a; Supplementary Text).

**Figure 5:**
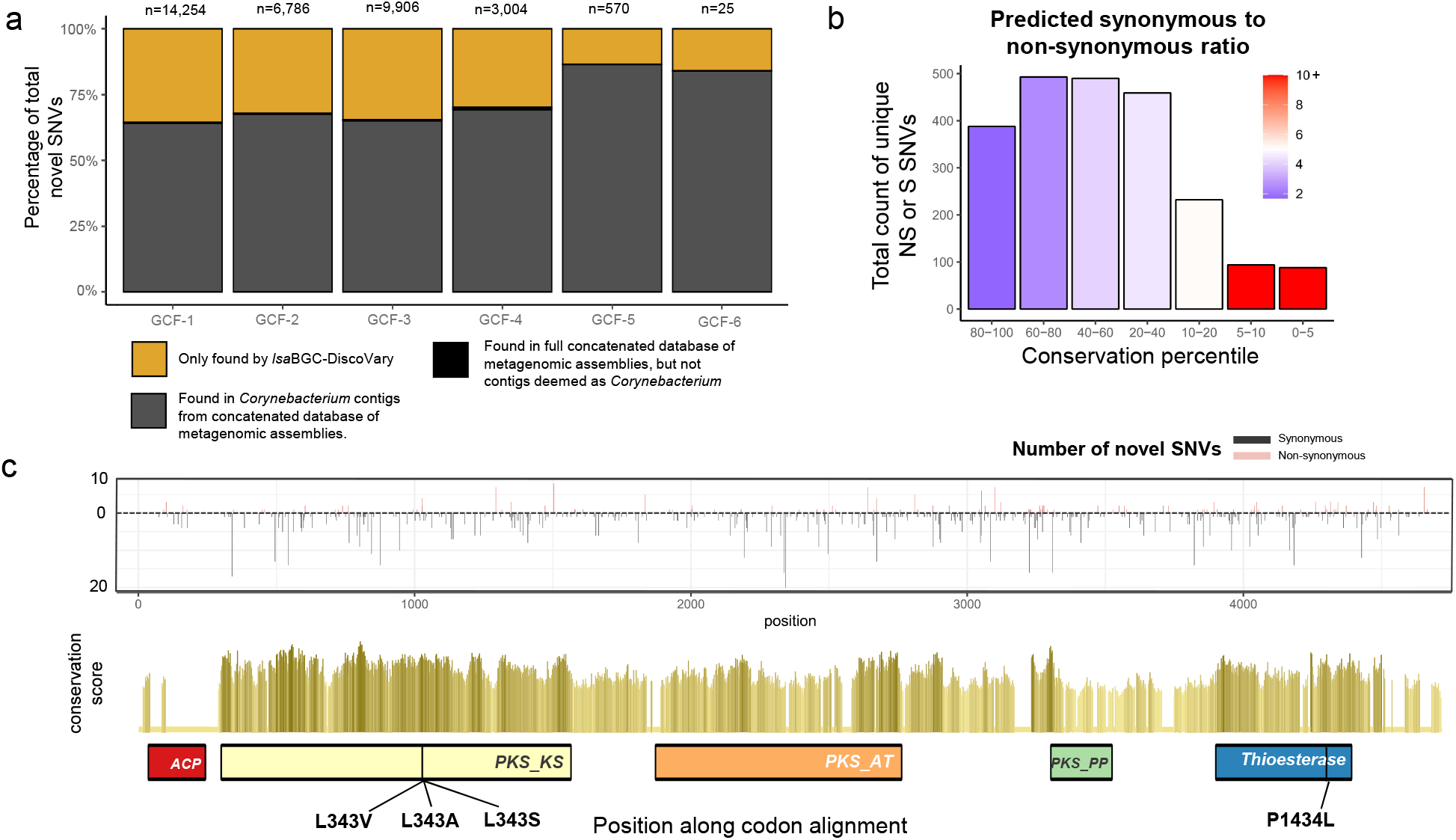
Thousands of novel SNVs identified within BGCs from the *C. tuberculostearicum* species complex. **a**) Assessment of novel SNVs identified by *lsa*BGC-DiscoVary, post-filtering, for each BGC from the *C. tuberculostearicum* species complex for presence within metagenomic assemblies. **b**) The number of novel SNVs is shown for ranges of conservation percentile across all homolog groups and metagenomes. The coloring corresponds to the ratio of synonymous to non-synonymous novel SNVs within each conservation range. **c**) Novel SNVs are shown for the polyketide synthase involved in mycolic acid biosynthesis. The top panel showcases the non-synonymous (up-facing) and synonymous (down-facing) novel SNVs found along the T1PKS gene, with heights corresponding to the number of metagenomes with the novel SNV. The middle panel shows the conservation scores colored by percentile ranges (lightest = least conserved sites; darkest = most conserved sites). Key domains common to T1PKS are shown underneath along with the location of predicted non-synonymous mutations within the top 5% of conserved sites for the gene. Two adjacent SNVs affecting the coding of the residue at position 343 were identified and these could result in a combined effect of encoding for alanine.

To further investigate the novel SNVs detected, we focused on 68 homolog groups overlapping or in close proximity to protocore regions of BGCs and further filtered based on standardized coverage metrics, retaining 5,802 instances of 2,343 unique novel SNVs. Novel SNVs were more likely to correspond to synonymous changes if they were found in multiple metagenomic samples (Fig. S11c). Further, metagenomic samples from the same subject at different body sites and from the same body sites across different subjects were more likely to share novel SNVs when compared to metagenomic samples from different body sites and different subjects (p<1E-8; two-sided Wilcoxon rank sum test) (Fig. S11d).

Finally, we expected and observed that the ratio of predicted synonymous to non-synonymous novel SNVs increases at deeply conserved sites along homolog groups (Fig. 5b; Table S13; Supplementary Text). The homolog group with the greatest number of novel SNVs, and novel, non-synonymous SNVs within highly conserved regions, was the PKS gene responsible for synthesis of mycolic acid, a major component of the cell wall in most *Corynebacterium* species recently shown to have immunomodulatory effects on the skin^30,50,51^. We identified and manually validated three non-synonymous SNVs along conserved sites of the PKS gene, including two adjacent SNVs predicted to affect one codon within the ketoacyl synthase domain, which catalyzes condensation^52^, and could alter reaction kinetics^53^ or scaffold structure^54^ (Fig. 5c, S12; Table S14).

## Discussion

The *lsa*BGC suite packages several fundamental, as well as novel, functionalities for high-throughput comparative and evolutionary analysis of BGCs from any taxa in a structured framework, such as the ability to detect GCFs within draft-quality genomic assemblies. Of practical relevance, it produces a comprehensive spreadsheet that includes annotation, conservation, and evolutionary statistics for homolog groups found in GCFs to enable efficient and high-throughput assessment by users. By incorporating GToTree^55^, *lsa*BGC will also automatically generate visualizations showing the distribution of GCFs across a species phylogeny.

One of the current limitations is that OrthoFinder based homology inference can be coarse and similar proteins encoded by recently duplicated genes can be collapsed into the same homolog group^13^. To mitigate this limitation, we clearly mark which homolog groups do not correspond to single-copy-orthologs within GCF contexts in resulting reports.

The metagenomic mining functionalities of *lsa*BGC further provide insight into the presence and sequence diversity of a specific taxa’s BGCs across diverse microbiomes. This is a particularly important feature for rare BGCs and taxa with a limited numbers of genomic assemblies available, such as taxa that are difficult to cultivate from complex metagenomes. In the future, the use of pan-genome alignment approaches to potentially increase the efficiency of *lsa*BGC for metagenomic identification of novel variants within BGC genes^56^ may be explored.

Here, we apply *lsa*BGC’s functionalities to major taxa commonly found in skin microbiomes as a proof of concept to demonstrate the potential to advance understanding of the diversity, evolutionary trends, and prevalence of both well studied and uncharacterized secondary metabolites. For example, we uncovered the ubiquity of the carotenoid staphyloxanthin across the genus of *Staphylococcus*, a molecule conferring the signature golden pigmentation in *S. aureus*^57,58^. These findings raise the hypothesis that staphyloxanthin, which has been identified primarily as a virulence factor conferring oxidative stress resistance in the pathogen *S. aureus*^44,59^, could similarly contribute to virulence in other staphylococcal species or serve additional unknown functions. Supporting this hypothesis, we observe that one staphyloxanthin encoding GCF appears ancestral to the skin-associated, multi-species clade which features both *S. aureus* and *S. epidermidis*. The observation that multiple copies of *crt* genes can be found within a single genome further suggests staphyloxanthin is of functional importance since evolutionary retention of paralogous genes has previously been shown to underlie ecological advantages^60,61^. Additionally, while this GCF is present in most skin-associated species, it is absent in greater than 95% of *S. epidermidis* genomes, suggesting large-scale loss within the species. It will thus be interesting to further explore how the loss of staphyloxanthin, if it contributes to oxidative stress resistance within *S. epidermidis*, coincides with the species’ reputation as a beneficial symbiont of host skin^34,62^.

Using the novel metagenomic mining functionality of *lsa*BGC, we identified novel SNVs distinguishing uncultured *C. acnes* and *Corynebacterium* strains from isolates with genomes available in public databases. With regards to *Corynebacterium*, we determined that some of the most common skin residing species^33,37^ belong to the *C. tuberculostearicum* species complex, which were widely distributed across skin metagenomes and highly abundant at moist body sites. Because this species complex is vastly underrepresented in public genomic databases and additional species and strains belonging to it likely remain to be discovered, application of *lsa*BGC-DiscoVary led to the identification of thousands of novel SNVs within the BGCs of the clade. Critically, some of these variants could have functional implications as they lie within highly conserved regions of important domains for natural product biosynthesis.

The comprehensive and open-source packaging of *lsa*BGC allows analyses to be applied to both public and proprietary microbial genomic databases. Besides accelerating fundamental evolutionary research of BGCs from diverse taxa, we envision *lsa*BGC will be particularly useful for natural product discovery and guiding mechanism of action studies. Evolutionary insight gathered from *lsa*BGC can be used as prior information to train algorithms in an emerging field using artificial intelligence to identify new antibiotics^63,64^. Further, an improved understanding of the relationship between intra-taxa genomic diversity and BGC content novelty at the base-resolution can also highlight which sub-lineages or sub-clades bear the largest reservoir of untapped secondary metabolic potential. Finally, we posit that identifying evolutionary trends of BGCs detected by highly-reliable rule-based approaches, such as antiSMASH^8^, can be used to assess the validity of BGC predictions from newly developed machine learning approaches, such as DeepBGC^11^ or GECCO^65^, which we now support usage of in *lsa*BGC.

## Acknowledgments

The authors gratefully acknowledge additional members of the Kalan lab, Dr. John-Demian Sauer, Dr. Jason Kwan, Dr. Karthik Anantharaman, Dr. Caroline Grunenwald, Dr. Chase Clark, and Dr. Milton Drott for helpful discussions.

## Author contributions

Conceptualization: RS, LRK. Computational Analysis and Software Development: RS. Investigation: RS, JC, LRK. Isolate Culturing and Sequencing: SS, MS, NLS, AS, LRK. Visualization: RS, JC. Supervision: LRK. Writing—original draft: RS, LRK. Writing—review & editing: RS, JC, SS, MS, RMS, NLS, CRC, AS, LRK.

## Materials and Methods

### Software availability

The *lsa*BGC software suite was developed in Python3 and R and is available on Github at: https://github.com/Kalan-Lab/lsaBGC. Algorithmic descriptions of programs can be found on the Github wiki and in the Supplementary Text document. A small test case of *Cutibacterium* genomes is included in the Github repository. Version 1.0 of the software was used for the analyses described in this study.

### An overview of the *lsa*BGC software suite

The first core program is *lsa*BGC-Cluster which clusters homologous and non-fragmented instances of BGCs, identified by antiSMASH from chromosomally complete genomic assemblies, into GCFs, similar to BiG-SCAPE^9^.Users can then generate a report to select for the most appropriate parameters for downstream analyses of interest. Because antiSMASH groups neighboring BGCs into hybrid gene clusters if they are located close to one another on the genome^7^, *lsa*BGC-Refiner has the option to manually separate hybrid clusters based on user-defined boundary homolog groups. Homologous instances of GCFs across all available genomic assemblies for the taxa of interest can then be identified through *lsa*BGC-Expansion (Figure S1b). Similar to ClusterFinder^10^, *lsa*BGC-Expansion uses profile Hidden Markov models (HMMs) to identify homologs of GCF associated genes in assemblies followed by a classical HMM approach to define BGC regions and their boundaries. Unlike ClusterFinder, *lsa*BGC-Expansion: (i) uses all coding genes associated with a GCF instead of being restricted to core protein domains, (ii) is specifically designed to account for assembly fragmentation through conditional assessment of predicted regions, and (iii) features a “quick mode” option for fast detection of BGCs using DIAMOND^66^.

The remaining four core programs are analytical and designed to be run on a single GCF for ease of parallelization: *lsa*BGC-See, *lsa*BGC-PopGene, *lsa*BGC-Divergence, and *lsa*BGC-DiscoVary. *lsa*BGC-See permits visualization of BGCs across a user-provided species tree or a BGC based phylogeny and can handle multiple fragmented BGC instances from the same genome, depicting them as neighboring leaves on phylogenies. *lsa*BGC-PopGene can then infer evolutionary and population genetics statistics for each homolog group associated with a GCF. It generates codon based alignments for each homolog group which can then be used as input for *lsa*BGC-Divergence and *lsa*BGC-DiscoVary. *lsa*BGC-PopGene also determines the consensus order and direction of homolog groups for a GCF across all BGC instances, which provides intuitive assessment of its reports. *lsa*BGC-Divergence calculates a comprehensive statistic between pairs of homologous BGCs from the same GCF but different genomes, which we refer to as Beta-RD, that captures how similar the BGC sequences are relative to what would be expected based on genome-wide similarity. Finally, *lsa*BGC-DiscoVary is a multi-functional program which can mine raw sequencing readsets from metagenomes or single genomes, for GCF instances and then reliably identify whether they possess novel single nucleotide variants (SNVs) yet to be observed in available genomic assemblies for a taxa (Figure S2).

### Automated workflows for running *lsa*BGC

To automate the application of these programs, we had initially developed three workflow programs called *lsa*BGC-AutoProcess.py, *lsa*BGC-AutoExpansion.py, and *lsa*BGC-AutoAnalyze.py. *lsa*BGC-AutoProcess.py is a preliminary workflow which does not execute any of the aforementioned core programs of the suite. It functions to take in a listing of genomic assemblies and perform gene-calling and basic annotation using Prokka^67^, annotate BGCs using antiSMASH^8^, and finally run OrthoFinder2^13^ for *de novo* delineation of homolog groups. It runs antiSMASH and OrthoFinder2 using the same Prokka output to ensure locus tag identifiers are able to be matched between the output of the two former programs (Figure S1a). *lsa*BGC-AutoExpansion.py is a wrapper of *lsa*BGC-Expansion.py which performs additional GCF instance identification in draft assemblies across all GCFs identified for a taxa. It is recommended that users run *lsa*BGC-AutoExpansion.py instead of *lsa*BGC-Expansion.py individually because the workflow additionally features a critical consolidation step in which BGC instances identified as potentially belonging to multiple GCFs are re-assessed and assigned to only the single best fitting GCF (Figure S1c). Finally, *lsa*BGC-AutoAnalyze.py is a major workflow which runs the analytical core programs across each GCF and creates a few consolidated reports after completion. This workflow starts by computing pairwise whole-genome similarity metrics using either CompareM^68^ or FastANI^48^ and then runs *lsa*BGC-See.py, *lsa*BGC-PopGene.py, *lsa*BGC-RelativeDivergence.py and, optionally, *lsa*BGC-DiscoVary.py for each GCF. At the end of the workflow, it generates consolidated report tables and visualizations from GCF-specific results for *lsa*BGC-PopGene.py and *lsa*BGC-RelativeDivergence.py. These workflows, similar to the core programs, are documented on the Github wiki.

### Recent additions to the *lsa*BGC suite

More recently, we have introduced two new core and workflow programs into the suite (release 1.2). The first is the core program *lsa*BGC-Ready.py, which serves to format existing BGC predictions in GenBank format and, optionally, BiG-SCAPE^9^ results, together with corresponding genomes, as input for downstream core programs in *lsa*BGC. Briefly, *lsa*BGC-Ready.py performs gene-calling on genomes using prodigal^69^, consolidates gene calling with those present in BGC GenBanks, performs annotation using KOfam and PGAP profile HMMs^70,71^, and can automatically run *lsa*BGC-Cluster.py and *lsa*BGC-AutoExpansion.py. We now recommend *lsa*BGC-Ready.py in place of the workflow *lsa*BGC-AutoProcess.py due to its simplified usage.

The second is an automated workflow for running *lsa*BGC on any bacterial species or genus called *lsa*BGC-Easy.py. Briefly, this workflow assesses GTDB^72^ for high-quality genomic assemblies belonging to a specified genus or species, downloads the genomes using ncbi-genome-download (https://github.com/kblin/ncbi-genome-download), performs gene-calling using prodigal^69^, constructs an alignment of single copy genes and phylogeny using GToTree^55^, dereplicates genomes based on the constructed alignment, and then runs *lsa*BGC-Ready.py, *lsa*BGC-Cluster.py, and *lsa*BGC-AutoAnalyze.py. The workflow also allows for users to include their own genomes belonging to the taxa, which might not yet be available on NCBI.

Analytical changes to the core and workflow programs since the initial release of *lsa*BGC (version 1.0) in relation to the most recent release of the software (version 1.3) have been minimal (Supplementary Text).

### Usage of genomic and metagenomic datasets

To overcome issues with misclassification and contamination in NCBI, we used GTDB (release 202) ^72^, to query for reliable genomic assemblies and respective species designations for the genera of *Corynebacterium, Cutibacterium,Micrococcus*, and *Staphylococcus* (Table S1). For *Micrococcus*, we further included 132 additional genomes of the type species *M. luteus* which our lab had sequenced (BioProject ID PRJNA803478) but were not yet uploaded to NCBI to be included in the GTDB release used. These isolates were classified as *M. luteus* by GTDB-tk^73^ and exhibited > 95% ANI to the NCBI representative genome for the species *M. luteus* NCTC 2665. A similar approach was used for gathering genome sets for species-level analyses performed (Table S1). For the *C. tuberculostearicum* species complex, an additional five genomes from NCBI were gathered, classified using GTDB-tk^73^, and incorporated into analysis if they belonged to one of the five species designated as belonging to the complex.

Because a single representative per species as provided by GTDB might not adequately capture the diversity of BGCs found across a genus, we instead employed a more granular approach to select for representative genomes for a genus. We performed dereplication using MASH^74^ with an estimated ANI cutoff of 0.99 to pair similar genomes and dynamically select representative genomes from each cluster based on assembly N50 (Table S1). This dereplication was performed using the script popSizeAndSampleSelector.py provided in the *lsa*BGC suite. Complete or chromosome level assemblies among the representative genomes were run through *lsa*BGC-AutoProcess.py and *lsa*BGC-Cluster.py to identify the core set of GCFs per genus. GCFs determined from the core set of genomes were comprehensively identified in the remaining genomes using *lsa*BGC-AutoExpansion.py, including those which were not deemed as representatives from the MASH based analysis to showcase scalability (e.g. all 15,055 *Staphylococcus* and not just the 229 representative genomes of which 77 corresponded to complete/chromosome level assemblies). To expedite population genetics analysis however, we only ran the downstream functional analytics of *lsa*BGC using the representative genomes for each genus. For species-level analyses, all genomes available were used for such downstream analyses and to comprehensively profile known alleles for genes in BGCs, which was critical to enable metagenomic mining for base-resolution novelty previously unseen in assemblies available for a species. For population level analyses within species, strain delineations were performed using the popSizeAndSampleSelector.py script in *lsa*BGC with strain groups determined from single-linkage clustering of pairs of genomes found to exhibit >98% ANI and >80% genomic content similarity by FastANI^48^.

For determining skin-associated species amongst the genera investigated in our study and to demonstrate mining functionalities with *lsa*BGC-DiscoVary, we used a paired-end metagenomics dataset sequenced on the Illumina NovaSeq where individual samples corresponded to microbiome surveys of one of eight body sites for one of 34 participants, collected in the state of Wisconsin (BioProject ID PRJNA763232)^37^. As previously described^37^, adapter removal, quality filtering, human sequence decontamination, and tandem repeat removal were performed using fastp (v0.21.0)^75^ and KneadData (v0.8.0) (https://huttenhower.sph.harvard.edu/kneaddata/).

### Genomic assessment for staphyloxanthin carriage in the lab isolate collection

We had previously constructed draft genomic assemblies for 44 *Staphylococcus* isolates from skin (BioProject ID PRJNA803478; BioProject ID PRJNA830888). Illumina sequencing was performed at the Microbial Genome Sequencing Center (MiGS). Briefly, assemblies were generated using Unicycler^76^ with default settings after quality and adapter trimming using fastp^75^. The set of 44 *Staphylococcus* genomes were searched for carriage of staphyloxanthin encoding GCFs identified in our study using *lsa*BGC-AutoExpansion (Table S9).

### Library preparation, sequencing and genomic assembly

#### *Reconstruction of Illumina draft assemblies for 132* M. luteus

We assessed the performance of multiple programs in the *lsa*BGC suite using a set of genomic assemblies for 132 *M. luteus* isolates which were cultured from skin. For these isolates, assemblies were reconstructed from raw-reads (BioProject ID PRJNA803478) and were uploaded to NCBI’s Genbank database (Table S1). Briefly, Illumina sequencing reads were processed using fastp (v. 0.20.1)^75^ to trim for Poly-G tail artifacts and TrimGalore (v. 0.6.5)^77^ was additionally used to detect and remove adapters. Afterwards, assembly was performed using Unicycler (v. 0.4.6)^76^ with default settings.

#### Construction of Illumina+Nanopore hybrid assemblies for staphyloxanthin, producing S. warneri LK413 and S. epidermidis LK1136

To learn the genomic location of staphyloxanthin encoding GCFs in *S. epidermidis* LK1136 and *S. warneri* LK413, we performed additional long-read sequencing using Oxford Nanopore Technologies (ONT). Library construction, using a PCR-free ligation method, and sequencing were performed at the Microbial Genome Sequencing Center (MiGS). Hybrid assemblies were constructed using a modified version of the Hybrid Assembly workflow in seQuoia (https://github.com/broadinstitute/seQuoia), Illumina reads, used for construction of the initial draft genomic assemblies for the isolates, were first reprocessed using fastp (v. 0.20.1)^75^ to trim for Poly-G tail artifacts and TrimGalore (v. 0.6.5)^77^ was subsequently used to detect and remove adapters. Similarly, ONT FASTQ reads, provided by MIGS after basecalling, were filtered for potential adapters using PoreChop (v. 0.2.3) (https://github.com/rrwick/Porechop) and filtered to retain only reads greater than 3 kb in length as well as subsampled to 300 mb (while being inclusive of all reads greater than 20 kb) using the script fastqfilter.py included in seQuoia. Afterwards, hybrid-assemblies were constructed using Unicycler (v. 0.4.6)^76^ with default settings and additional Pilon (v. 1.23)^78^ polishing was iteratively performed by aligning ONT reads^79^ as well as the Illumina reads^80^ to assemblies and refining them, allowing for a maximum of 10 iterations. Quality assessment of the two near-complete assemblies was performed using GAEMR (https://github.com/broadinstitute/GAEMR) (Supplementary Text).

### Experimental validation of staphyloxanthin production in non-aureus *Staphylococcus*

Isolates were grown overnight at 37°C on tryptone soy agar plates and inoculated into tryptone soy broth at a concentration of 1×10^6^ CFU/mL. Liquid cultures were grown in shaking conditions at 37°C for 24 hours. 10 mL of the liquid culture was then pelleted and washed twice with phosphate-buffered saline before pigment extraction in 1 mL of methanol at 55°C for 15 mins. Spectral scans from 300 nm to 600 nm were taken using a plate reader (EPOCH2, Biotek, Winooski, VT). Absorbance readings were blanked with methanol and normalized to liquid culture cell density based on OD600. Spectra shown are representative of at least two biological replicates.

### MASH distance neighbor-joining tree construction

Average nucleotide identity estimates between pairs of genomes were computed with MASH^74^ and used to create a distance matrix. The distance matrix was then used to construct a neighbor-joining tree with the ape library in R^81^.

### Ribosomal and BGC phylogeny construction

#### Ribosomal phylogeny construction

Ribosomal phylogenies for each species were constructed based on a concatenated alignment of 16 ribosomal proteins^82^. HMMER (v3) was used to search for profile HMMs of each ribosomal protein in the predicted proteomes of samples of interest belonging to a particular taxonomic group. Afterwards, the best matching hits for each sample based on alignment E-value were identified and used to construct individual alignments for each ribosomal protein with the L-INS-I method in MAFFT^83^. Protein alignments were then converted to codon alignments using PAL2NAL^84^ and subsequently concatenated together. Samples which featured > 25% gaps in the concatenated alignment were filtered after which core SNV sites were extracted and used for constructing a maximum likelihood phylogeny with RAxML^85^, specifying a GTRCAT model and performing 100 bootstraps. Where needed, ribosomal phylogenies were subset to showcase relationships between select samples using PareTree (http://emmahodcroft.com/PareTree.html; v1.0.2).

#### Staphyloxanthin phylogeny construction

A maximum likelihood phylogeny was constructed from a concatenated protein alignment of CrtM and CrtN amongst representative *Staphylococcus* genomes to understand the relationship between different GCFs encoding for staphyloxanthin. Proteins designated as HG0001876 (CrtM) and HG0001904 (CrtN) were gathered for each staphyloxanthin encoding GCF found in each sample and aligned using the L-INS-I method in MAFFT^83^. To account for assembly fragmentation resulting in multiple proteins from a sample being assigned to the same homolog group for a specific GCF, a consensus sequence was determined and multi-allele sites were replaced as gaps. A concatenated alignment of such consensus sequences was subsequently constructed and used for phylogeny construction with RAxML using a PROTCAT model with JM selected as the best amino-acid replacement matrix through the automatic maximum likelihood model selector^85^. The phylogeny of CrtMN sequences exhibited high correspondence to GCF delineations with the exception of a single *S. lugdunensis* BGC which is likely misclassified as GCF-17 and should be GCF-3. Upon further examination, it was identified that the reason for the misclassification was likely due to the absence of mevalonate pathway genes, classified as a type III polyketide synthase (T3PKS), within *S. lugdunensis* which are found downstream of the *crt* operon in all other species with GCF-3 (Figure S7b).

#### Pyrazinone phylogeny construction

A maximum likelihood phylogeny was constructed from a concatenated protein alignment of PznA and PznB amongst representative *Staphylococcus* genomes using the same methods as described for phylogeny construction from CrtMN proteins.

### Designation of species belonging to *Mammaliicoccus*

The *Staphylococcus_A* genus designation in GTDB R202 was largely concordant with the recent reclassification of five species from *Staphylococcus* to *Mammaliicoccus*^46^. In the latest GTDB R207 release, this genus has now been reclassified to *Mammaliicoccus*. The recent update to taxonomic names also includes the reclassification of species *S. pasteuri_A* to *M. fleurettii*.Notably, we do not regard *S. schleiferi* as part of *Mammaliicoccus* because it is categorized as *Staphylococcus* by GTDB (both R202 and R207) and groups within the *Staphylococcus* genus in our ribosomal protein based maximum-likelihood phylogeny. Similar to our assessment, the reclassification of this species to *Mammaliicoccus* was recently questioned^86^.

### Ubiquity assessment of skin residing species and strains of *Staphylococcus* and *Corynebacterium*

We searched for the presence of representative *Staphylococcus* and *Corynebacterium* strains in 270 skin metagenomes from 34 participants across 8 body sites using StrainGST^38^. To construct the database of distinct representative genomes for each genus, the protocol described in the StrainGE documentation was followed. *S. aureus* genomes 16405 and 278 were dropped from the analysis because at the time of downloading them, the plasmid and chromosome were switched. Briefly, genomes were downloaded from NCBI using ncbi-genome-download (https://github.com/kblin/ncbi-genome-download) and dereplication was performed using built-in StrainGE functionalities. As some representative genomes were not part of the GTDB release R202, GTDB-tk^73^ was used to annotate such genomes and perform species classification. We considered a species as skin-associated if it was found in at least 10 of the 270 skin metagenomes, regardless of the abundance (Table S4). Certain skin-associated species from each genus were also identified in the 19 negative control metagenomic samples, which included air-swab, bench, desk, and door-handle samplings (Table S4). Because we expect heavy overlap in species content between these environmental samples and skin microbiome samples, we still considered such species as skin-associated if they were found in 10 or more skin metagenomes.

We found the majority of skin-associated *Staphylococcus* belonged to the *S. epidermidis/aureus* clade, as has previously been noted^42^. Similarly, most of the skin-associated *Corynebacterium* belonged to *C. tuberculostearicum* and it’s close neighboring species (including *C. kefirresidentii*,*C. aurimucosum* type E, C. sp. 900539985, *C. tuberculostearicum*,*C. tuberculostearicum* type C), which we denote in this study as the *C. tuberculostearicum* species complex. The minimal ANI separating genomes belonging to this species complex was determined to be 88.59% using FastANI^48^. To assess whether any of the 24 new *Corynebacterium* species reported by Saheb Kashaf et al. 2022^36^ belonged to this species complex, FastANI was used to look at their ANI similarity to genomes from the clade and regard them as additional members if they exhibited 88.59% ANI or greater. Of note, many additional *Corynebacterium* species have been routinely isolated from the skin^37^ which we did not regard as skin-associated based on our criteria in this study. In particular, the species of *C. amycolatum*(detected in 4 metagenomes), *C. kroppenstedtii* (detected in 4 metagenomes), and *C. simulans* (detected in 5 metagenomes) are commonly found on skin.

To estimate the relative abundance of *Cutibacterium* (see Supplementary Text section “Identification of *Cutibacterium* Strains within Metagenomes and Determination of Near-Fixed Sites Differentiating Cutimycin Biosynthesis Alleles between Clade I and Clade III *C. acnes*”)and *Corynebacterium* strains within metagenomic samples we normalized the estimated relative abundances for strains (*rapct*) by the estimated relative abundance of the genus (*pan*%) as reported by StrainGST^38^.

### Identification of homolog groups associated with skin residing species of *Staphylococcus* and *Corynebacterium*

We developed a script called crawlingFisher.py, provided in the *lsa*BGC suite, which tests for enrichment or depletion of homolog group presence amongst genomes in focal clades using Fisher’s exact test. Provided with a phylogenetic tree and matrix specifying the presence of homolog groups across genomes, the program automatically performs such tests for each homolog group at each inner-node of the phylogeny. Multiple testings correction was performed comprehensively at the end using Benjamini-Hochberg false discovery rate and cases were considered statistically significant if the adjusted p-value was less than 0.05. Results were further filtered to retain only homolog groups which were present in at least 80% of genomes under the focal node and at most 20% of other genomes or, alternatively, at most 20% of genomes under the focal node and at least 80% of other genomes. Finally, only results pertaining to focal nodes where at least 80% of genomes were classified as skin-associated (see section “Ubiquity Assessment of Skin Residing Species and Strains of *Staphylococcus* and *Corynebacterium”*)were reported (Table S5).

### Assessing the divergence of BGCs relative to their background genomic contexts

BGC divergence does not always track with background genomic divergence and speciation^14^. The program *lsa*BGC-Divergence allows for easy assessment of how BGC divergence compares to whole-genome divergence, primarily through computing a statistic we refer to as Beta-Relative-Divergence (Beta-RD) between pairs of genomes for a single GCF. The Beta-RD statistic is simply the average amino or nucleotide identity between homologous instances of proteins for a GCF observed in two genomes normalized by the average amino or nucleotide identity observed between the full genomes. *lsa*BGC allows users to specify whether amino or nucleotide divergence should be used and automated workflows (see section Automated Workflows for Running *lsa*BGC) can be used to compute pairwise AAI or ANI between genomes using CompareM (https://github.com/dparks1134/CompareM) or FastANI^48^, respectively.

To better understand how distributions of Beta-RD differ across GCFs, in this study, we applied a Bayesian hierarchical model to perform shrinking analysis and appropriately account for differences in the number of genomes GCFs are found (Figure S4cd; Table S6). For computational efficiency, a maximum of 500 Beta-RD observations were used for each GCF. Posterior predictions for Beta-RD were made using rstan (Stan Development Team 2021) with the following hierarchical model:

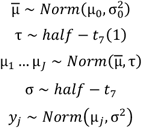

where for each GCF j, the posterior Beta-RD distribution is sampled as y_j_ using Markov chain Monte Carlo with four chains, each featuring 2,000 iterations, including a burnin of 1,000 iterations. A hierarchical prior was specified for *μ*_j_, which represents a GCF-specific baseline. The values of *μ*_0_ and *σ*_0_ were set to the respective mean and standard deviation of raw Beta-RD values across all GCFs. R and STAN code used for this analysis are available within the scripts subdirectory of the *lsa*BGC software package.

### Scrutinization of putative novel variants for *lsa*BGC-DiscoVary application to cutimycin in *C. acnes* and the comprehensive set of GCFs for the *C. tuberculostearicum* species complex

By default, *lsa*BGC-DiscoVary.py only considers reads aligning at 95% identity to reference gene sequences and avoids reporting novel variants on genes if either annotation or coverage (relative to other genes in the BGC context) suggests they might be MGEs. The program also automatically extracts the subset of reads supporting the existence of novel variants, which can then be further screened with high stringency filters using otherwise computationally expensive methods. This can involve mapping reads supporting putative novel SNVs to a representative database of whole-genomes to observe for higher quality alignments or checking whether reads are classified as a particular taxonomy by Kraken2 (v2.0.8-beta)^87^. As described previously^37^, we used a customized database for Kraken2 analysis constructed from chromosomally complete bacterial, viral, archaeal, fungal, protozoan genomes as well as the human genome available on RefSeq, where plasmid sequences were distinguished to alleviate potential misclassifications of reads to a particular taxonomy.

For reads supporting putative novel variants in the cutimycin BGC of *C. acnes*, we required them to be classified as *Cutibacterium* or *Priopionibacterium* by Kraken2^87^ and to not map with a higher alignment score to a comprehensive database of all *Cutibacterium* genomes than was observed for the alignment to the reference gene sequence in *lsa*BGC-DiscoVary.py (Figure S2c). We generally do not recommend screening variant-supporting reads for an appropriate taxonomic match with Kraken2 as it can generally lead to false negatives (on average only 92.8% of read-pairs per sample were taxonomically classified and misclassification can also occur; Figure S12) but performed this here to provide further assurance that novel variants discovered are reliable. Standardized coverage metrics within *lsa*BGC-DiscoVary.py’s novel SNV report were further used to conservatively filter sites and variants if they exhibited site or allele-specific coverage values greater than two median absolute deviations from the median coverage for the entire BGC.

For reads supporting putative novel variants within GCFs of the *C. tuberculostearicum* species complex, we required them to not align with a higher alignment score to a comprehensive database of all *Corynebacterium* genomes. We did not require reads to be classified by Kraken2 as *Corynebacterium* because genomes from the species complex are likely to be poorly represented in the Kraken2 database. We did however further validate that reads supporting the existence of variants at conserved sites in the mycolic acid biosynthesis gene mapped best to the *C. tuberculostearicum* species and confirmed our suspicion that they could be misclassified by Kraken2 given the representation in the underlying database (Figure S12). This *ad hoc* analysis was performed using paired-end information when mapping in Bowtie2 and accounting for concordant read pair alignments. While many reads supporting the existence of novel SNVs in the biosynthesis gene were classified as *C. acnes* by Kraken2, we found that none of the reads concordantly aligned to the comprehensive *Cutibacterium* genomes Bowtie 2 database whereas 85.07% concordantly aligned to the comprehensive *Corynebacterium* genomes Bowtie 2 database.

Note, the reasoning behind mapping SNV supporting reads to comprehensive genomic databases to purge false positive SNV calls, is that if a read truly aligns best to the BGC-associated homolog group where it was used to call a SNV, then it should at most generate a mapping to this comprehensive database of equal mapping quality. We align paired-end reads individually as was performed in *lsa*BGC-DiscoVary.py to have mapping scores be comparable with those from the original alignment. Furthermore, because our initial database within *lsa*BGC-DiscoVary.py consisted of allelic representatives of homolog groups, we realized that reads mapping to the start and end of genes could result in a lower mapping score than if a larger genomic context was provided. To ensure we do not remove such reads mapping to gene edges, scrutunizeNovelSNVSupport.py, the program we developed to perform Kraken 2 and genomic database alignment based filtering of novel SNV reports from *lsa*BGC-DiscoVary.py, also compares the reference sequences when it finds a better Bowtie 2 mapping for SNV-supporting reads in genomic databases. If the reference sequence in the genomic database encompasses the reference sequence in the original alignment to the *lsa*BGC-DiscoVary.py homolog group database and the read was previously noted to map to a gene edge, then reads are still retained and SNV read support is not decremented. Five retained and supportive reads are required for an SNV to be featured in the filtered report produced by the program.

### *lsa*BGC-DiscoVary based analyses of BGCs from the *C. tuberculostearicum* species complex

For the following analyses, we used 5,802 instances of 2,343 unique novel SNVs found in 68 homolog groups which were core or adjacent to core biosynthesis machinery of BGCs from the *C. tuberculostearicum* species complex. These novel SNVs had site and allele-specific coverage values less than two median absolute deviations from the median coverage for the entire BGC.

#### Comparing SNVs shared across body sites and subjects

To comprehensively assess whether novel SNVs were shared more frequently for microbiomes of the same body site or from the same individual, we performed a pairwise, multi-iteration resampling analysis. For each pair of metagenomes, where each sample had at least 30 novel SNVs, we calculated the average Jaccard similarity across 1,000 simulations in which we randomly drew sets of 30 SNVs from each metagenomic sample to control for sequencing depth. Pairs of metagenomes were classified as one of three categories: “different body site, different participant”, “different body site, same participant”, and “same body site, different participant” (Figure S11c). A two-sided Wilcoxon rank sum test for differences between the three distributions of average Jaccard similarities revealed that there was a statistically significant difference between “different body site, same participant’ and “same body site, different participant” (*p*=6.01E-3). Of greater relevance, there were statistically strong significant differences between the “different body site, different participant” distribution and the two other distributions: “different body site, same participant” and “same body site, different participant” (*p*=3.67E-09 and *p*=1.39E-116, respectively). Thus, pairs of metagenomes were more likely to share novel SNVs if they were either from the same body site or same participant compared to if they were from different participants and different body sites. Two metagenomic samples for participant S002 were not used because they corresponded to a resampling of two of their body sites and we did not have enough of such samples to pursue a temporal analysis. We assessed this testing using different considerations/cutoffs for: (i) the number of novel SNVs needed for a metagenome to be considered in the analysis and (ii) whether or not to account for singleton SNVs. In all cases, statistical testing for differences between distributions of pairwise metagenome categories yielded similar conclusions.

#### Assessing trends for novel SNV predicted synonymous to non-synonymous rate with metagenome ubiquity and site conservation

Using reports from *lsa*BGC-DiscoVary.py, we performed a systematic prediction of whether novel SNVs identified on the 68 homolog groups nearby or overlapping with protocore regions of BGCs in *C. tuberculostearicum* corresponded to synonymous or non-synonymous substitutions. As described in other sections, SNV alleles were compared to corresponding alleles observed at the site on the reference gene sequences which SNVs were called upon. We aimed to check whether the rate of synonymous to non-synonymous novel SNVs showed a relationship to: (i) how common SNVs were across metagenomic samples and (ii) conservation levels across protein sequences.

#### Novel SNV predicted synonymous to non-synonymous rate increases with SNV metagenome ubiquity amongst metagenomic samples

The occurrence of novel SNVs, a novel allele observed at a specific site in the multiple sequence codon alignment of a particular homolog group, was tabulated across metagenomic samples. While novel SNVs found in a single metagenome were only twice as likely to correspond to a synonymous change as opposed to a non-synonymous change, more prevalent novel SNVs, found in ten or more samples, were eight times more likely to correspond to a synonymous change (Figure S11b).

#### Novel SNV predicted synonymous to non-synonymous rate increases at conserved sites in protein sequences

We searched RefSeq’s bacterial NR database for remote homologs of the 68 homolog groups within or adjacent to protocore regions of BGCs. The top 20 homologs which belonged to classified bacterial species outside of the *Corynebacterium* genus or *Corynebacteriales* family were identified using HMMER3 with profile-HMMs gathered from *lsa*BGC-Expansion for each homolog group and an E-value threshold of 1E-20 (Table S13). The sequences of these representative homologs were added to protein alignments for homolog groups in *lsa*BGC-PopGene results using the ‘--add’ option in MAFFT^83^. Afterwards, we used the alignments for scoring protein sequence conservation with Jensen-Shannon divergence^88^. Conservation scores were used to determine conservation percentiles of sites along protein alignments for individual homolog groups. Rates of predicted synonymous to non-synonymous novel SNVs were calculated for each conservation percentile across all homolog groups. For the three SNVs predicted to encode for non-synonymous differences and also in the top five percentile of conserved sites along the mycolic acid PKS, we inspected their codon contexts to check that there were no additional variants which, in aggregate, might result in a synonymous change (Table S14).

## References

1. Katz, L. & Baltz, R. H. Natural product discovery: past, present, and future. J. Ind. Microbiol. Biotechnol. 43, 155–176 (2016).

2. Watve, M. G., Tickoo, R., Jog, M. M. & Bhole, B. D. How many antibiotics are produced by the genus Streptomyces? Arch. Microbiol. 176, 386–390 (2001).

3. Hopwood, D. A. Streptomyces in nature and medicine: The antibiotic makers. (Oxford University Press, 2007).

4. Chevrette, M. G. et al. The antimicrobial potential of Streptomyces from insect microbiomes. Nat. Commun. 10, 516 (2019).

5. Waterworth, S. C. et al. Horizontal Gene Transfer to a Defensive Symbiont with a Reduced Genome in a Multipartite Beetle Microbiome. MBio 11, (2020).

6. Drott, M. T. et al. Microevolution in the pansecondary metabolome of Aspergillus flavus and its potential macroevolutionary implications for filamentous fungi. Proc. Natl. Acad. Sci. U. S. A. 118, (2021).

7. Steinke, K., Mohite, O. S., Weber, T. & Kovács, Á. T. Phylogenetic Distribution of Secondary Metabolites in the Bacillus subtilis Species Complex. mSystems 6, (2021).

8. Blin, K. et al. antiSMASH 6.0: improving cluster detection and comparison capabilities. Nucleic Acids Res. 49, W29–W35 (2021).

9. Navarro-Muñoz, J. C. et al. A computational framework to explore large-scale biosynthetic diversity. Nat. Chem. Biol. 16, 60–68 (2020).

10. Cimermancic, P. et al. Insights into secondary metabolism from a global analysis of prokaryotic biosynthetic gene clusters. Cell 158, 412–421 (2014).

11. Hannigan, G. D. et al. A deep learning genome-mining strategy for biosynthetic gene cluster prediction. Nucleic Acids Res. 47, e110 (2019).

12. Kautsar, S. A., van der Hooft, J. J. J., de Ridder, D. & Medema, M. H. BiG-SLiCE: A highly scalable tool maps the diversity of 1.2 million biosynthetic gene clusters. Gigascience 10, (2021).

13. Emms, D. M. & Kelly, S. OrthoFinder: phylogenetic orthology inference for comparative genomics. Genome Biol. 20, 238 (2019).

14. Chase, A. B., Sweeney, D., Muskat, M. N., Guillén-Matus, D. G. & Jensen, P. R. Vertical Inheritance Facilitates Interspecies Diversification in Biosynthetic Gene Clusters and Specialized Metabolites. MBio 12, e0270021 (2021).

15. Chevrette, M. G. et al. Evolutionary dynamics of natural product biosynthesis in bacteria. Nat. Prod. Rep. 37, 566–599 (2020).

16. Sélem-Mojica, N., Aguilar, C., Gutiérrez-García, K., Martínez-Guerrero, C. E. & Barona-Gómez, F. EvoMining reveals the origin and fate of natural product biosynthetic enzymes. Microb. Genom. 5, (2019).

17. Felnagle, E. A., Rondon, M. R., Berti, A. D., Crosby, H. A. & Thomas, M. G. Identification of the biosynthetic gene cluster and an additional gene for resistance to the antituberculosis drug capreomycin. Appl. Environ. Microbiol. 73, 4162–4170 (2007).

18. Del Carratore, F. et al. Computational identification of co-evolving multi-gene modules in microbial biosynthetic gene clusters. Commun Biol 2, 83 (2019).

19. Crits-Christoph, A., Bhattacharya, N., Olm, M. R., Song, Y. S. & Banfield, J. F. Transporter genes in biosynthetic gene clusters predict metabolite characteristics and siderophore activity. Genome Res. (2020) doi:10.1101/gr.268169.120.

20. Mungan, M. D. et al. ARTS 2.0: feature updates and expansion of the Antibiotic Resistant Target Seeker for comparative genome mining. Nucleic Acids Res. 48, W546–W552 (2020).

21. Crits-Christoph, A., Diamond, S., Butterfield, C. N., Thomas, B. C. & Banfield, J. F. Novel soil bacteria possess diverse genes for secondary metabolite biosynthesis. Nature 558, 440–444 (2018).

22. Sugimoto, Y. et al. A metagenomic strategy for harnessing the chemical repertoire of the human microbiome. Science 366, (2019).

23. Andreu, V. P. et al. A systematic analysis of metabolic pathways in the human gut microbiota. bioRxiv 2021.02.25.432841 (2021) doi:10.1101/2021.02.25.432841.

24. Pereira-Flores, E. et al. Mining metagenomes for natural product biosynthetic gene clusters: unlocking new potential with ultrafast techniques. bioRxiv 2021.01.20.427441 (2021) doi:10.1101/2021.01.20.427441.

25. Chevrette, M. G. & Handelsman, J. Needles in haystacks: reevaluating old paradigms for the discovery of bacterial secondary metabolites. Nat. Prod. Rep. 38, 2083–2099 (2021).

26. Zipperer, A. et al. Human commensals producing a novel antibiotic impair pathogen colonization. Nature 535, 511–516 (2016).

27. Claesen, J. et al. A Cutibacterium acnes antibiotic modulates human skin microbiota composition in hair follicles. Sci. Transl. Med. 12, (2020).

28. Nakatsuji, T. et al. Antimicrobials from human skin commensal bacteria protect against Staphylococcus aureus and are deficient in atopic dermatitis. Sci. Transl. Med. 9, (2017).

29. Wyatt, M. A. et al. Staphylococcus aureus nonribosomal peptide secondary metabolites regulate virulence. Science 329, 294–296 (2010).

30. Ridaura, V. K. et al. Contextual control of skin immunity and inflammation by Corynebacterium. J. Exp. Med. 215, 785–799 (2018).

31. Swaney, M. H. & Kalan, L. R. Living in Your Skin: Microbes, Molecules, and Mechanisms. Infect. Immun. 89, (2021).

32. Grice, E. A. et al. Topographical and temporal diversity of the human skin microbiome. Science 324, 1190–1192 (2009).

33. Oh, J. et al. Temporal Stability of the Human Skin Microbiome. Cell 165, 854–866 (2016).

34. Byrd, A. L., Belkaid, Y. & Segre, J. A. The human skin microbiome. Nat. Rev. Microbiol. 16, 143–155 (2018).

35. Acosta, E. M. et al. Bacterial DNA on the skin surface overrepresents the viable skin microbiome. bioRxiv 2021.08.16.455933 (2021) doi:10.1101/2021.08.16.455933.

36. Saheb Kashaf, S. et al. Integrating cultivation and metagenomics for a multi-kingdom view of skin microbiome diversity and functions. Nat Microbiol 7, 169–179 (2022).

37. Swaney, M. H., Sandstrom, S. & Kalan, L. R. Cobamide sharing drives skin microbiome dynamics. bioRxiv 2020.12.02.407395 (2021) doi:10.1101/2020.12.02.407395.

38. van Dijk, L. R. et al. StrainGE: a toolkit to track and characterize low-abundance strains in complex microbial communities. Genome Biol. 23, 74 (2022).

39. Zimmermann, M. & Fischbach, M. A. A family of pyrazinone natural products from a conserved nonribosomal peptide synthetase in Staphylococcus aureus. Chem. Biol. 17, 925–930 (2010).

40. Tauch, A. et al. Complete genome sequence and analysis of the multiresistant nosocomial pathogen Corynebacterium jeikeium K411, a lipid-requiring bacterium of the human skin flora. J. Bacteriol. 187, 4671–4682 (2005).

41. Pelz, A. et al. Structure and biosynthesis of staphyloxanthin from Staphylococcus aureus. J. Biol. Chem. 280, 32493–32498 (2005).

42. Balibar, C. J., Shen, X. & Tao, J. The mevalonate pathway of Staphylococcus aureus. J. Bacteriol. 191, 851–861 (2009).

43. Grice, E. A. et al. A diversity profile of the human skin microbiota. Genome Res. 18, 1043–1050 (2008).

44. Clauditz, A., Resch, A., Wieland, K.-P., Peschel, A. & Götz, F. Staphyloxanthin plays a role in the fitness of Staphylococcus aureus and its ability to cope with oxidative stress. Infect. Immun. 74, 4950–4953 (2006).

45. Seel, W. et al. Carotenoids are used as regulators for membrane fluidity by Staphylococcus xylosus. Sci. Rep. 10, 330 (2020).

46. Madhaiyan, M., Wirth, J. S. & Saravanan, V. S. Phylogenomic analyses of the Staphylococcaceae family suggest the reclassification of five species within the genus Staphylococcus as heterotypic synonyms, the promotion of five subspecies to novel species, the taxonomic reassignment of five Staphylococcus species to Mammaliicoccus gen. nov., and the formal assignment of Nosocomiicoccus to the family Staphylococcaceae. Int. J. Syst. Evol. Microbiol. 70, 5926–5936 (2020).

47. Dong, P.-T. et al. Photolysis of Staphyloxanthin in Methicillin-Resistant Staphylococcus aureus Potentiates Killing by Reactive Oxygen Species. Adv. Sci. 6, 1900030 (2019).

48. Jain, C., Rodriguez-R, L. M., Phillippy, A. M., Konstantinidis, K. T. & Aluru, S. High throughput ANI analysis of 90K prokaryotic genomes reveals clear species boundaries. Nat. Commun. 9, 1–8 (2018).

49. Li, D., Liu, C.-M., Luo, R., Sadakane, K. & Lam, T.-W. MEGAHIT: an ultra-fast single-node solution for large and complex metagenomics assembly via succinct de Bruijn graph. Bioinformatics 31, 1674–1676 (2015).

50. Gande, R. et al. Acyl-CoA carboxylases (accD2 and accD3), together with a unique polyketide synthase (Cg-pks), are key to mycolic acid biosynthesis in Corynebacterianeae such as Corynebacterium glutamicum and Mycobacterium tuberculosis. J. Biol. Chem. 279, 44847–44857 (2004).

51. Portevin, D. et al. A polyketide synthase catalyzes the last condensation step of mycolic acid biosynthesis in mycobacteria and related organisms. Proc. Natl. Acad. Sci. U. S. A. 101, 314–319 (2004).

52. Chen, Y. et al. Structural classification and properties of ketoacyl synthases. Protein Sci. 20, 1659–1667 (2011).

53. Klaus, M., Buyachuihan, L. & Grininger, M. Ketosynthase Domain Constrains the Design of Polyketide Synthases. ACS Chem. Biol. 15, 2422–2432 (2020).

54. Medema, M. H., Cimermancic, P., Sali, A., Takano, E. & Fischbach, M. A. A systematic computational analysis of biosynthetic gene cluster evolution: lessons for engineering biosynthesis. PLoS Comput. Biol. 10, e1004016 (2014).

55. Lee, M. D. GToTree: a user-friendly workflow for phylogenomics. Bioinformatics 35, 4162–4164 (2019).

56. Garrison, E. et al. Variation graph toolkit improves read mapping by representing genetic variation in the reference. Nat. Biotechnol. 36, 875–879 (2018).

57. Kloos, W. E. & Schleifer, K. H. Isolation and Characterization of Staphylococci from Human Skin II. Descriptions of Four New Species: Staphylococcus warneri, Staphylococcus capitis, Staphylococcus hominis, and Staphylococcus simulans. Int. J. Syst. Bacteriol. 25, 62–79 (1975).

58. Becker, K., Heilmann, C. & Peters, G. Coagulase-negative staphylococci. Clin. Microbiol. Rev. 27, 870–926 (2014).

59. Liu, C.-I. et al. A cholesterol biosynthesis inhibitor blocks Staphylococcus aureus virulence. Science 319, 1391–1394 (2008).

60. Blount, Z. D., Barrick, J. E., Davidson, C. J. & Lenski, R. E. Genomic analysis of a key innovation in an experimental Escherichia coli population. Nature 489, 513–518 (2012).

61. Ghosh, S. & O’Connor, T. J. Beyond Paralogs: The Multiple Layers of Redundancy in Bacterial Pathogenesis. Front. Cell. Infect. Microbiol. 7, 467 (2017).

62. Otto, M. Staphylococcus epidermidis--the “accidental” pathogen. Nat. Rev. Microbiol. 7, 555–567 (2009).

63. Stokes, J. M. et al. A Deep Learning Approach to Antibiotic Discovery. Cell 180, 688–702.e13 (2020).

64. Melo, M. C. R., Maasch, J. R. M. A. & de la Fuente-Nunez, C. Accelerating antibiotic discovery through artificial intelligence. Commun Biol 4, 1050 (2021).

65. Carroll, L. M. et al. Accurate de novo identification of biosynthetic gene clusters with GECCO. bioRxiv 2021.05.03.442509 (2021) doi:10.1101/2021.05.03.442509.

66. Buchfink, B., Xie, C. & Huson, D. H. Fast and sensitive protein alignment using DIAMOND. Nat. Methods 12, 59–60 (2014).

67. Seemann, T. Prokka: rapid prokaryotic genome annotation. Bioinformatics 30, 2068–2069 (2014).

68. Parks, D. CompareM: A toolbox for comparative genomics. (Github).

69. Hyatt, D. et al. Prodigal: prokaryotic gene recognition and translation initiation site identification. BMC Bioinformatics 11, 119 (2010).

70. Aramaki, T. et al. KofamKOALA: KEGG Ortholog assignment based on profile HMM and adaptive score threshold. Bioinformatics 36, 2251–2252 (2019).

71. Tatusova, T. et al. NCBI prokaryotic genome annotation pipeline. Nucleic Acids Res. 44, 6614–6624 (2016).

72. Parks, D. H. et al. GTDB: an ongoing census of bacterial and archaeal diversity through a phylogenetically consistent, rank normalized and complete genome-based taxonomy. Nucleic Acids Res. (2021) doi:10.1093/nar/gkab776.

73. Chaumeil, P.-A., Mussig, A. J., Hugenholtz, P. & Parks, D. H. GTDB-Tk: a toolkit to classify genomes with the Genome Taxonomy Database. Bioinformatics 36, 1925–1927 (2019).

74. Ondov, B. D. et al. Mash: fast genome and metagenome distance estimation using MinHash. Genome Biol. 17, 132 (2016).

75. Chen, S., Zhou, Y., Chen, Y. & Gu, J. fastp: an ultra-fast all-in-one FASTQ preprocessor. Bioinformatics 34, i884–i890 (2018).

76. Wick, R. R., Judd, L. M., Gorrie, C. L. & Holt, K. E. Unicycler: Resolving bacterial genome assemblies from short and long sequencing reads. PLoS Comput. Biol. 13, e1005595 (2017).

77. Krueger, F., James, F., Ewels, P., Afyounian, E. & Schuster-Boeckler, B. FelixKrueger/TrimGalore: v0.6.7 - DOI via Zenodo. (2021). doi:10.5281/zenodo.5127899.

78. Walker, B. J. et al. Pilon: an integrated tool for comprehensive microbial variant detection and genome assembly improvement. PLoS One 9, e112963 (2014).

79. Li, H. Minimap2: pairwise alignment for nucleotide sequences. Bioinformatics 34, 3094–3100 (2018).

80. Li, H. & Durbin, R. Fast and accurate short read alignment with Burrows–Wheeler transform. Bioinformatics 25, 1754–1760 (2009).

81. Paradis, E., Claude, J. & Strimmer, K. APE: Analyses of Phylogenetics and Evolution in R language. Bioinformatics 20, 289–290 (2004).

82. Hug, L. A. et al. A new view of the tree of life. Nature Microbiology 1, 1–6 (2016).

83. Katoh, K. & Standley, D. M. MAFFT multiple sequence alignment software version 7: improvements in performance and usability. Mol. Biol. Evol. 30, 772–780 (2013).

84. Suyama, M., Torrents, D. & Bork, P. PAL2NAL: robust conversion of protein sequence alignments into the corresponding codon alignments. Nucleic Acids Res. 34, W609–12 (2006).

85. Stamatakis, A. RAxML version 8: a tool for phylogenetic analysis and post-analysis of large phylogenies. Bioinformatics 30, 1312–1313 (2014).

86. Kania, S. A. Reclassification of Staphylococcus schleiferi by Madhaiyan et al. lacks key supporting data. Int. J. Syst. Evol. Microbiol. 72, (2022).

87. Wood, D. E., Lu, J. & Langmead, B. Improved metagenomic analysis with Kraken 2. Genome Biol. 20, 1–13 (2019).

88. Capra, J. A. & Singh, M. Predicting functionally important residues from sequence conservation. Bioinformatics 23, 1875–1882 (2007).

