## Supplementary material for "Evolutionary investigations of the biosynthetic diversity in the skin microbiome using *lsa*BGC": Fig. S

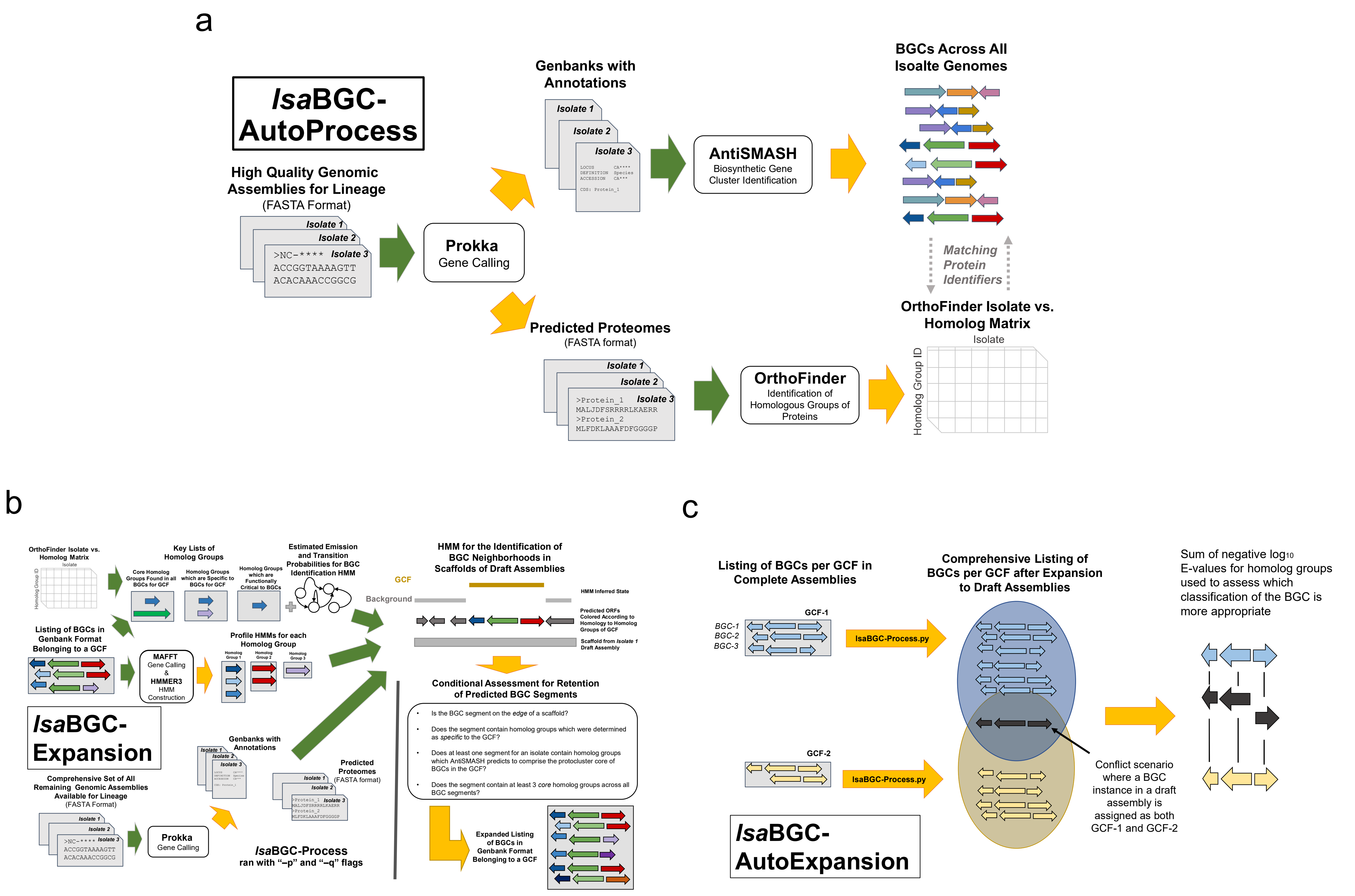


**Figure S1: Schematics of select IsaBGC programs and workflows.** a) A schematic of the *Isa*BGC-AutoProcess workflow which generates the inputs required for IsaBGC analyses. b) An overview of the algorithm behind *Isa*BGC-Expansion used to identify homologous instances of GCFs in a sensitive and efficient manner from potentially draft-quality assemblies. c) A schematic of the *Isa*BGC-AutoExpansion workflow which automatically runs *Isa*BGC-Expansion for each GCF and then resolves conflicts of overlap and consolidates results.

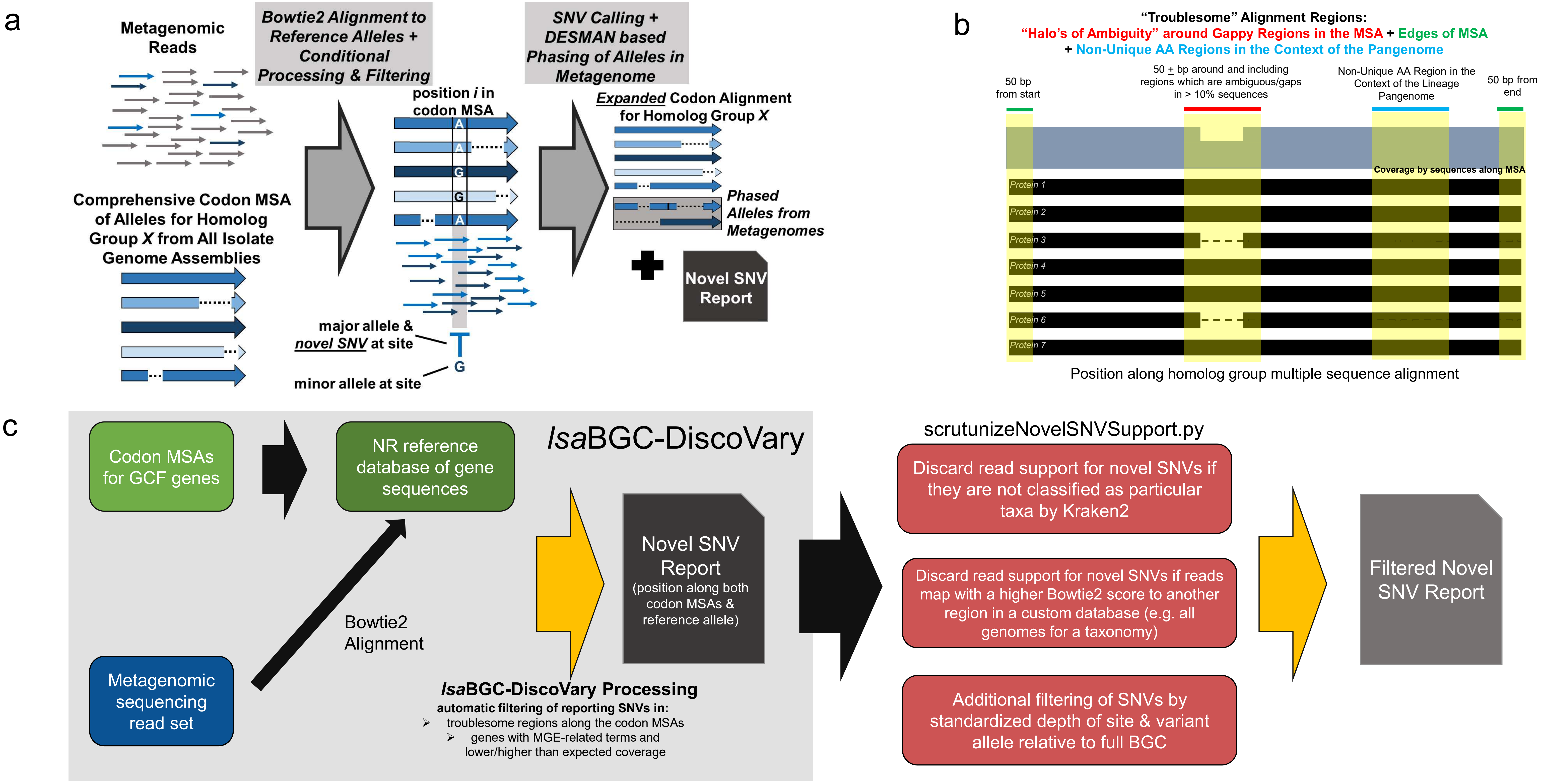


**Figure S2: An overview of *lsa*BGC-DiscoVary. a**) An overview of the *lsa*BGC-DiscoVary algorithm is shown. Codon alignments for each BGC homolog group, as constructed by *lsa*BGC-PopGene, are used to dereplicate and select representative alleles for mapping metagenomic readsets with Bowtie2. Afterwards, alignments are processed and used to identify whether homolog groups are represented in metagenomic samples. SNV sites along individual representative genes for homolog groups are mapped to comprehensive codon alignments and used to assess whether the allele represented by the SNV has previously been observed at the codon alignment position. Optional phasing of multiple alleles for a homolog group within a lineage or taxa can also be performed using DESMAN. **b**) Novel SNVs are not reported if they are within specific regions along codon alignments, including regions which are towards the beginning or end of the alignment, regions which are highly redundant, and regions where >10% of gene instances have deletions or lack sequence. **c**) An overview of *lsa*BGC-DiscoVary for determining putative novel SNVs which can then be further filtered using exhaustive methods to assert that reads supporting their presence do not map better to other taxa or genomic regions.

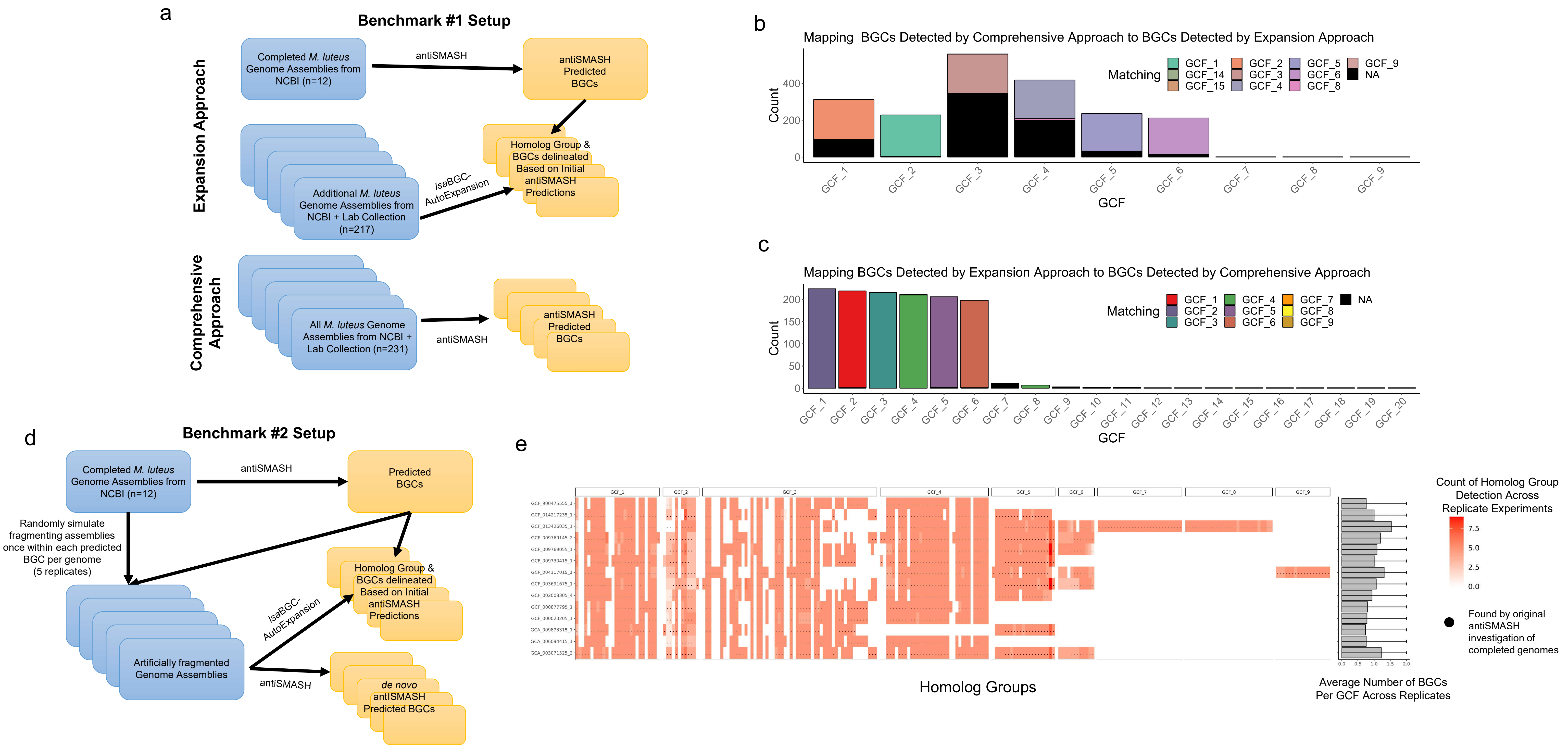


**Figure S3: Benchmarking *Isa*BGC-AutoExpansion using *M. luteus* genomes. a**) An overview of the first benchmarking experiment is shown comparing the use of IsaBGC-AutoExpansion to comprehensive antiSMASH to profile GCFs in 132 *M. luteus* genomes. **b**) Results from the first benchmarking experiment are shown. BGC instances identified by antiSMASH in genomes were mapped to BGC segments identified by running *Isa*BGC-AutoExpansion trained on GCFs from complete *M. luteus* genomes. "NA", shown in black, represent GCF segments which were undetected by antiSMASH. **c**) Results from the first benchmarking experiment are shown. BGC instances identified by *Isa*BGC-AutoExpansion were mapped to BGC segments identified by running *Isa*BGC-AutoExpansion. "NA" , shown in black, represent GCF segments which were undetected by *Isa*BGC-AutoExpansion. **d**) An overview of the second benchmarking experiment is shown comparing the use of IsaBGC-AutoExpansion to antiSMASH for predicting BGCs in 14 artificially fragmented complete genomes of *M. luteus*. **e**) Results from the second benchmarking experiment are shown. The heatmap shows whether homolog groups were identified by *Isa*BGC-AutoExpansion across 5 replicate simulations where assemblies were fragmented differently along BGC coordinates. Dots signify whether homolog groups were detected for a particular genome in the original antiSMASH annotation of the completed genome (unfragmented). The bar plot to the right shows that no more than two BGC segments were ever detected by *Isa*BGC-AutoExpansion as would be expected since each BGC was randomly fragmented into two pieces.

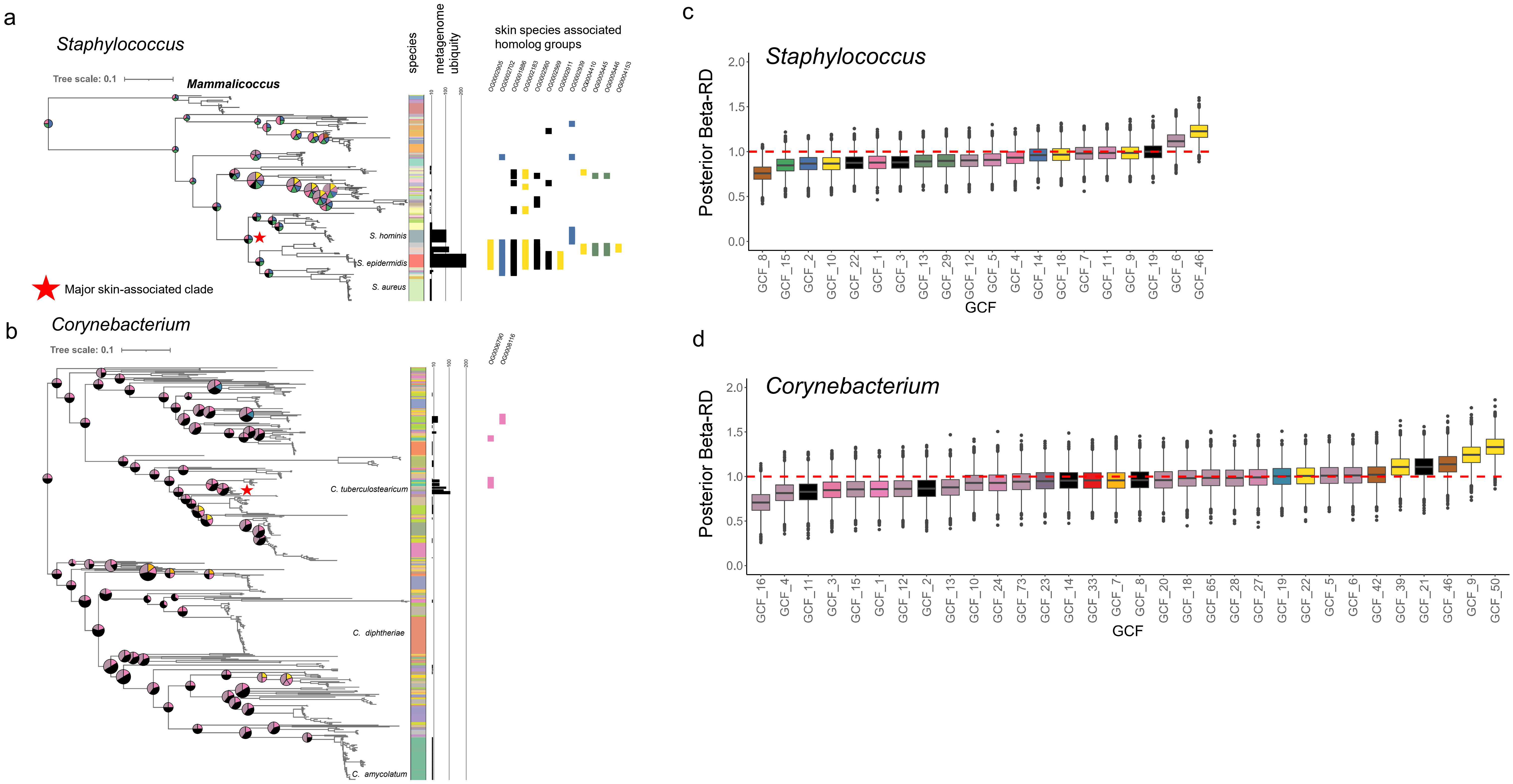


**Figure S4: Systematic identification of skin-associated BGC genes and signatures of intra-GCF horizontal gene transfer.** A maximum likelihood phylogeny was constructed from ribosomal protein encoding genes for **a**) 229 representative *Staphylococcus* genomes and **b**) 456 representative *Corynebacterium* genomes. Ancestral state reconstruction for GCF carriage was performed using maximum parsimony with the ACCTRAN algorithm and shown as piecharts for innernodes which encapsulate five or more species. Color strips to the right of the phylogenies correspond to the species classification of the genomes. The bar chart to the right of the species color strip corresponds to the number of metagenomes which were found to feature the species. The final track shows the presence of homolog groups found to be enriched in phylogenetic clades where $\geq$80% of genomes are classified as skin-associated species. IsaBGC-Divergence was used to calculate the Beta-RD statistic between pairs of genomes carrying a particular GCF in representative **c**) *Staphylococcus* and d) *Corynebacterium* genomes. Bayesian shrinking analysis was performed of raw Beta-RD calculates using STAN.

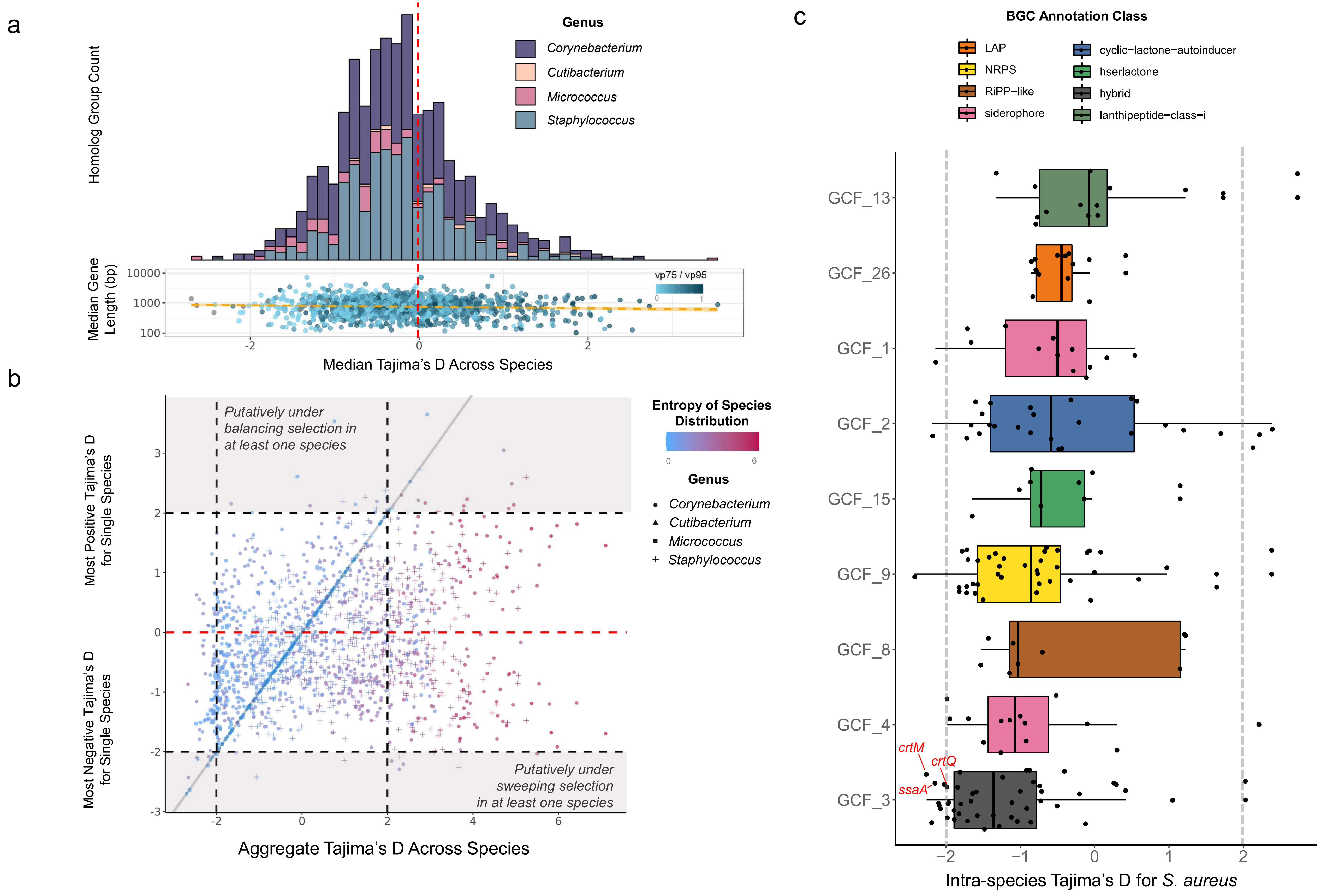


**Figure S5: Tajima's D statistic highlights GCFs and homolog groups under selective pressure. a**) For each homolog group found in a GCF, Tajima's D statistic was independently calculated per species and the median value across species was determined. The histogram of the median Tajima's D across species exhibits a normal distribution roughly centered around 0. The scatterplot below the histogram showcases that the median Tajima's D across species per homolog group exhibited no correlation with the median length of homolog groups. Each homolog group in the scatterplot is colored according to an analogous statistic to Tajima's D which is the ratio of sites along the homolog group's codon alignment where the major allele is found in ≥75% of sequences to sites where the major allele is found in ≥95% of sequences. **b**) The aggregate Tajima's D statistic was calculated over all sequences per homolog group and found to be biased by the number of species the homolog group was found in and thus the maximum and minimum Tajima's D per species was investigated instead. c) Intra-species Tajima's D calculations using 24 distinct *S. aureus* genomes highlight GCF-3, predicted to encode a hybrid terpene/T3PKS BGC, as feature seven homolog groups with Tajima's D below -2.0, including *crtM*, involved in staphyloxanthin biosynthesis, and *ssaA*, encoding a staphylococcal secretory antigen.

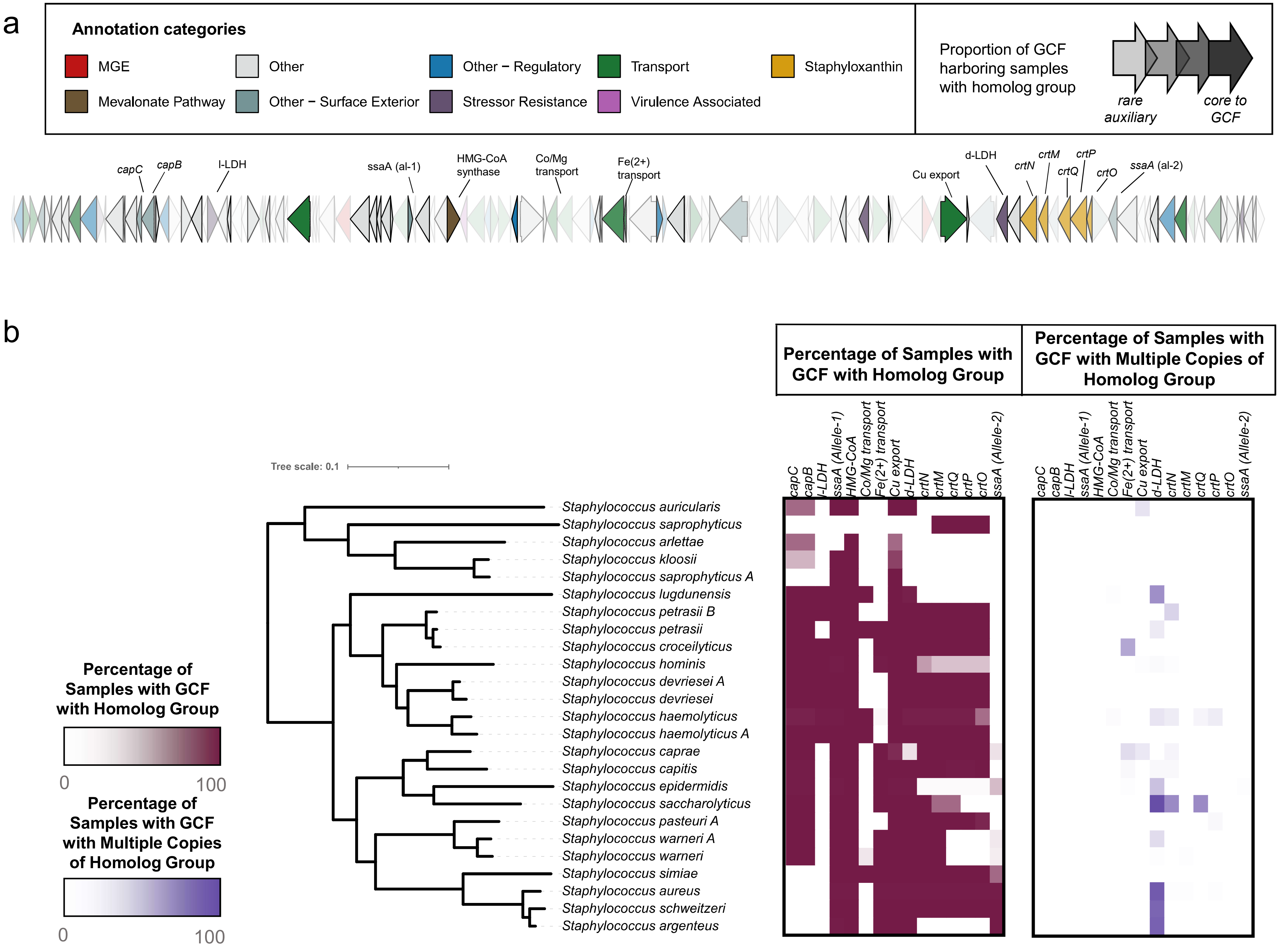


**Figure S6: Signatures of conservation and speciation in the surrounding context of staphyloxanthin encoding GCF-3.** GCF-3 is a predicted hybrid BGC which includes the staphyloxanthin encoding crt operon and the mevalonate pathway related enzyme hydroxymethylglutaryl-CoA synthase, a false positive detection due to homology with type-III polyketide synthases. **a**) A consensus schematic of GCF-3 is shown, generated from individual instances from 103 representative staphylococci found to feature it. Genes are colored according to broad annotation categories and transparency illustrates the proportion of GCF-3 carrying samples found to possess a specific homolog group. **b**) A maximum-likelihood phylogeny for the 25 species within the *S. aureus* / *S. epidermidis* clade found to carry GCF-3 is shown alongside three heatmaps depicting species-specific metrics for select homolog groups. The left heatmap showcases the percentage of the total species genomes found to carry select genes from GCF-3, including the staphyloxanthin encoding *crt* genes. The right heatmap depicts whether homolog groups are found in multiple-copies within the GCF-3 context for different species.

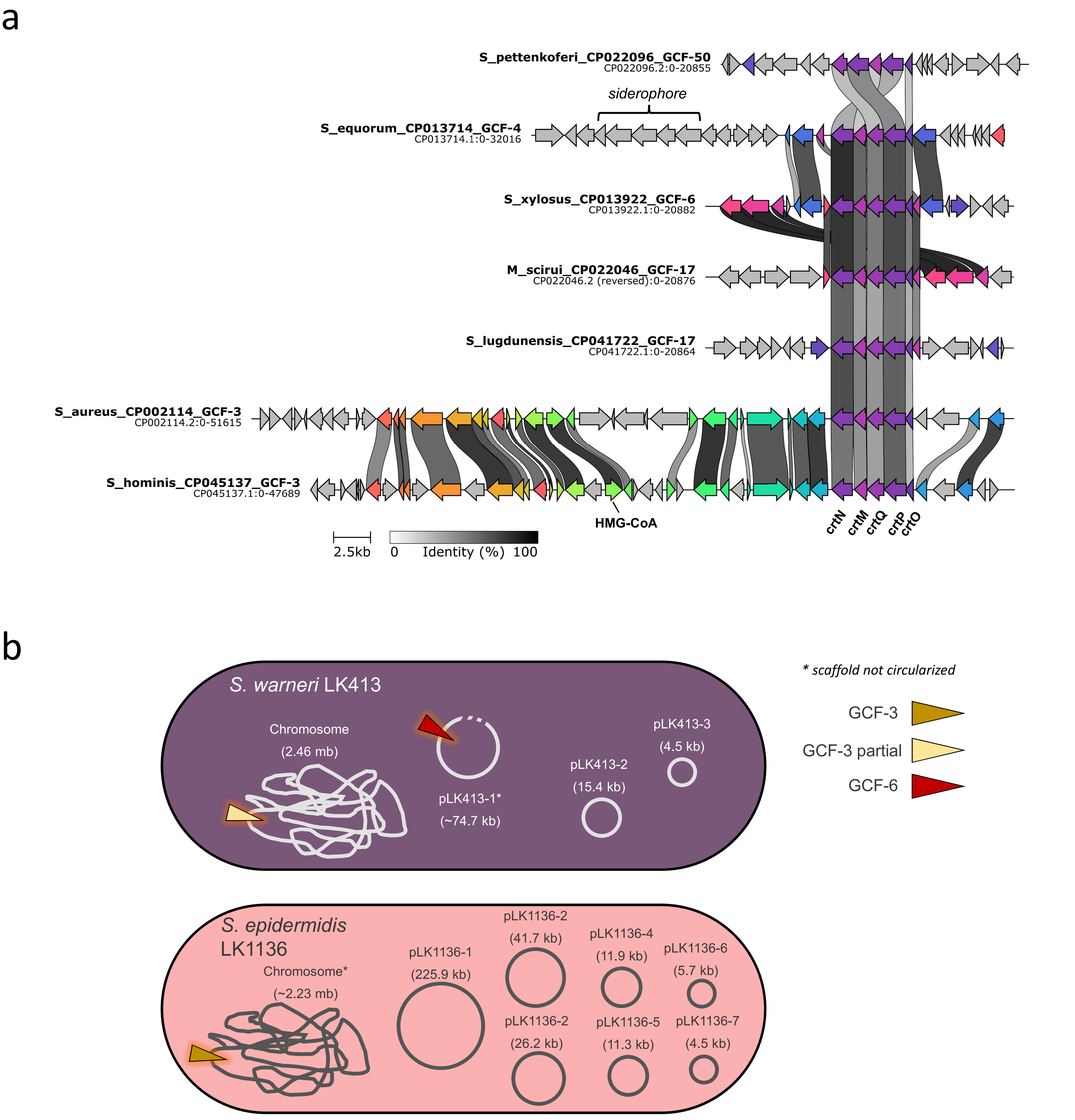


**Figure S7: The staphyloxanthin encoding crt operon is found in multiple GCFs. a**) Sequence and syntenic comparisons between representative instances of GCFs featuring the crt operon encoding for staphyloxanthin were performed and illustrated using clinker. Based on phylogenetic analysis of CrtMN sequences (Figure 3b), the *crt* operon is likely misclassified as GCF-17 for *S. lugdunensis* and should instead be GCF-3. The cause of the misclassification is because the *crt* operon is in a different genomic context within the species as compared to other species in the *S. aureus/epidermidis* clade with GCF-3. **b**) Schematics of the near completed genomic assemblies for *S. epidermidis* LK1136 and *S. warneri* LK413 isolated from skin showcasing the location of staphyloxanthin encoding GCFs.

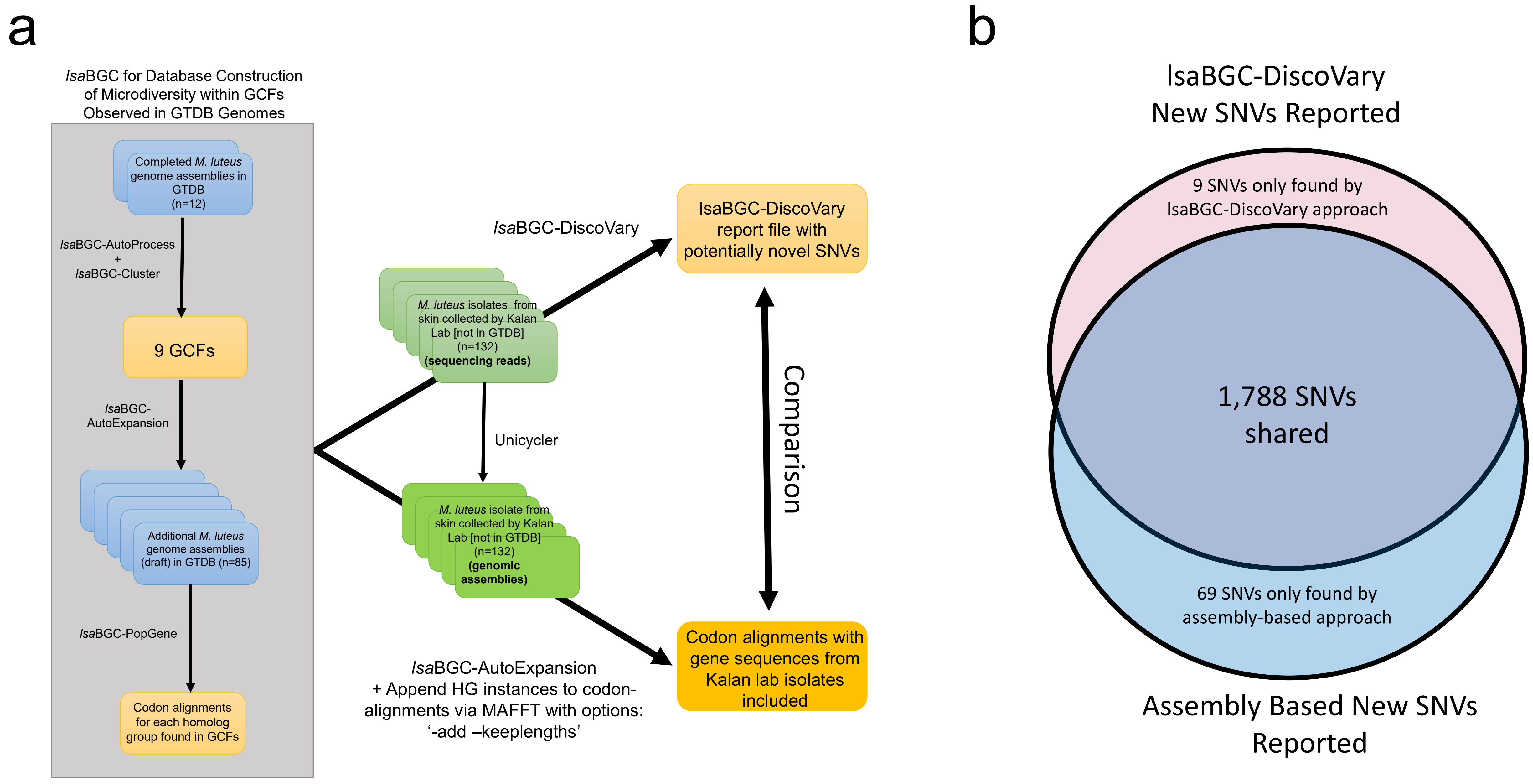


**Figure S8: Benchmarking *Isa*BGC-DiscoVary to assembly based SNV identification using single isolate genomic sequencing data for 132 *M. luteus*. a**) A schematic of the benchmarking setup for comparing *Isa*BGC-DiscoVary identification of novel SNVs from sequencing reads for 132 *M. luteus* compared to an assembly based identification of novel SNVs for the same isolates. Novel SNVs corresponded to alleles which were not previously represented at specific sites in homolog group codon alignments constructed from publicly available *M. luteus* genomes gathered from NCBI. **b**) A venn diagram showcasing the number of novel SNVs reported by *Isa*BGC-DiscoVary as compared to the novel SNVs found by the assembly-based approach.

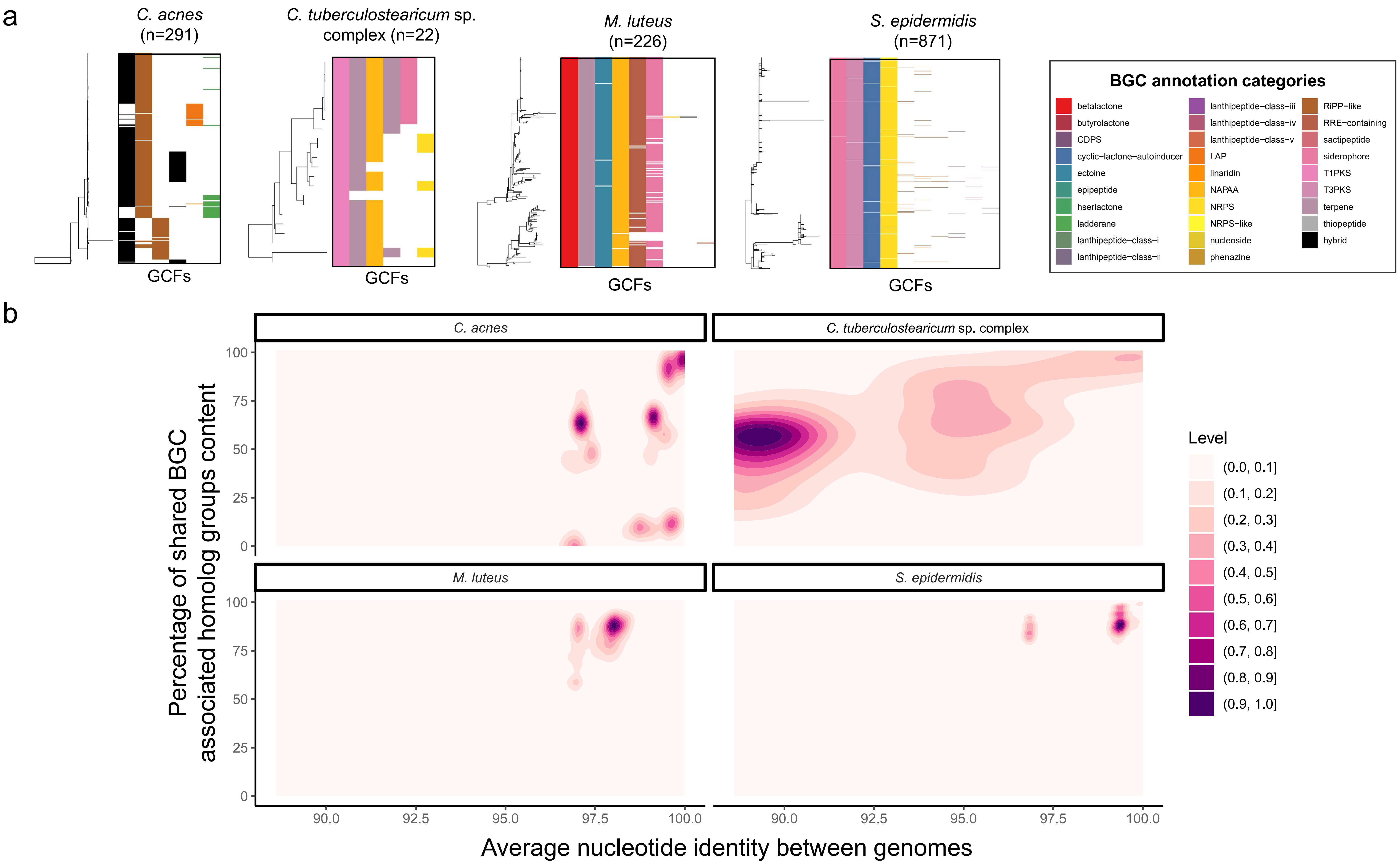


**Figure S9: Highly similar *C. acnes* isolates can have vastly divergent BGC content. a**) The presence of GCFs is shown across phylogenies constructed from genes encoding for ribosomal proteins for the species *C. acnes*, *M. luteus*, and *S. epidermidis* and the *C. tuberculostearicum* species complex. **b**) For each species, the genome-wide average nucleotide identity (ANI) between pairs of genomes was compared to the number of BGC encoded homolog groups they share.

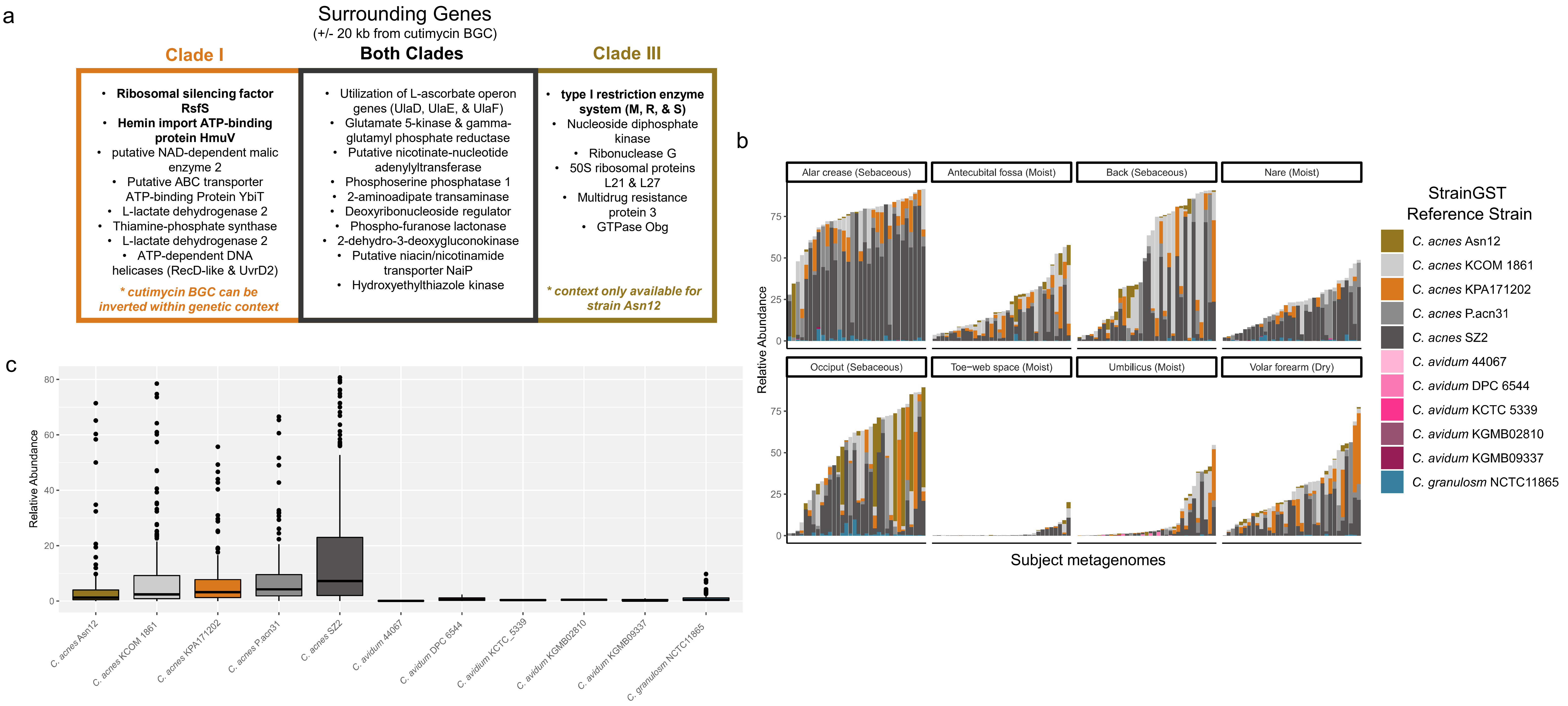


**Figure S10: The distribution of cutimycin encoding *C. acnes* across skin metagenomes. a**) The surrounding context of the cutimycin encoding BGC featured many genes in common between clade IB and clade III *C. acnes* as well as certain genes that were specific to each clade. **b**) The relative abundance of *Cutibacterium* and representative strains of the genus are shown across skin metagenomes from different body sites and individuals. **c**) Strains representative of clade IB and clade III *C. acnes*, which encode cutimycin, are less common in skin metagenomes compared to two other representative *C. acnes* strains (different subclades of clade I).

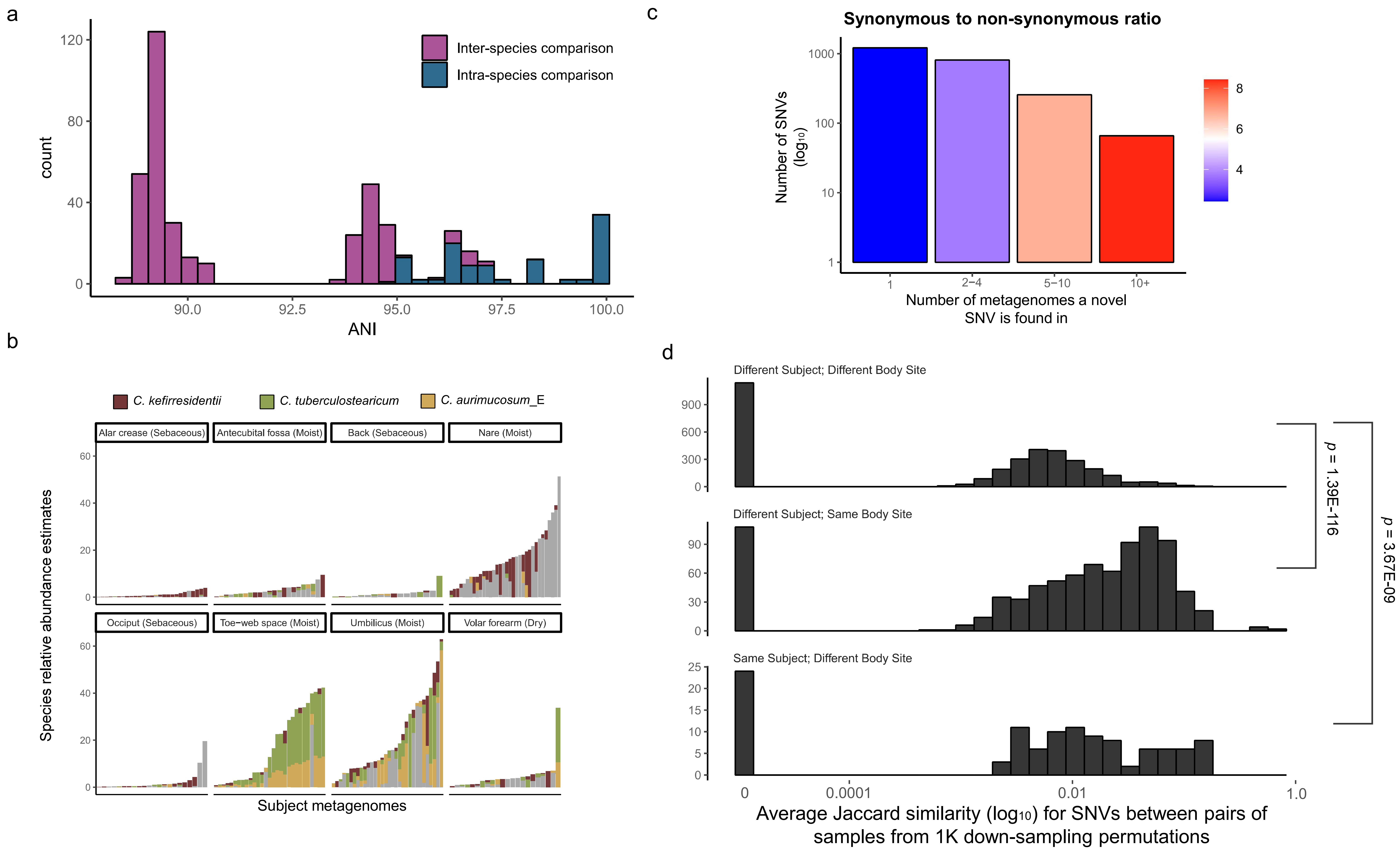


**Figure S11: The *C. tuberculostearicum* species complex is highly prevalent across skin metagenomes. a**) The average nucleotide identity between 22 genomes in the C. *tuberculostearicum* species complex is shown with coloring representing intra-species vs. inter-species comparisons. **b**) The relative abundance of *Corynebacterium* and representative strains of the *C. tuberculostearicum* species complex in the StrainGST database are shown across skin metagenomes from different body sites and individuals. **c**) Novel SNVs were tabulated by the number of metagenomes they were observed in. The color of bars corresponds to the ratio of suspected synonymous to non-synonymous SNVs for the sets of novel SNVs. **d**) For metagenomic samples where 30 or more novel SNVs were identified, a multi-iteration, down-sampling based approach was used to compute the average Jaccard similarity for number of shared novel SNVs between pairs of metagenomes. Pairwise comparisons of metagenomes were categorized by whether samples were from the same body-site or subject.

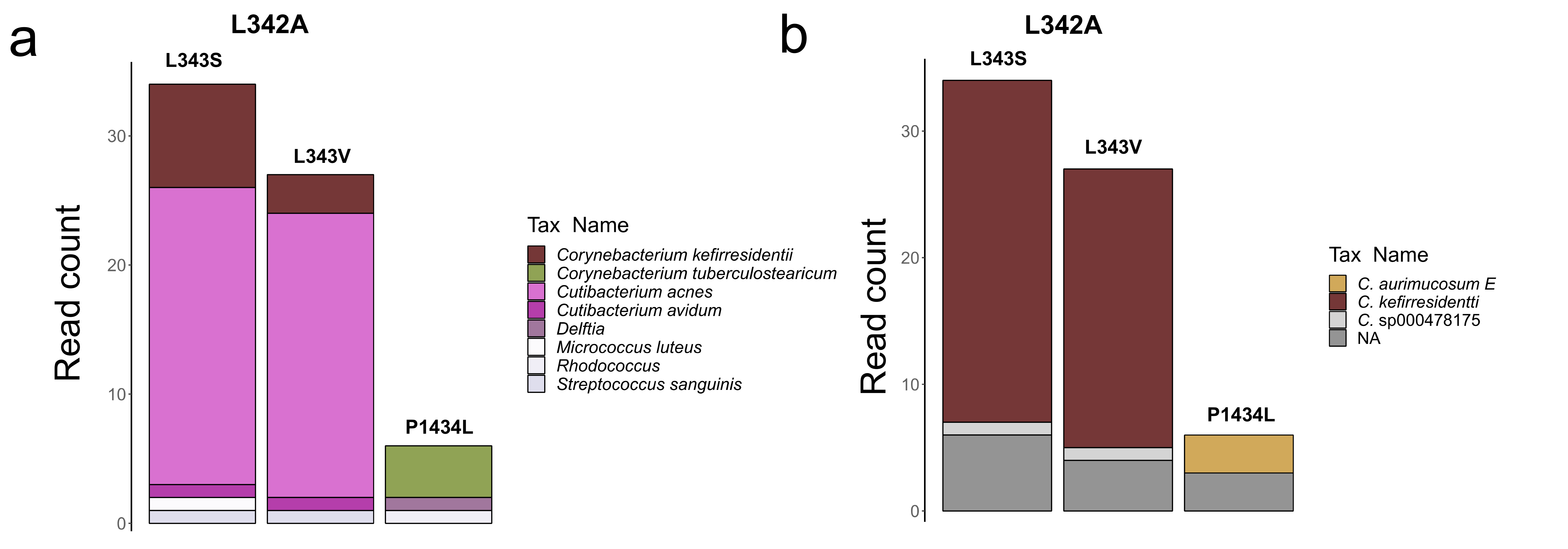


**Figure S12: Validation of taxonomic origin for reads supporting the presence of non-synonymous SNVs at highly conserved sites as identified by *lsa*BGC-DiscoVary.** a) Reads supporting the existence of three novel SNVs in highly conserved sites of the mycolic acid biosynthesis polyketide synthase (*HG0001691*) were classified taxonomically with Kraken2. Despite many of the reads supporting novel SNV existence being classified as *Cutibacterium*, paired-end alignment to a comprehensive database of all *Cutibacterium* genomes featured in GTDB R202 showed that none aligned concordantly. **b**) Of the reads supporting the existence of novel SNVs, 85.07% aligned as concordant pairs to a comprehensive database of all *Corynebacterium* genomes, with the majority aligning to species belonging to the *C. tuberculostearicum* species complex.
