## Supplementary Text for "Evolutionary investigations of the biosynthetic diversity in the skin microbiome using *lsa*BGC"

#### Updates to *lsa*BGC since release 1.0

Changes to lsaBGC since its initial release have primarily focused on simplifying usage of the suite through development of lsaBGC-Ready.py and lsaBGC-Easy.py. Analytical changes have mostly been minimal but include: (i) updated formatting of result files from the lsaBGC-AutoAnalyze.py workflow, (ii) changing how gaps are accounted for in calculating the Beta-RD statistic from codon-alignments, (iii) an adjustment to key-word searches used to prevent lsaBGC-DiscoVary from calling variants upon genes for which annotation suggests are MGEs, (iv) using MAGUS[89] in place of MAFFT[83] for protein alignment, to allow for better scalability, and (v) introducing a more stringent requirement for homolog groups regarded as part of the protocore of BGC predictions by antiSMASH[8]. Incorporation of GToTree[55] also provided the opportunity to efficiently and more easily estimate expected similarities between genomes using protein alignments of single copy genes used by the software to construct phylogenies. As such, we have removed support for inferring Beta-RD using nucleotide similarity between genomes. We have also added support for GECCO[65] and DeepBGC[11] predictions of BGCs. Finally, we have begun to introduce code to enable application of *lsa*BGC to fungi and plants but are still testing these functionalities.

#### Assessment of hybrid genomic assemblies constructed for *S. epidermidids* LK1136 and *S. warneri* LK413

We chose to investigate staphyloxanthin production in *S. epidermidis* and *S. warneri* because they are *Staphylococcus* species commonly isolated from skin. Specifically, the former species *S. epidermidis*, is the most abundant staphylococcal species on skin and the latter species, *S. warneri*, featured two distinct staphyloxanthin encoding GCFs. For the *S. epidermidis* LK1136 hybrid ONT and Illumina genome assembly, we found that the isolate featured eight plasmids, including a 225 kb mega-plasmid (Table S7b). All eight plasmids were regarded as circularized; however, the chromosome was not. For the *S. warneri* LK413 hybrid genome assembly, we found that the isolate featured three plasmids, of which two were circularized (Table S7b). The chromosome for this isolate was circularized and regarded as complete. The incomplete scaffold in *S. warneri* was found to feature staphyloxanthin encoding GCF-6 and to validate it represented a plasmid, we used the BLASTn[90] to NCBI's nt analysis performed in GAEMR(https://software.broadinstitute.org/software/gaemr/) to confirm that all reported alignments were to *Staphylococcus* plasmid sequences. Additionally, this scaffold featured a slightly elevated coverage, ~3X greater than the chromosome.

#### Genome annotation, homology determination in predicted proteomes, and clustering of BGCs into GCFs

Complete or chromosome level assemblies were run through lsaBGC-AutoProcess.py which invokes Prokka[67] for gene calling and annotation, antiSMASH for BGC detection (v6)[8], and OrthoFinder2[13] for delineating homologous clusters of proteins (Figure S1a).

Subsequent clustering of detected BGCs into GCFs was performed using lsaBGC-Cluster.py. For consistency and simplicity, we ran both the genus-level and species-level analyses using identical parameters for lsaBGC-Cluster.py. The MCL inflation parameter was set to 4.0, the minimal syntenic similarity threshold was set to 0.7, and the Jaccard similarity of homolog groups shared vs. observed in union between two BGCs threshold was set to 20.

lsaBGC-Cluster.py leverages homology information from OrthoFinder2[13] together with syntenic similarity akin to the approach taken in BiG-SCAPE[9]. However, unlike BiG-SCAPE, which is reliant on domains, lsaBGC-Cluster.py uses entire protein sequences designated to discrete homolog groups, offers a tailored report for users on appropriate parameter selection, and employs Markov chain clustering (MCL)[91] instead of affinity propagation for granular GCF delineation. Similar to BiG-SCAPE, lsaBGC-Cluster.py also uses both the presence of coding units (domains for BiG-SCAPE; homolog groups for lsaBGC-Cluster.py) as well as sequence similarity between such units to appropriately cluster BGCs. In lsaBGC-Cluster.py, the presence of homolog groups and their sequence similarity is implicit to the OrthoFinder2 algorithm which determines appropriate thresholds for designating homologous, ideally orthologous, protein instances by accounting for genome-wide similarity[13]. Also, similar to BiG-SCAPE in concept[9], lsaBGC-Cluster.py can be set to require syntenic similarity between BGC instances for clustering, but the methodology in lsaBGC-Cluster.py is based on global syntenic similarity measured using the absolute value of the Spearman correlation for sets of three homolog groups shared between BGC instances. BiG-SCAPE on the contrary uses more localized information on adjacent-pairs of domains to infer overall syntenic similarity. Finally, both lsaBGC-Cluster.py and BiG-SCAPE offer flexibility for whether to perform preliminary partitioning of BGCs based on the predicted class by antiSMASH. By default, lsaBGC-Cluster.py does not perform such partitioning.

Unique to lsaBGC-Cluster.py is an option to produce a user-friendly PDF report guiding users for optimal parameters for clustering. This report showcases how different values of the MCL inflation parameter and the Jaccard similarity threshold for shared homologs between pairs of BGCs affects the final clustering. In this report, users can see how different parameter combinations can influence the number of singleton GCFs, GCFs with a single BGC member, the number of the core homolog groups observed in all BGCs belonging to a particular GCF (Table S2, S3). Such a report is critical for users to be able to move forward with an appropriate delineation of GCFs and not to re-track and manually experiment with different clustering configurations. An example report, pertaining to clustering BGCs from 77 *Staphylococcus* genomes with completed or chromosome-level assemblies, as well as descriptions for each of the figures featured in its 52 pages can be found on the Github wiki page for the lsaBGC-Cluster.py program.

To additionally provide users with versatility to better define the boundaries of a GCF, which could be particularly useful when dealing with hybrid BGCs or multiple BGCs co-located nearby each other, we further provide lsaBGC-Refiner.py. lsaBGC-Refiner.py takes in as input a GCF listing file of BGC instances belonging to it and a pair of user-defined boundary homolog groups. It then filters BGC Genbanks to retain only genes found in between the two boundary homolog groups. This functionality is particularly useful for instances where antiSMASH is inconsistent in defining hybrid BGCs due to variable inter-protocore content which has recently been highlighted[7].

For biosynthetic class designations of GCFs in this study, we regarded GCFs as a single class if at least 90% of BGC instances from complete genomes were predicted by antiSMASH as encoding for the class; otherwise, the GCF was regarded as a hybrid.

### High throughput identification of homologous instances of GCFs in assemblies

To mine metagenomic datasets for base-resolution novelty within BGCs previously unobserved from available assemblies for a given taxa, it is first necessary to comprehensively profile all allelic variants of a homolog group from such a set of assemblies. We thus developed lsaBGC-Expansion.py to efficiently and systematically identify orthologous instances of GCFs across the comprehensive set of assemblies available for a species, which for certain *Staphylococcus* species could range in the thousands on NCBI's Genbank database (Figure 1cd, S1b). In concept, lsaBGC-Expansion.py is similar to the now common bioinformatics practice of defining homolog groups for proteins upfront and then searching for their presence in new genomes[70,92,93]. In fact, re-identifying instances of homolog groups found within GCFs is the first component of lsaBGC-Expansion.py. This is performed by first constructing profile HMMs for such homolog groups using MAFFT local alignment[83] with standard settings and HMMER3[94], followed by searching genomic assemblies directly using HMMER3 or emitting the consensus sequence and searching via DIAMOND[66], as inspired by the approach developed by Melnyk et al. 2019[92]. For the analyses presented in this paper, we used the DIAMOND-based approach (specified by --quick_mode argument).

In a preliminary step, profile HMMs are similarly aligned to the predicted proteome of genomes from the initial set of genomes which were run through antiSMASH and used to establish GCFs. This reflexive alignment enables the determination of appropriate E-value thresholds to gauge the presence or absence of homolog groups in new genomes. Specifically, the E-value thresholds for each homolog group is determined as either: (i) the lowest E-value of false positive alignments multiplied by a factor of 1E-5 or (ii) a default threshold of 1E-10 if E-values for some true positive alignments are found to be higher than those for false positive alignments. Importantly, this methodology for selecting E-value thresholds assumes that new genomes being searched are phylogenetically interspersed with genomes used for initial analysis and construction of the profile HMMs. We additionally required proteins which aligned to homolog group profile HMMs to also be of similar length to the proteins used to construct the profile HMMs, requiring them to be at most 1.5X the max(25, median absolute divergence) base-pairs shorter or longer than the median length of the representative proteins of the homolog group.

A classical HMM framework, leveraging the pomegranate library[95], is used to scan predicted coding genes across assembly scaffolds, where coding genes are regarded to follow a binary state based on whether they exhibit homology to a GCF associated homolog group. Our approach mimics the algorithm of ClusterFinder[10], but features several important differences to identify fragmented instances matching well characterized BGCs rather than search for novel BGCs with remote homology to known instances. We estimate the emission probability of each homolog group based on whether it's profile HMM is able to distinguish true hits from false hits from reflexive alignments (see previous paragraph). If it is able to distinguish true-hits from false-hits, then detecting a homolog group at the E-value threshold described above is very likely to represent a predicted protein belongs to the GCF and the homolog group will have an emission probability of 0.99 for the "GCF State" and a emission probability of 0.01 for the "Background State". If the profile HMM is unable to distinguish true-hits from false-hits, then the emission probability for the "Background State" is set to a maximum of either: (i) 1.0 - (# of GCF instances / # of total instances) or (ii) 0.2. The emission probability for the "GCF State" in this case will be set to the complement of the "Background State" probability. The transition probabilities between states are set to 0.1 by default and to 0.9 for transitioning to the same state. The start and end of a scaffold are equally likely to correspond to either the states of GCF or Background.

Genomic neighborhoods of predicted coding genes detected by the HMM as corresponding to the GCF (herein referred to as "potential GCF segments") are next conditionally assessed to avoid reporting false positive neighborhoods which are likely not related to the GCF in question. Each potential GCF segment must feature at least 3 homolog groups found in a known instance of the GCF. Additionally, all potential GCF segments must display reasonable syntenic similarity (gene positioning) to a known instance of the GCF found in the high quality genomic assemblies. Unlike the syntenic similarity filtering applied in the clustering of BGCs into GCFs, here we use Pearson's correlation instead of Spearman's correlation to more reliably assess syntenic similarity of shorter segments to known GCF instances. Correlations are only calculated if the genes in the segment under consideration and the comparing known BGC instance have genes in the same relative direction. A default correlation of 0.8 to at least one known GCF instance is required by each segment (with p-value < 0.1).

Segments are reported as part of the GCF automatically if any of the following criteria are met: (i) The segment features >= 5 homolog groups and segment features >= 3 "core" homolog groups ("core" homolog groups are those which are observed in all known instances of a GCF from the high-quality genomic assemblies), (ii) the segment features a GCF "specific" homolog group (these are homolog groups which are only observed within the GCF in the high-quality genome assemblies, thus their presence is a good indication the segment belongs to the GCF and is not a false positive), or (iii) the segment features homolog groups which overlap with the core of protoclusters in GCFs as delineated by antiSMASH. If segments do not meet any of the three criteria above, they can still be reported if they are on the edge of scaffolds (within 500 bp from the end of a scaffold). These segments do not need to meet criteria (i) individually, but do need to feature >=5 homolog groups and >=3 core homolog groups of the GCF in unison. Additionally, only one edge segment per scaffold is allowed.

To ensure we are not missing more evolutionarily diverged variants of homolog groups not represented in the initial lsaBGC analysis, we perform a final "polishing" step for segments which are considered to belong to a GCF. This involves reassessing genes on GCF segments which are unassigned to any homolog group and additionally searching for surrounding genes (up to 10 genes on each side) which display homology to GCF associated homolog groups. A gene only needs to display homology at a level of less than 1E-10 E-value to be assigned to a homolog group. Critically, we only assign these genes to homolog groups after the segment is determined as belonging to the focal GCF to ensure our detection of segments is based on more concrete/significant homology-based evidence

Ultimately, we do not recommend running lsaBGC-Expansion.py individually on a GCF-by-GCF basis, but to instead use the wrapper workflow program lsaBGC-AutoExpansion.py, to automatically search for homologous instances across all GCFs. This is because lsaBGC-AutoExpansion.py is able to resolve potential conflicts where some part of a genomic assembly might independently be assigned to two separate GCFs (Figure S1c). After running lsaBGC-Expansion.py individually per GCF, it consolidates results and resolves any conflict in genes from the same genomic assembly being assigned to multiple GCFs. It resolves such conflicts by performing pairwise comparisons of the gene sets across every BGC instance from every GCF. Overlap between two BGCs from distinct GCFs is considered a conflict if they overlap with more than 5% of the number genes of one of the BGCs. If this is the case, the sum of the exponents of E-values of each BGC's genes to their respective homolog group profile HMMs are compared and the BGC with the lower sum (indicating more genes in the BGC match a GCF profile or that genes match better to the GCF profile) is retained while the other is discarded. lsaBGC-AutoExpansion.py also creates consolidated result files, including: (i) an updated/expanded sample vs. homolog group gene matrix, (ii) an updated/expanded sample listings file (sample Genbank assembly file predicted proteome file), and (iii) an updated listing files of BGCs for each GCF.

*Rapid determination of 63 GCFs in >15K* Staphylococcus *genomes*: We were able to use lsaBGC-AutoExpansion.py to rapidly identify GCFs determined from complete *Staphylococcus* genomes in all ~15K *Staphylococcus* genomes represented in GTDB release R202. As expected, we found that fewer homolog groups in BGCs were detected for low quality assemblies (N50 < 10K; median of 105 homolog groups in BGCs per genome) as compared to high-quality assemblies (N50 > 100K; median of 164 homolog groups in BGCs per genome) ($p$=8.58E-36; two-sided Wilcoxon rank sum test). Furthermore, we found a lower percentage of homolog groups found together on the same BGC fragment with core biosynthetic machinery for low quality assemblies (N50 < 10K; median of 54%) as compared to high-quality assemblies (N50 > 100K; median of 100%) ($p$=8.64E-39; two-sided Wilcoxon rank sum test) (Figure 1c). Note, the time taken to run lsaBGC-AutoExpansion.py on all draft-quality *Staphylococcus* genomes described in the main text is not inclusive of preliminary gene-calling using Prokka[67] performed via lsaBGC-AutoProcess.py.

M. luteus - *Benchmarking comprehensive antiSMASH vs. lsaBGC framework*: To benchmark the sensitivity and specificity of the *lsa*BGC framework, whereby we perform antiSMASH[8] based identification on a subset of complete genomes for a taxa and then identify homologous instances in draft quality genomes, against simply running antiSMASH on all the available genomes for the taxa, we performed two comparative analyses using *M. luteus* genomes (Figure S3). Because lsaBGC-Expansion.py accounts for all homolog groups found in a GCF rather than being dependent on a smaller subset of BGC associated domains, lsaBGC-Expansion.py should have increased sensitivity for detection of genomic regions belonging to a GCF compared to antiSMASH. For instance, if in truth a BGC exists across two separate scaffolds, but only one segment features all the core domains encoding for the secondary metabolite biosynthesis machinery, then a rule-based approach seeking specific domains might struggle with detection of the second segment consisting of only auxiliary cargo, even if some of those genes encode enzymes critical for the production of the final metabolite.

In the first experiment, to assess the sensitivity of lsaBGC-Expansion.py, we compared the final GCFs of two different lsaBGC analyses: (i) an expansion based analysis where we ran initial processing, antiSMASH based BGC identification, and GCF clustering for 14 high-quality "complete" / "chromosome" quality *M. luteus* assemblies and then perform expansion of the GCFs with 213 additional *M. luteus* draft genomic assemblies of lower quality, and (ii) a comprehensive clustering analysis where we ran initial processing, antiSMASH based BGC identification, and GCF clustering across all 227 genomic assemblies in consideration (Figure S3a).

This comparative analysis essentially allowed us to assess how many instances (in this case contiguous segments) of a particular GCF are missed by the *lsa*BGC expansion-based framework (Figure S3c). We found that almost all instances of BGCs found when running antiSMASH on the draft assemblies directly, were also identified through lsaBGC-AutoExpansion.py using BGCs from the 14 high-quality genomic assemblies as references (97.8%; 1206/1233). Nearly all of the exceptions  not detected in the expansion based analysis (88.9%; 24 of 27) corresponded to rare GCFs which had no representatives in the high-quality assemblies. These cases demonstrate a significant limitation of the *lsa*BGC core framework in which BGCs are identified *de novo* in only a subset of genomic assemblies and is an important consideration for users.

The analysis also allowed us to assess how many instances of a particular GCF were only found by lsaBGC-AutoExpansion.py and not by running antiSMASH directly on draft genomes (Figure S3b). Given that our approach is a taxa-focused analysis and utilizes all homolog groups associated with a GCF, not just those containing core BGC domains, it is not surprising that we were able to find substantially more instances of BGC segments in the draft assemblies using lsaBGC-AutoExpansion.py which were missed by antiSMASH. This is likely because of assembly fragmentation, as described in the example scenario above. In total, we found 689 new GCF instances only through lsaBGC-Expansion. Of these, most (87.1%, 600 of 689) correspond to true "expansions" (the GCF was detected as present in the samples in the comprehensive clustering analysis using antiSMASH, but additional segments containing auxiliary content on different scaffolds of the assembly were only detected by lsaBGC-AutoExpansion.py).

In the second benchmarking experiment, we again use the 14 completed *M. luteus* genomes available, but in this setup our aim was to compare how the *lsa*BGC approach compared to running *de novo* antiSMASH when these genomes were artificially fragmented via simulation (Figure S3d). This benchmarking setup allowed us to assess how many coding genes initially identified as part of distinct GCFs on complete genomes are re-identified after genomes are artificially fragmented. We performed *in silico* simulation of assembly fragmentation for each of the 14 genomes in five replicates. For each such replicate simulation, we randomly fragmented genomic assemblies at a single point within each BGC predicted for it by antiSMASH. This resulted in 70 fragmented assemblies (5 for each of the 14 genomes), which were then searched *de novo* for BGCs using antiSMASH or searched based on similarity to the initial BGCs identified in the completed (unfragmented) genomes using lsaBGC-AutoExpansion.py. Coding genes were matched between the initial completed genomes with their fragmented versions based on exact sequence matching and also checking if ORFs from the fragmented genome were subsequences of ORFs from the complete genomes.

As expected based on the algorithms employed, we found that lsaBGC-AutoExpansion.py was able to re-identify more of the initial coding genes associated with BGCs in complete genomes compared to rerunning antiSMASH *de novo* (Figure S3e). antiSMASH detected only 64.7% on average of the homolog groups it had originally associated with BGCs in complete genomes from the artificially fragmented assemblies, while lsaBGC-AutoExpansion.py was able to recover 95.4% of them on average. Despite this increased sensitivity, lsaBGC-AutoExpansion.py never predicted more than 2 BGCs per GCF for any of the 70 fragmented genomes, which, because GCFs were split in two, would indicate that it was overpredicting BGC instances and incurring false positives (Figure S3e). However, lsaBGC-AutoExpansion.py did identify multiple instances of homolog groups which were not present in the initial BGC predictions on complete genomes. This is because lsaBGC-AutoExpansion.py used as reference all BGC instances from across the 14 genomes to identify GCFs in the fragmented genomes and thus increased the boundary of GCFs beyond what they might have initially been set to by the initial antiSMASH run on the complete (unfragmented) genomes. This can be seen as another key advantage of lsaBGC-AutoExpansion.py, in that it allows smoothing of BGC boundaries upon identification and can enable more comprehensive and robust comparative genomics analytics downstream.

### Visualization of BGCs across phylogenies

We have provided the lsaBGC-See.py program to allow users to visualize GCF instances across either a user-provided species phylogeny or automatically generated GCF-specific phylogeny. While the latter functionality is present in CORASON[9], the former feature, which can provide a differing and important evolutionary perspective, is currently unique to *lsa*BGC. *lsa*BGC creates automated PDF reports using R visualization libraries ggplot2[96] and ggtree[97] as well as creates a track for visualization in iTol[98]. Additionally, the constructed GCF phylogeny and user-provided species phylogeny are reformatted to a Newick file in which genome identifiers are expanded if multiple instances of a GCF are found for it, which is critical for visualizing fragmented BGCs from draft genomes. The program generates a GCF specific phylogeny if requested by creating codon alignments (as described in the section "Understanding GCF Conservation and Composition through Evolutionary and Population Genetic Statistics"), concatenating these, filtering for conserved SNVs and generating an approximate maximum-likelihood phylogeny using FastTree2[99].

For comparative visualization between staphyloxanthin encoding GCF-6 instances in the chromosome and on a plasmid in *S. equorum* C2014 as well as between different staphyloxanthin encoding GCF representatives (Figure 3d, S7a) we used clinker[100]. For visualization of staphyloxanthin encoding GCFs across the *Staphylococcus* phylogeny (Figure 3a, S6b), only a single representative genome from each species, as classified by GTDB[72], was selected and used to prune the original ribosomal phylogeny created from the diversity representation genome set. The percentage of species members with GCFs was determined using lsaBGC-AutoExpansion.py results for all ~15K *Staphylococcus* genomes and was shown in $\log_{10}$ scale.

### Ancestral inference of gene cluster family carriage

We performed ancestral inference of GCF carriage to predict vertical descent of GCFs within the skin-associated phylogenetic clades of *Staphylococcus* and *Corynebacterium* (Figure S4ab)[38]. GCF carriage across genomes was encoded as a binary trait and AncestralGeneRator[101] was used to perform ancestral state reconstruction to infer carriage for inner-nodes of the *Corynebacterium* and *Staphylococcus* ribosomal phylogenies. Briefly, PAUP (v4b)[102] was used to perform maximum parsimony with the ACCTRAN algorithm, setting gain and loss costs to 10 and 5, respectively. Afterwards, result files from AncestralGeneRator were further processed and were visualized using iTol[98]. This analysis revealed that four GCFs were ancestral to the *S. epidermidis/aureus* clade (predicted siderophore, terpene/T3PKS, cyclic lactone autoinducer, and hserlactone) and five GCFs were ancestral to the *C. tuberculostearicum* species complex (predicted siderophore, terpene, T1PKS, NAPAA, and NAPAA/betalactone).

### Inspection of *Staphylococcus* and *Corynebacterium* GCFs with high Beta-RD values

*Investigation into High Beta-RD of lugdunin encoding NRPS GCF-46 in* Staphylococcus: Among staphylococcal GCFs, GCF-46 depicted the highest posterior Beta-RD distribution (Figure S4c). This GCF encodes for the lugdunin NRPS, which was determined to be an antibiotic produced by *S. lugdunensis* which is active against *S. aureus*[26]. The ubiquity of this GCF being regarded as present in multiple staphylococcal species by lsaBGC-AutoExpansion.py was unexpected; however, further investigation revealed a rare instance of a small insertion-sequence element, which is ubiquitous across *Staphylococcus*, within the protocore region of the lugdunin BGC, thus resulting in lsaBGC-AutoExpansion.py over-classifying the presence of the GCF as it assumes the genes within the element are highly BGC associated. Although we suspect such cases to be rare, we have since introduced more stringent requirements for defining homolog groups as part of the protocore of BGCs predicted by antiSMASH in more recent releases of *lsa*BGC.

*Investigation into* Corynebacterium *GCF with the second highest Beta-RD value*: In addition to *Corynebacterium* NRPS encoding GCF-50, the NRPS encoding GCF-9 exhibited a similarly high posterior Beta-RD distribution (Figure S4d). GCF-9 was found to also be present in multiple species on the skin and was most prevalent in the well known skin pathogen *C. diphtheria*[103]. This GCF exhibited a similar scenario to GCF-50, in which it featured a highly conserved region with NRP synthase(s) flanked by MGEs in addition to a nearby phage and toxin/antitoxin system. The NRP synthase (HG0002384) within GCF-9 could be found in multiple copies within certain genomes.

### BGC codon assimilation relative to background genomic contexts

Codon usage of individual genes can be compared to the general distribution of all coding genes across a genome to infer potential genomic islands[104]. Depending on when the horizontal gene transfer event occurred, genes horizontally inherited could potentially assimilate for codon usage within the genome through vertical evolution. We developed a standalone script, compareBGCtoGenomeCodonUsage.py, for calculating codon usage differences between coding genes within BGCs of a particular GCF and genes observed outside of the BGC context. Briefly, this program calculates the frequency of codons for coding genes within BGCs and then compares those to the frequency of codons observed in the complementary set of coding genes across the genome. For comparison of the two codon frequency distributions, the program computes both the cosine distance and Spearman's correlation rho. Genes of a length not divisible by three are ignored.

### Understanding GCF conservation and composition through evolutionary and population genetic statistics

A core program of the *lsa*BGC suite is lsaBGC-PopGene.py which generates a table report of conservation and evolutionary statistics for each homolog group found within a specific GCF. To calculate certain homolog group statistics within this report, lsaBGC-PopGene.py begins by constructing protein alignments[83] for each homolog group and translating those to codon-based alignments using PAL2NAL[84]. Codon-based alignments are additionally used to visualize conservation and domain structure for homolog groups and downstream in lsaBGC-

DiscoVary.py. Three of the evolutionary statistics which codon alignments are used for computing are: (i) homolog-group specific variants of the Beta-Relative Divergence statistic, (ii) the rate of non-synonymous mutations relative to the rate of synonymous mutations (dN/dS), and (iii) Tajima's D statistic, which can be used to detect signatures of sweeping vs. balancing selection. lsaBGC-PopGene.py also allows users to specify population designations for each isolate with the focal GCF and computes additional statistics per homolog group pertaining to this information.

Assuming users provide genome-wide similarity estimates, either ANI or AAI, to lsaBGC-PopGene.py, it will calculate how pairwise homolog group similarities in codon alignments or protein alignments compare to such genome-wide expectations. This is reported as the median Beta-RelativeDivergence (analogous to the BGC-wide metric described in the *Materials and Methods*) across all pairs of genomes with the homolog group found in the focal GCF.

Tajima's D is a simple statistic where values below -2 typically indicate conservation or sweeping selection and a low ratio of rare minor alleles to high-frequency minor alleles while values above 2 indicate balancing selection. To calculate Tajima's D we modified a previous implementation[105] into a function within *lsa*BGC. Namely, we adjusted the calculation of Tajima's D to better reflect the mathematical derivation described by Simonsen, Churchill and Aquadro 1995[106]. Additionally, we only considered sites as segregating or pairwise differences if both sequences being compared had a valid allele at the position within the alignments (alignment sites with gaps were ignored). As an alternate assessment of sequence variation to Tajima's D, we also report the proportion of sites across homolog group multiple sequence alignments where multiple alleles exist (major allele < 98%) and the proportion of sites where the major allele is non-dominant (<75%).

For the comprehensive analysis of Tajima's D reported in this study, we only regarded homolog groups found in four or more genomes and which were strictly single-copy within GCFs (Table S7). We comprehensively investigated Tajima's D values for common homolog groups across all four genera in our study and found that it reflected expectations of a null model, with the median intra-species Tajima's D being centered at a value of roughly 0 (Figure S5a). When calculating Tajima's D in aggregate across species, a large fraction of homolog groups (17.3%) were found to exhibit high values ($\geq$2) suggesting balancing selection; however, most of these homolog groups appear to be ancestrally acquired and likely diverged in sequence simply due to speciation rather than targeted selective pressures (Figure S5b). To better discern true instances of balancing selection (Tajima's D $\geq$2) and conservation or sweeping selection (Tajima's D $\leq$-2) within species, we compared the aggregate genus-wide Tajima's D with the maximum and minimum value of Tajima's D observed for a single species (Figure S5b). Species-specific Tajima's D values for *S. aureus* homolog groups in GCFs highlighted the staphyloxanthin encoding GCF-3 as featuring the greatest number of highly conserved homolog groups (Tajima's D $\leq$ -2) (Figure S5c).

The rate of non-synonymous mutations relative to the rate of synonymous mutations is a classical statistic used to infer the effect of positive versus negative selection. If the rate for non-synonymous mutations is higher than the rate for synonymous mutations across different instances of a gene (dN/dS > 1), it could suggest positive selection; whereas, if the reverse is true (dN/dS < 1), it could suggest negative or purifying selection. We used Biopython's codonalign.codonseq module[107] to calculate dN/dS using the method described by Nei and Gojobori 1986[108]. To more robustly calculate the statistic, we replace singleton codon instances (those observed in only one in one BGC) with gaps. Additionally, to ensure we avoid excessive computation when the number of sequences with a homolog group is large, we have implemented an empirical but non-exhaustive framework in which we calculate the median dN/dS between 1000 randomly selected pairs of samples. This random sampling and calculation of dN/dS is performed for 20 iterations and the median of dN/dS estimates across iterations, along with the absolute median deviation to assess robustness, is reported. Because we are actively seeking to improve our calculations of dN/dS in *lsa*BGC and Biopython's codonalign module is under development currently, we do not report results related to dN/dS in this study.

**Inference of consensus order and directionality of homolog groups for a GCF**

We developed an algorithm to infer the consensus order of homolog groups relative to each other in the GCF, as well as their relative consensus directionality (sense vs. antisense). This algorithm works by first computing how many times a homolog group proceeds another homolog group across BGC instances belonging to a GCF. To gather this information, a single BGC instance, with the most homolog groups, is selected as the reference and used for configuring the general direction of the remaining GCF instances, deciding whether to flip them to better align with the order of the reference. The gene order information for each BGC, encoded as a dictionary, is then used to construct a primary path, starting from the homolog group most often found at one edge of a GCF to the homolog group most often found on the opposite edge. Afterwards, homolog groups which were not featured in this consensus path, potentially because they are infrequently found and not core to the GCF, are attempted to be placed in their most appropriate locations. The core ordering path will be structured primarily by more prevalent homolog groups. When looking at the lsaBGC-PopGene.py report table sorted by the consensus ordering of the homolog groups, it is thus important to consider the proportion of samples with the GCF which have a particular homolog group. The consensus directionality of homolog groups is simply based on whether most instances of it are forward or reverse relative to a reference gene from some chosen representative BGC.

**Metagenomic mining for BGCs, homolog groups, and novel SNVs**

Metagenomic methods to explore microdiversity within microbiomes are continually advancing[109,110]. We developed lsaBGC-DiscoVary.py to explore the micro-diversity of BGCs beyond the limited set of single-isolate genomes available for a lineage or taxa of interest and allow users to identify BGC genes and base-resolution novel SNVs within metagenomic datasets (Figure S2). Briefly, lsaBGC-DiscoVary.py serves two roles:

1. Assessing the presence of GCFs, individual homolog groups and if requested, phasing them (uses a custom approach leveraging DESMAN[111]), and
2. If users construct a comprehensive database of homolog groups alleles observed across all available assemblies for a taxonomy, lsaBGC-DiscoVary.py can also be used to identify putatively novel SNVs never previously observed at a particular site in a homolog group.

*Selection of Representative Alleles for Reference Database:* lsaBGC-DiscoVary.py begins by identifying representative allelic sequences for each homolog group in the focal GCF from the codon alignments listing file provided. To select representative alleles, it parses each homolog group's MSA FASTA file and determines the number of differences between pairs of sequences. Sequences are deemed to be members of the same allelic cluster if they exhibit $\geq$99% identity of the shorter sequence and they differ at less than 10 sites from each other. Afterwards, pairs of such similar sequences are joined into larger clusters through single-linkage clustering and a representative sequence is chosen based on the minimal summed differences to the other sequences in the allelic cluster (e.g. the centroid). A Bowtie2[112] reference database is finally constructed from the representative sequences.

_Read Alignment to Reference Database and Alignment Parsing_: Processed sequencing reads are next aligned against the database of representative sequences for homolog group alleles using Bowtie2 in "--very-sensitive-local" mode[112]. Because our database is currently based on individual genes (which are rather short), we found that aligning paired-end reads individually (as unpaired reads), increased sensitivity and did not significantly compromise specificity. This is similar to the approach for mapping used in MetaMLST[113]. After alignment, sorting and indexing of BAM files is performed using samtools[114]. Alignments for each homolog group are then processed and investigated using the pysam library in Python[115]. An allele of a homolog group is considered potentially present if 90% of its sites are covered by at least one read, accounting for whether a read has lower than 30 base-quality and whether a site is a skipped region in the reference or corresponds to a deletion. Each alignment of a read to an allelic representative of a homolog group is then assessed as to whether it exhibits:

1. Alignment to multiple allelic representatives of the homolog group. Only alignments with the top/maximum alignment score of the read to any of the allelic representatives of the homolog group will be considered. This allows partitioning a read into one or more allelic representatives.

2. The read displays at least 95% identity to the reference allele sequence within the core alignment, where the core alignment is defined as the part of the alignment in between the first and last positions where reference and query sequences both have valid nucleotides (even if non-matching). If the core alignment length is >= 100 bp, 95% identity is required, while if it is shorter, >= 60 bp, 99% identity is required.

3. The total indel length within the core alignment is < 5 bp. This allows for some leniency around small deletions and insertions, so as to not discard otherwise high-quality alignments.

If an alignment meets the above criteria, it is next assessed at each position for high base quality (>= 30 PHRED). If so, then the reference allele site is considered to be covered and the base of the read/query is noted. Further, because the reference allelic sequences were already aligned to each other and provided as the codon alignment inputs, we can translate the position of a site on the reference allele to a position in the codon multiple sequence alignment and ultimately gain a universal tally of base counts at particular sites of a homolog group. The synchronization of reference gene positions to codon alignment positions is a key feature of lsaBGC-DiscoVary.py which enables it to identify novel SNVs.

*Final Assessment of Homolog Group Presence*: As is often cautioned in gene-based metagenomic analysis, faulty alignment of reads belonging to the lineage of interest or from other taxa can lead to a misinterpretation of enzyme presence or association with the lineage of interest. To further filter out faulty alignments, we parse the codon alignments of each homolog group, featuring reference and representative allelic sequence, and mark regions along it which are particularly "gappy", >10% of sequences have no allelic residue, as troublesome, including +/- 50bp around the start and end of each region. These regions were observed to present problems as default gap opening and extension costs in the aligning algorithm can occasionally lead to faulty alignments. Based on a similar logic that alignment and alignment scores could behave irrationally when only part of reads should properly align to a reference, we also deem the first and last 50 bp of a MSA as "troublesome".

Additionally, we self-align the full predicted proteome of the genomes used to establish representative alleles for homolog groups with DIAMOND[66] to identify regions along the homolog group MSAs which are similar at high-identity to potential paralogs. The criteria for defining these regions matches our criteria for mapping reads to allelic sequences. Thus, alignments which are ≥20 residues long and exhibit ≥99% amino acid identity or ≥33 residues long and exhibit ≥90% amino acid identity for ≥5% of the initial sample set are marked as "troublesome" for accurate alignment of reads.

For each sequencing read set, each homolog group is next more thoroughly assessed for carriage in the context of the full BGC. Up to this stage, the criteria for consideration of a BGC homolog group as present is pretty lenient and simply requires 1X coverage at 90% of sites for one of the representative alleles of the homolog group. Here, we further refine this criteria to require 1X coverage at ≥90% of sites in the codon alignment of the homolog group which are deemed as non-troublesome to align. For homolog groups which meet this requirement, the median depth of the middle 80% of positions is computed. The median of these homolog group specific median depths is then calculated along with the median absolute deviance. Because these homolog groups are expected to be co-located together in a BGC, potential variability in sequencing coverage across the genome, associated with active replication[116], should not result in a difference in coverage between homolog groups of the BGC. Based on this assumption, we next aim to identify and disregard homolog groups which are outliers in terms of their coverage (>2 median absolute deviances from the median of 80% trimmed median depths). These homolog groups will be difficult to gauge from raw metagenomics sequencing data as they will likely either lack enough coverage for resolved allele typing or have too much coverage and correspond to potentially multi-copy or common enzymes where undesired reads (from outside the focal BGC or lineage) are being aligned to the homolog group. Homolog groups are also disregarded if one of their predicted products contains mobile genetic element MGE suggestive keywords 'integrase' or 'transp'. In the most recent release of *lsa*BGC, we have updated the second keyword to ‘transpos’ to still allow for novel SNV detection on transporters.

If there are at least 5 homolog groups which are deemed present and not filtered by the above criteria, then BGC presence is assessed as a whole based on whether ≥70% of the core homolog groups are present (where the core homolog groups are those found in all BGCs from the initial *lsa*BGC processing/clustering analysis - e.g. the BGCs from high-quality genomes) or if just a single GCF specific homolog group is observed. Homolog groups are also disregarded if ≥5% of the initial set of the high-quality genomes used for initial lsaBGC analysis featured multiple copies of the homolog group (paralogs were common). Finally, a report file will be generated featuring only homolog groups deemed present (not filtered by above criteria) for samples which are regarded as featuring the BGC.

Note for investigations of cutimycin in *C. acnes*, we manually specified the core homolog groups involved in the thiopeptides biosynthesis[27] to increase sensitivity through an optional setting in lsaBGC-DiscoVary.py.

*Consensus allele determination, allelic phasing, and phylogenetic visualization*: For the set of retained homolog groups deemed as present within a sequencing readset for the focal GCF, the proportion of positions which are heterozygous are next computed and used to determine whether allelic phasing is needed or appropriate (current default $\geq$5% of sites along present homolog groups in the sequencing dataset need to be heterozygous to turn on phasing mode). If phasing mode is initiated and specified by users, DESMAN[111] is used to first determine the most likely number of strains and then phase the distinct haplotypes. If phasing mode is not initiated, then the consensus / majority-rule allele is selected for each site along the homolog group. Critically, the output from emission of the consensus sequence or multiple phased alleles for the homolog groups is not an independent sequence but rather an allele call for the sequencing / metagenomic read set at each position in the homolog group codon alignment. This allows us to avoid inferring faulty frameshifts in our sample-specific sequence(s) of homolog groups and allows direct incorporation into the codon alignment to build phylogenetic views. Additionally, we require that each base emitted / inferred has a minimum depth (current default is 5) and for the total depth at each position to be within a reasonable range of the median depth observed across all homolog groups regarded as present. For each inferred sequence, if an in-frame stop codon is observed, then downstream sites are automatically emitted as gaps.

The sequences inferred for each homolog group are brought into the context of the full codon alignments of reference alleles, after which filtering is performed to remove sequences which have gaps at more than 25% of non-"troublesome" sites of the alignments. Of the sequences retained after this filtering, individual sites are filtered to retain only those in which at most 10% of sequences have gaps / ambiguity. The resulting FASTA files are input into FastTree2[99] to infer a quick homolog group specific phylogeny which is then visualized in R and used to display the similarity of newly identified alleles / sequences to known / reference sequences.

*Identification of novel SNVs:* The cornerstone feature of lsaBGC-DiscoVary.py is its ability to search metagenomic / raw-read datasets and then assess whether they possess any potential novel SNVs not previously observed in the comprehensive set of known alleles gathered from all available assemblies for a taxa. Currently, only SNVs which are supported by at least 5 reads and not located in sites along the codon-alignment marked as "troublesome" are reported. Additionally, SNVs are not reported after the first in-frame stop codon observed from preliminary scanning. A comprehensive report of putatively novel SNVs is produced for all sequencing samples and homolog groups. A subset of reads which are supportive of putatively novel SNVs (last column in the report) are written in gzipped FASTQ format for each sequencing / metagenomic sample. This allows users to quickly taxonomically profile and assess that reads in fact belong to the lineage of interest (see subsection below).

**Benchmarking *lsa*BGC-DiscoVary against assembly-based novel variant detection using *M. luteus* single-isolate sequencing readsets**

The correspondence for novel single nucleotide variants (SNVs) reported by lsaBGC-DiscoVary.py was assessed using whole-genome sequencing readsets for single isolates and compared to SNV identification based on an assembly based approach (Figure S8). lsaBGC-AutoProcess.py was run on all complete instances of *M. luteus* genomes present in GTDB to identify BGCs which were then clustered into 9 GCFs using lsaBGC-Cluster.py. Afterwards, lsaBGC-AutoExpansion.py was used to search all remaining, draft-quality, genomes from the species for homologous instances of the 9 GCFs identified and lsaBGC-PopGene.py was run to generate codon-based alignments for homolog groups associated with each GCF. We then used the sequencing reads from 132 *M. luteus* isolates sequenced by our lab, and absent in the current GTDB release, to run lsaBGC-DiscoVary.py and identify novel variants not previously observed in available genomes for the species. Because we had constructed draft assemblies for these same samples, we also identified GCF and homolog group instances in each assembly using lsaBGC-AutoExpansion.py. MAFFT[83] was then used to incorporate the homolog group sequences identified in the draft assemblies into the codon alignments used for lsaBGC-DiscoVary.py analysis and the expanded codon alignments were subsequently parsed and assessed for novel SNVs. We found a high concordance between the lsaBGC-DiscoVary.py and assembly-based approaches for novel SNV detection, with 1788 of 1798 (99.5%) novel SNVs reported by DiscoVary also being found by the assembly based approach and 1788 of 1858 (96.3%) novel SNVs found by the assembly based approach also being reported by lsaBGC-DiscoVary (Table S11). Of the 69 novel SNVs only found by the assembly based approach, 30 were identified by lsaBGC-DiscoVary.py but not reported due to alleles exhibiting more or less coverage than expected given the median coverage of the BGC. Among the remaining 39 novel SNVs, manual examination of a common SNV (homolog group OG0001039 - position 65 in the codon alignments; found in 15 samples), revealed that it involved a cytosine allele within a lengthy stretch of 21 C/Gs, which could lead to less confident read alignment in lsaBGC-DiscoVary.py. Of the nine novel SNVs found by lsaBGC-DiscoVary, six correspond to minor alleles, explaining why they were not represented in sample assemblies.

#### ***lsa*BGC-DiscoVary leads to the identification of truncated uracil DNA glycosylase in clade III *C. acnes***

To demonstrate *lsa*BGC-DiscoVary's application on metagenomic datasets, its primary intended usage, we chose to investigate the microdiversity of BGCs from *C. acnes* within healthy skin metagenomic sequencing datasets generated by our lab[37]. *C. acnes* are the most prominent species on the skin and have recently been shown to encode for a BGC producing the anti-Staphylococcal thiopeptide cutimycin[27]. We predicted GCFs from complete genomes available for the species and subsequently identified additional instances in draft genomes. We found that relative to other representative species from the four genera investigated in our study, *C. acnes* BGCs were largely clade specific (Figure S9). These clade specific differences in BGC content could potentially be due to bottlenecking of *C. acnes* populations across skin microenvironments, such as pores, as was recently reported[117]. Of the six GCFs from *C. acnes*, the one encoding for cutimycin demonstrated high codon usage dissimilarity with the background genome, suggestive of horizontal acquisition (Figure 4a). This GCF is encoded by two of the three distinct phylogenetic clades of *C. acnes*, clades I (primarily subclade IB) and III (Figure 4b). Similar codon usage assimilation patterns for the BGC between the two clades along with a common flanking genetic context, further suggested cutimycin was ancestrally acquired by *C. acnes* and likely lost in clade II and other subclades of clade I (Figure S10a). Application of *lsa*BGC-DiscoVary to search for the presence of the cutimycin GCF in 270 metagenomes across eight body sites for 34 participants revealed that 35.9% of metagenomes encoded for the gene cluster (Table S12). Reference sequences for the core biosynthetic genes were assessed for nearly-fixed sites differentiating genetic alleles between the clade I and clade III *C. acnes* (see subsection '*Identification of* Cutibacterium *strains within metagenomes and …*'). Using output from *lsa*BGC-DiscoVary, we found that presence of clade specific alleles in genes aligned with predictions of clade I and clade III presence in metagenomes using StrainGST (Figure 4c-d, S10bc)[38].

Putatively novel SNVs reported by *lsa*BGC-DiscoVary were further scrutinized through exhaustive taxonomic and genomic localization assessment of sequencing reads supporting SNV presence in metagenomes (Figure S2c). Novel variants in the core cutimycin BGCs were most commonly found in the gene encoding for the N-terminal domain of the lantibiotic dehydratase (Figure 4e). Of the 59 instances of novel SNVs detected in the lantibiotic dehydratase gene, 36 (61%) corresponded to a single participant in our skin microbiome sampling survey, S012. The majority of these 36 novel variant instances were repeatedly observed at multiple body sites and corresponded to nine unique SNVs rarely observed in other participants; aside from participant S003, which featured five of the nine (55.6%) novel SNV sites at a single body site, their occiput (Figure 4f). Participant S003 worked in the same building and floor as participant S012. Further examination revealed that most of the corresponding novel variants from S012 involved C to T deaminations and were found on reference gene alleles from clade III rather than clade I *C. acnes* (Figure 4g). In accordance with this observation, we found that the uracil DNA glycosylase gene, responsible for repairing such deaminations, was predicted to be truncated for clade III *C. acnes* alone (Figure 4h-k; *Materials and Methods*). This suggests that the variation identified by *lsa*BGC-DiscoVary is legitimate and not due to unaccounted sequencing artifacts.

*Identification of* Cutibacterium *strains within metagenomes and determination of near-fixed sites differentiating cutimycin biosynthesis alleles between clade I and clade III* C. acnes: We used the StrainGE suite[38] to profile *Cutibacterium* strains within metagenomic samples. Recommendations for running the suite in the documentation were largely followed, whereby we used mainly complete genomes as references and performing dereplication; however, we also manually included *C. acnes* Asn12 as a distinct reference to represent clade III *C. acnes*, for which no complete genome exists.

To differentiate alleles of the seven cutimycin core biosynthesis homolog groups belonging to clade I and clade III *C. acnes*, we identified 170 near-fixed differences, where at least 95% of genomes in one clade featured a particular allele which was different or absent in the other clade.

*Assessing cytosine deamination rates between clade I and clade III* C. acnes*:* A majority of the novel SNVs discovered for the cutimycin BGC in *C. acnes* by *lsa*BGC-DiscoVary from participant S012's skin metagenomes were in the N-terminal subunit of the lantibiotic dehydratase and corresponded to C -> T deaminations. These represented instances where the novel SNV detected in metagenomes was a thymine in place of an expected cytosine, as observed on the reference allele for which the SNV-supporting reads mapped to the best (Figure 4g). Based on observations that the majority of these deamination SNVs corresponded to reference alleles belonging to clade III *C. acnes*, we investigated whether such C -> T deaminations were common amongst clade III *C. acnes*. To assess this we first performed whole-genome alignment using Parsnp[118] with all clade I and clade III *C. acnes* found to carry the cutimycin BGC, along with an outgroup genome *C. namnetense* T34998, which represents the closest neighbor of *C. acnes*. The GCF_000008345 genomic assembly for *C. acnes* KPA171202 was used as the reference for the alignment. Based on the resulting core whole-genome alignment, we first looked at the proportion of total pairwise differences which corresponded to cytosine and thymine differences within and between clade I and clade III *C. acnes* genomes (Figure 4i). Amongst pairs of genomes separated by 500 or more differences along the core alignment, pairs of clade III *C. acnes* genomes were found to have a greater proportion of differences owing to cytosine and thymine transitions. Next, to more directly assess whether clade III *C. acnes* experienced elevated rates of C to T deaminations compared to clade I *C. acnes*, we treated alleles observed in the outgroup *C. namnetense* genome as ancestral. For each site where a cytosine was observed in *C. namnetense*, 100 simulations were performed, where for each simulation six alleles were randomly drawn from clade I (n=43) and clade III (n=6) genomes without replacement at the site in the whole-genome alignment and the frequency of thymine alleles was aggregated separately for the two clades of *C. acnes*. Across all sites and simulations, this resulted in a sum of 2,057,933.33 for clade III *C. acnes* as compared to 1,750,291.83 in clade I *C. acnes* (Figure 4j). Note, we excluded the GCF_004136215 assembly for *C. acnes* KPA171202 and used only the GCF_000008345 assembly for the strain for these analyses to avoid double-counting in metrics.

To understand why C to T deanimations were more prevalent in clade III *C. acnes*, we looked at whether they were lacking or featuring an altered form of uracil DNA glycosylase, the enzyme responsible for correcting such mutations. We found that the canonical variant of the enzyme from clade I *C. acnes* strain 6609 (278 aa) was present at ≥50% identity and ≥70% query coverage in all predicted-proteomes of clade I and clade II *C. acnes*, but missing at these thresholds in all clade III *C. acnes* through BLASTp analysis (Figure 4k)[90]. Further investigation revealed that the open-reading frame in *C. acnes* clade III genomes appears to feature a stop-codon and the reference protein matches two adjacent predicted proteins one with ~94% identity and ~64% query coverage and the other with ~99% identity and ~29% query coverage. The high-level of conservation might suggest that the enzyme might potentially still be produced, though potentially through stop codon readthrough at lower throughput[119]. Additionally, four of the six non-acnes *Cutibacterium* genomes which were not deemed to possess the clade I *C. acnes* uracil DNA glycosylase protein at the thresholds mentioned were *C. granulosum.* These four *C. granulosum* genomes did not feature any considerable homologs to the enzyme in contrast to the clade III *C. acnes* genomes.

**Determination of whether novel SNVs detected for BGCs of the *C. tuberculostearicum* species complex would be identified in metagenomic assemblies:**

Because the *C. tuberculostearicum* species complex can be found at low abundance at certain body sites within individuals, we wanted to determine whether the 34,545 novel SNVs found in BGC contexts of the species complex, after filtering SNVs where reads mostly map with higher scores to other regions of *Corynebacterium* genomes (as described in the previous subsection), would also be detected through metagenomic assembly. MEGAHIT[49] assembly was performed individually for all metagenomic readsets using default options and filtered to only retain 2 kb or longer contigs. Sample specific metagenomic assemblies were then combined into one consolidated FASTA file and used to construct a Bowtie 2 database[112]. No pooling of samples for metagenomic assembly or subsequent dereplication was performed to ensure strain-specific variability is retained. After assessing the presence of putative novel SNVs and filtering those where supportive reads are mostly mapped to alternate regions in *Corynebacterium* genomes at higher scores (as described in the *Materials and Methods*), we performed similar investigations through Bowtie 2 mapping of reads supporting novel SNV presence to the concatenated database of contigs from individual metagenomes. Exact mappings of SNV-supporting reads to contigs were identified and used to assess whether novel SNVs are represented in metagenomic assemblies. Of the 34,545 novel SNVs deemed as reliable after mapping to the comprehensive database of *Corynebacterium* genomes, we found that 22,886 would also be identified from metagenomic assembly. Thus, 11,659 SNVs detected by lsaBGC-DiscoVary.py would be missed by metagenomic assembly and are not represented in contigs of considerable length. This is expected to be an underestimate as a concatenated database of metagenomic contigs from multiple samples was used. We additionally tested mapping SNV supporting reads to only a subset of the metagenomic contigs which we deemed as belonging to *Corynebacterium.* These contigs were identified by BLASTn analysis[90] of the metagenomic contigs to the comprehensive *Corynebacterium* genomics database used to filter putative novel SNVs in the previous subsection. Contigs with HSPs with query coverage greater than 25% and sequence identity greater than 85% or query coverage greater than 70% and sequence identity greater than 70% to one of the known *Corynebacterium* genomes were classified as *Corynebacterium*. Of the 556,030 total concatenated metagenomic contigs, 91,713 were classified as *Corynebacterium*. However, using this subset of contigs instead of the full metagenomic assembly database resulted in a minor increase in the number of SNVs which would be undetected by metagenomic assembly (11,763 instead of 11,659; an increase of 0.9%), suggesting that reads supporting novel SNVs detected by lsaBGC-DiscoVary.py were >99% *Corynebacterium.*

**Additional details on the relation of predicted synonymous to non-synonymous rates of novel SNVs with enzymatic ancestral conservation percentiles for BGCs from the *C. tuberculostearicum* species complex.**

Of the 2,343 unique novel SNVs identified in protocore or protocore adjacent homolog groups of BGCs from the *C. tuberculostearicum* species complex, 2,244 were found in 34 homolog groups with conservation information calculated from alignments featuring distantly related homologs from diverse bacteria. Novel SNVs found in the top five percentile of conserved sites for a homolog group were approximately 10X more likely to correspond to a synonymous change as opposed to a non-synonymous change (Figure 5b). In contrast, the bottom 20% of conserved sites (the least conserved sites), were only 1.7X as likely to represent non-synonymous substitutions as synonymous substitutions. Further, the number of novel SNVs, either synonymous or non-synonymous observed at conserved sites, amongst the top 20% of conserved sites was 414, and lower than the number of novel SNVs found in the 20-40 (n=459), 40-60 (n=490), and 60-80 (n=493). Only 388 novel SNVs were found in the 80-100 percentile ranges of conserved sites and we suspect that this decrease is due to read alignments to such regions becoming more challenging at the thresholds required by lsaBGC-DiscoVary.py.

**Additional References**


89. Smirnov, V. Recursive MAGUS: Scalable and accurate multiple sequence alignment. *PLoS Comput. Biol.* **17**, e1008950 (2021).
90. Camacho, C. *et al.* BLAST+: architecture and applications. *BMC Bioinformatics* **10**, 421 (2009).
91. van Dongen, S. & Abreu-Goodger, C. Using MCL to extract clusters from networks. *Methods Mol. Biol.* **804**, 281–295 (2012).
92. Melnyk, R. A., Hossain, S. S. & Haney, C. H. Convergent gain and loss of genomic islands drive lifestyle changes in plant-associated Pseudomonas. *ISME J.* **13**, 1575–1588 (2019).
93. Cantalapiedra, C. P., Hernández-Plaza, A., Letunic, I., Bork, P. & Huerta-Cepas, J. eggNOG-mapper v2: Functional Annotation, Orthology Assignments, and Domain Prediction at the Metagenomic Scale. *Mol. Biol. Evol.* **38**, 5825–5829 (2021).
94. Eddy, S. R. Accelerated Profile HMM Searches. *PLoS Comput. Biol.* **7**, e1002195 (2011).
95. Schreiber, J. Pomegranate: fast and flexible probabilistic modeling in python. *J. Mach. Learn. Res.* (2017).
96. Wickham, H. *ggplot2: Elegant Graphics for Data Analysis*. (Springer, New York, NY, 2009).
97. Yu, G., Smith, D. K., Zhu, H., Guan, Y. & Lam, T. T.-Y. Ggtree : An r package for visualization and annotation of phylogenetic trees with their covariates and other associated data. *Methods Ecol. Evol.* **8**, 28–36 (2017).
98. Letunic, I. & Bork, P. Interactive Tree Of Life (iTOL) v4: recent updates and new developments. *Nucleic Acids Res.* **47**, W256–W259 (2019).
99. Price, M. N., Dehal, P. S. & Arkin, A. P. FastTree 2--approximately maximum-likelihood trees for large alignments. *PLoS One* **5**, e9490 (2010).

100. Gilchrist, C. L. M. & Chooi, Y.-H. clinker & clustermap.js: automatic generation of gene cluster comparison figures. *Bioinformatics* **37**, 2473–2475 (2021).

101. *AncestralGeneRator: This repo contains two programs for studying evolutionary gene flux across a phylogeny: GeneFluxAnalysis.py and CorStrictor.py*. (Github).

102. Swofford, D. L. PAUP: phylogenetic analysis using parsimony. *Mac Version 3. 1. 1.(Computer program and manual).* (1993).

103. Hadfield, T. L., McEvoy, P., Polotsky, Y., Tzinserling, V. A. & Yakovlev, A. A. The pathology of diphtheria. *J. Infect. Dis.* **181 Suppl 1**, S116-20 (2000).

104. Waack, S. *et al.* Score-based prediction of genomic islands in prokaryotic genomes using hidden Markov models. *BMC Bioinformatics* **7**, 142 (2006).

105. Whalley, T. *LICENSE at master · WhalleyT/tajima*. (Github).

106. Simonsen, K. L., Churchill, G. A. & Aquadro, C. F. Properties of statistical tests of neutrality for DNA polymorphism data. *Genetics* **141**, 413–429 (1995).

107. Cock, P. J. A. *et al.* Biopython: freely available Python tools for computational molecular biology and bioinformatics. *Bioinformatics* **25**, 1422–1423 (2009).

108. Nei, M. & Gojobori, T. Simple methods for estimating the numbers of synonymous and nonsynonymous nucleotide substitutions. *Mol. Biol. Evol.* **3**, 418–426 (1986).

109. Olm, M. R. *et al.* inStrain profiles population microdiversity from metagenomic data and sensitively detects shared microbial strains. *Nat. Biotechnol.* (2021) doi:10.1038/s41587-020-00797-0.

110. Gregory, A. C. *et al.* MetaPop: a pipeline for macro- and microdiversity analyses and visualization of microbial and viral metagenome-derived populations. *Microbiome* **10**, 49 (2022).

111. Quince, C. *et al.* DESMAN: a new tool for de novo extraction of strains from metagenomes. *Genome Biol.* **18**, 1–22 (2017).

112. Langmead, B. & Salzberg, S. L. Fast gapped-read alignment with Bowtie 2. *Nat. Methods* **9**, 357–359 (2012).

113. Zolfo, M., Tett, A., Jousson, O., Donati, C. & Segata, N. MetaMLST: multi-locus strain-level bacterial typing from metagenomic samples. *Nucleic Acids Res.* **45**, e7 (2017).

114. Li, H. *et al.* The Sequence Alignment/Map format and SAMtools. *Bioinformatics* **25**, 2078–2079 (2009).

115. Gilman, P. *et al. PySAM (Python Wrapper for System Advisor Model "SAM")*. https://www.osti.gov/servlets/purl/1559931 (2019) doi:10.11578/dc.20190903.1.

116. Bremer, H. & Churchward, G. An examination of the Cooper-Helmstetter theory of DNA replication in bacteria and its underlying assumptions. *J. Theor. Biol.* **69**, 645–654 (1977).

117. Conwill, A. *et al.* Anatomy promotes neutral coexistence of strains in the human skin microbiome. *Cell Host Microbe* (2022) doi:10.1016/j.chom.2021.12.007.

118. Treangen, T. J., Ondov, B. D., Koren, S. & Phillippy, A. M. The Harvest suite for rapid core-genome alignment and visualization of thousands of intraspecific microbial genomes. *Genome Biol.* **15**, 524 (2014).

119. Eggertsson, G. & Söll, D. Transfer ribonucleic acid-mediated suppression of termination codons in Escherichia coli. *Microbiol. Rev.* **52**, 354–374 (1988).
